# Purging of deleterious mutations during domestication in the predominant selfing crop soybean

**DOI:** 10.1101/2020.03.12.989830

**Authors:** Myung-Shin Kim, Roberto Lozano, Ji Hong Kim, Dong Nyuk Bae, Sang-Tae Kim, Jung-Ho Park, Man Soo Choi, Jaehyun Kim, Hyun Choong Ok, Soo-Kwon Park, Michael A. Gore, Jung-Kyung Moon, Soon-Chun Jeong

## Abstract

As a predominant plant protein and oil source for both food and feed, soybean is unique in that both domesticated and wild types are predominantly selfing. Here we present a genome-wide variation map of 781 soybean accessions that include 418 domesticated (*Glycine max*) and 345 wild (*Glycine soja*) accessions and 18 of their natural hybrids. We identified 10.5 million single nucleotide polymorphisms and 5.7 million small indels that contribute to within- and between-population variations. We describe improved detection of domestication-selective sweeps and drastic reduction of overall deleterious alleles in domesticated soybean relative to wild soybean in contrast to the cost of domestication hypothesis. This resource enables the marker density of existing data sets to be increased to improve the resolution of association studies.

## Introduction

Soybean (*Glycine max* [L.] Merr.) is an important crop species. It is a major source of protein and oil. Cultivated soybean (*G. max*) has been domesticated as early as 7,000–9,000 years ago from wild soybean (*Glycine soja* Sieb. & Zucc.) with distribution in East Asia^1,2^. The cultivation of soybean has been historically confined to East Asia and only recently expanded to North America, South America, and India, positioning it as one of the top crops in terms of growing area worldwide^3^. Both domesticated and wild soybean types are predominantly selfing^4^. The accumulation of recombination events in such selfing crops may result in a rapid fixation of both beneficial and deleterious mutations. Deleterious mutations are considered to be the genetic basis of inbreeding depression and heterosis in other major crops including maize and cassava that have outcrossing mating systems^5^. Thus, understanding of genome-wide patterns of historical deleterious allele fixation in soybean provides insight into breeding strategy of soybean itself as well as other major crops. After the release of the draft soybean genome sequence^6^, efforts to map soybean genetic variation by single nucleotide polymorphism (SNP) array genotyping^2,7^ and whole genome resequencing^8-11^ have resulted in the global picture of common and rare SNPs across the genome. However, those data have been poorly used as an integrated manner to serve as haplotype information by imputation approaches that enrich the above SNP genotype data with whole genome SNP data^12^. In addition, genetic variation of wild soybean, which contains a large amount of untapped and unexplored soybean diversity, remains poorly characterized relative to that of domesticated soybean.

Here we analyze genomic variation of 781 soybean accessions consisting of 418 *G. max*, 345 *G. soja*, and 18 hybrid (*G. max* x *G. soja*) accessions obtained through high-coverage (>13×) whole-genome sequence data. We conducted the detection of domestication-selective sweeps and the identification of deleterious mutations in soybean populations, with the goal of providing soybean breeders with efficient means to approach untapped wild soybean alleles. We then showed the usefulness of our data in soybean genetics by imputing the SNP data set from SoySNP50K array genotyping^7,13^ using variants identified here for genome-wide association study (GWAS) of seed protein and oil traits.

## Results

### Genomic variation

We collected genome resequencing data for a total of 855 samples from 833 soybean accessions (Supplementary Table 1 and Supplementary Note). The 855 samples included 22 repeated samples that were added to examine the cause of high heterozygosity rate in some samples observed at the initial stage of this study. Of the 855 samples, 74 that showed higher than two thirds of heterozygous to homozygous non-reference SNPs ratios or inbreeding coefficient per individual of less than 0.8 were excluded from further population analyses (Supplementary Figs. 1 and 2). Final non-redundant 781 accessions as a haplotype map panel consisted of 418 *G. max* including 332 landraces and 86 improved lines, 345 *G. soja*, and 18 hybrid (*G. max* x *G. soja*) accessions. The *G. soja* and hybrid accessions were obtained from China, Korea, Japan, and the Russian Far East. The 781 data were mapped to the soybean Williams 82 reference genome ver. Wm82.a2.v1^6^ with mean depths ranging from 14.09 to 61.27 after removing duplicate reads and covered > 95.2% of the reference genome by more than one read and > 85.4% by more than fine reads for all accessions. After variant calling and filtration steps, we retained 10,597,683 high-quality SNPs to perform most of the population analyses (Supplementary Fig. 3, Supplementary Table 2, and Supplementary Note). In case of indel calls, 5,717,052 bi-allelic indels, approximately two-thirds of raw calls, were defined for population analyses of the genomes of the 781 accessions. The indels were then divided into 5,578,041 of small indels and 139,011 of structural variants (SV) (> 50 bp) (Supplementary Fig. 4).

### Population structure and diversity patterns

The population structure of the 781 soybean set assessed using the 10.5 million SNPs (Supplementary Figs. 5 to 6, Fig. 1, and Supplementary Note) was similar to that from our recent analysis of 3,016 non-redundant soybean accessions genotyped using the 180K SoyaSNP array^2^. However, unlike the tree topology constructed from 180K SNP array data that had some ascertainment bias that favored selection of *G. max* soybean SNPs^14^, branch length differences between *G. max* and *G. soja* in our phylogenetic tree (Fig. 1) reflected almost two times higher nucleotide diversity (π) in *G. soja* (0.0023) than *G. max* (0.0012) in our 781 soybean genome population with an intermediate level (0.0020) in hybrid. The results indicate that, consistent with the previous observations^10,15^, roughly half of the genetic diversity has been lost during domestication from wild (*G. soja*) to domesticated soybean, which supports the occurrence of a bottleneck in the genetic pool during the soybean domestication process.

**Fig. 1.**
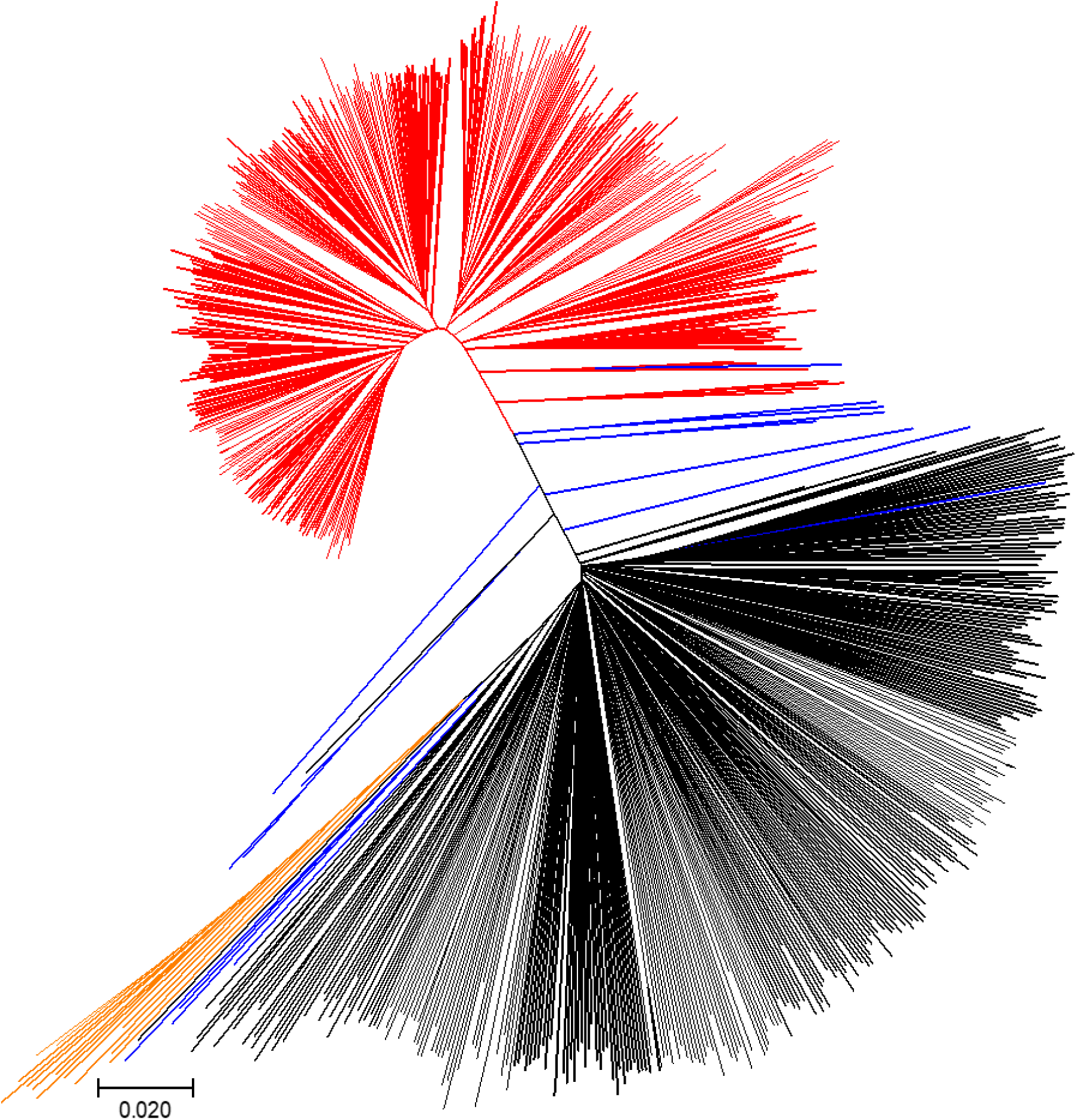
Neighbor-joining tree of the 781 haplotype soybean accessions. The accessions were divided into four color lines: *Glycine max* is red, most of *Glycine soja* black, *G. soja* collected from the middle region of the Yellow River basin orange, and hybrids blue.

Genome-wide profiling of variants identified in the 781 soybean accessions was performed on the Williams 82 reference genome to reveal diversity patterns in soybean (Fig. 2). Historical recombination rates (ρ) varied substantially along chromosomes, consistent with observations in other plants^16,17^. All chromosomes had lower recombination near the centromere repeat regions, which are presumed to be pericentromeric regions spanning more than 10 Mbp, relative to that in euchromatin regions. This pattern of recombination frequency distribution have been well supported experimentally by studies of multi-parental maize mapping populations^16,18^, although recombination rates were detected to be almost entirely suppressed in pericentromeric regions in those mapping populations. With available estimates of the recombination rate (*R*) from four soybean inter-crossed bi-parental populations, which captured ∼38,000 meiotic crossovers^19^, we compared our estimates of historical recombination rates with empirical estimates of the recombination rate. Overall, *R* and ρ were weakly, significantly correlated, indicating that our historical recombination rate estimates inferred on the basis of the SNP distribution likely reflected naturally occurring recombination patterns (Spearman correlation coefficient = 0.256, *P* = 7.945e-16).

**Fig. 2.**
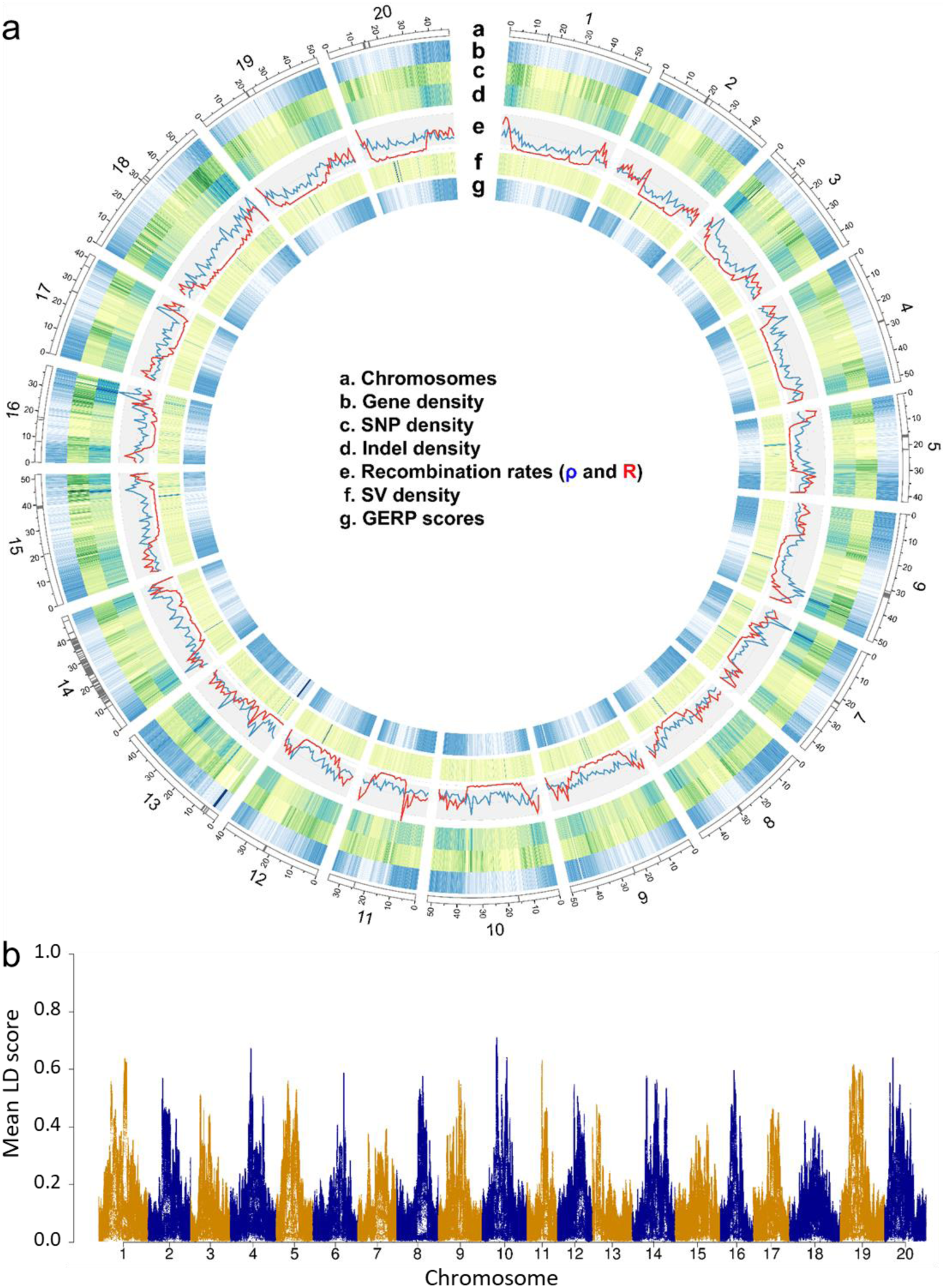
Genomic landscape of soybean. **a** Chromosomes based on the Williams 82 reference genome sequence v. Wm82.a2.v1. Centromere repeat regions are indicated by gray bands (a). Gene density heatmap (b). SNP density (c). Indel density (d). Population recombination rates calculated in 1 Mb windows (blue = historical recombination rate and red = estimates of recombination rate from mapping populations (e). SV density (f). Average GERP score density (> 0), with dark blue of high GERP score (g). All window sizes are 100 kb except recombination rates. **b** Mean LD scores estimated with a 1 Mb window.

The overall chromosomal distribution patterns of gene density, SNP density, indel density, and genomic evolutionary rate profiling (GERP) scores were similar to those of recombination rates (Fig. 2). A detailed description of GERP scores is provided below. The patterns of these variables we observed across the genome were significantly correlated with the highest correlation between gene density and GERP score density (Supplementary Table 3) and thus reflected other reports on plant genomes^20,21^. Interestingly, although correlation coefficients between historical recombination rates and other variables are relatively low with the highest value of 0.344 between recombination and indel density, our visual inspection indicated that the highest peak (hotspot) of historical recombination rates in each of half of the 20 chromosomes corresponded with the densest region of indels in each chromosome. A variety of genomic features such as gene density, CpG islands, and structural variants have been identified as being associated with regions of high recombination^17,18,22^. However, the association between high indel density and recombination hotspot has so far been poorly examined. Interestingly, when we searched indel motifs from genomic regions showing extremely high historical recombination rate (Rho), we found the ‘AARATA’ and ‘CTCHA’ motifs (Supplementary Fig. 7) occurred with 15.5% and 14.2% frequencies, respectively, while the other motifs showed less than 3.5%.

We then estimated the patterns of linkage disequilibrium (LD), which is strongly influenced by the mutation and recombination history among many factors. LD decay rates were higher in *G. soja* than *G. max*, while that of hybrid was in the middle of the two large groups (Supplementary Fig. 8). LD (indicated by *r*^*2*^) dropped to half of its maximum value at ∼ 7.9 kb in wild soybean, similar to those from previous studies in soybean^10^, rice (*Oryza. rufipogon*, 20 kb)^23^, and wild maize (*Z. mays ssp. parviglumis*, 22 kb)^24^. In the domesticated soybean, LD increased to 94 kb similar to that of predominantly selfing cultivated rice (123 kb and 167 kb in *indica* and *japonica*, respectively)^25^ but much higher than outcrossing cultivated maize (30 kb)^24^. We found that the local LD of pericentromeric regions was much higher than that of euchromatic regions (Fig. 2B). Thus, chromosomal distribution pattern of LD is negatively correlated with those of historical recombination rates, gene density, SNP density, indel density, and GERP scores (Supplementary Table 3).

### Signals of selection for domestication in soybean

Our dataset derived from a collection of 418 domesticated accessions and a comparable number of wild accessions provides an unprecedented opportunity for the scanning of selective sweep regions during domestication. Previously, several notable genome resequencing-based studies have used less than 100 wild accessions for genome scans^10,24,26^. To identify potential selective signals during soybean domestication (wild versus domesticated soybean), we scanned genomic regions with extreme allele frequency differentiation over extended linked regions using a likelihood test (the cross-population composite likelihood ratio, XP–CLR)^27^. A total of 183 domestication-selective sweep regions were detected (Fig. 3). Selective sweep regions had a mean size of 368 kb containing an average of 20 genes and accounted for 6.4% of coding sequence (CDS) in the soybean genome (7,215,740 bp of CDS for selective sweeps versus 104,886,718 bp CDS for the rest of the genome). They showed multiple signatures of selection, including elevated differentiation and an expected profile of nucleotide diversity reduction in domesticated soybean relative to wild soybean (Fig. 3). More selective sweep regions were detected on chromosomes 3, 5, 11, 13, and 20, consistent with previous results that used small numbers of wild soybean accessions^10,11^. A notable exception is two adjacent large selective sweep regions spanning roughly 13 Mb at the pericentromeric region of chromosome 1. In this region, both domesticated and wild soybean had low nucleotide diversity reflecting a general pattern of pericentromeric region in plant genomes. However, Tajima’ s D values for the domesticated soybean population were highly negative, indicating that this large pericentromeric region might contain key loci that have been selected for domestication.

**Fig. 3.**
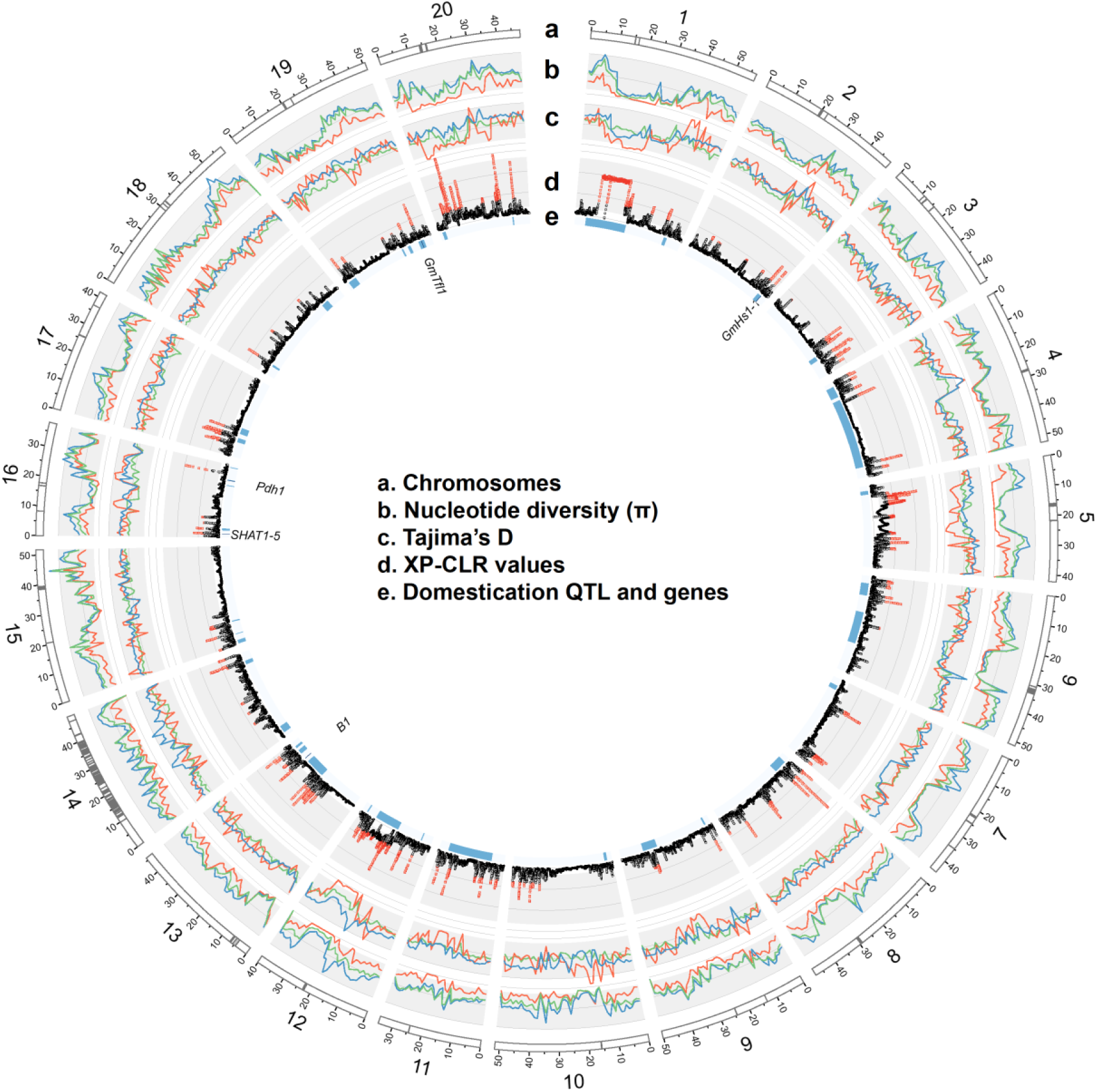
Genomic landscape of soybean. Chromosomes based on the Williams 82 reference genome sequence v. Wm82.a2.v1 (a). Centromere repeat regions are indicated by gray bands. Nucleotide diversity (*π*) in 1 Mb windows for each soybean subpopulation (red=*G. max*, dark blue=*G. soja*, green=hybrid) (b). Tajima’ s D for each soybean subpopulation (red=*G. max*, dark blue=*G. soja*, green=hybrid) (c). Distribution of genome-wide likelihood (XP-CLR) values for selection during domestication (d). Plot is based on XP-CLR scores of 100-kb block with 10-kb sliding windows. Domestication quantitative trait loci (QTL) and genes on chromosomes as detected in a large mapping population Williams 82 × PI 479752^28^. (QTL = blue bands and genes = dark blue bands) (**e**). Gene names are also shown.

When peaks on soybean chromosomes identified as putative selective sweeps were compared with domestication-related QTL from a recent comprehensive study using bi-parental (domesticated by wild) cross soybean populations (Fig. 3)^28^, the comparison supported the selective sweeps identified in this study. Out of 42 chromosomal regions containing unique and overlapping QTL, about 70% corresponded to chromosomal regions detected by XP-CLR. Among 17 QTL that had more than 5% of phenotypic variation explained (PVE), 13 corresponded to the selective sweep regions that were detected by XP-CLR. However, because several QTL spanned more than 20 Mb around pericentromeric regions that have low recombination rates, these comparisons should not be considered conclusive but rather suggestive of questions for further study. None of several genes that have been cloned with implication of domestication selection in soybean properly overlapped with XP-CLR peaks

### Enhanced genetic load in selective sweep regions

Deleterious alleles that are tightly linked to the strongly selected allele on selective sweeps may be less effectively purged relative to those on neutral backgrounds. Studies with several <u>predominant</u> or mandatory outcrossing species^21,29-31^ showed that process of domestication have resulted in an increased number of deleterious variants in the domesticated genome, providing a basis for the “cost of domestication” hypothesis^32,33^. Here, to quantify the extent of purifying selection on deleterious alleles in the self-compatible, predominantly selfing plant soybean, we used Sorting Intolerant From Tolerant (SIFT)^34^ and GERP^35^ scores. In soybean, of the 397,869 SNPs identified within coding sequences (CDS), 17.8% (70,795) were considered putatively deleterious (SIFT < 0.05). GERP^35^ scores were obtained by computing constraint for individual positions on the basis of comparative genomic approaches. To allow for a comparative analysis, genomic evolution and amino acid conservation modeling^21^ was used to catalog candidate deleterious variants across the soybean genome. GERP identified 237.5 Mb of the soybean genome (24.3%) as evolutionarily constrained (GERP > 0), and 111.5 Mb (11.4%) as highly evolutionarily constrained (GERP > 2) (Supplementary Fig. 9). As expected from the distribution pattern of GERP scores on chromosomes (Fig. 2), we found that 71.8% of the deleterious SNPs inside CDS were also evolutionarily constrained (GERP > 0) in soybean. GERP scores were combined with SIFT scores to identify the deleterious mutations in the constrained portions of the genome. As a result, the putative deleterious mutations (SIFT <0.05) in CDS were categorized into conserved deleterious (nonsynonymous, GERP ≥ 2,) and moderately-conserved deleterious (nonsynonymous, 0 < GERP < 2, SIFT < 0.05) mutations. Additionally, stop mutations (either gain or loss, GERP > 0) were categorized as a deleterious mutation group, although their SIFT scores were not available. We explored the mutation burden in domesticated and wild soybean populations. To examine impact of recent crossing and selfing, the domesticated soybean accessions were further divided into landraces and improved lines. Contrary to an increase of mutation burden in domesticated types predicted by the cost of domestication hypothesis, results showed approximately 25% to 35% decrease of overall deleterious alleles in landraces relative to wild soybean accessions and approximately 5% additional decrease in improved lines (Supplementary Fig. 10). Because the three initial categories showed similar tendencies among three soybean subpopulations, we combined the three categories into one (Fig. 4A), and used the combined group in subsequent analysis. Our results are in contrast to previous studies from cassava, grape, maize, rice, and sunflower^21,29-31,33,36^. Interestingly, our results are similar to substantial decrease of the homozygous-mutation burden in domesticated cassava and grape accessions, compared with progenitors^21,31^. Decrease from landraces to improved lines were observed between inbred elite maize lines and their landraces^37^.

**Fig. 4.**
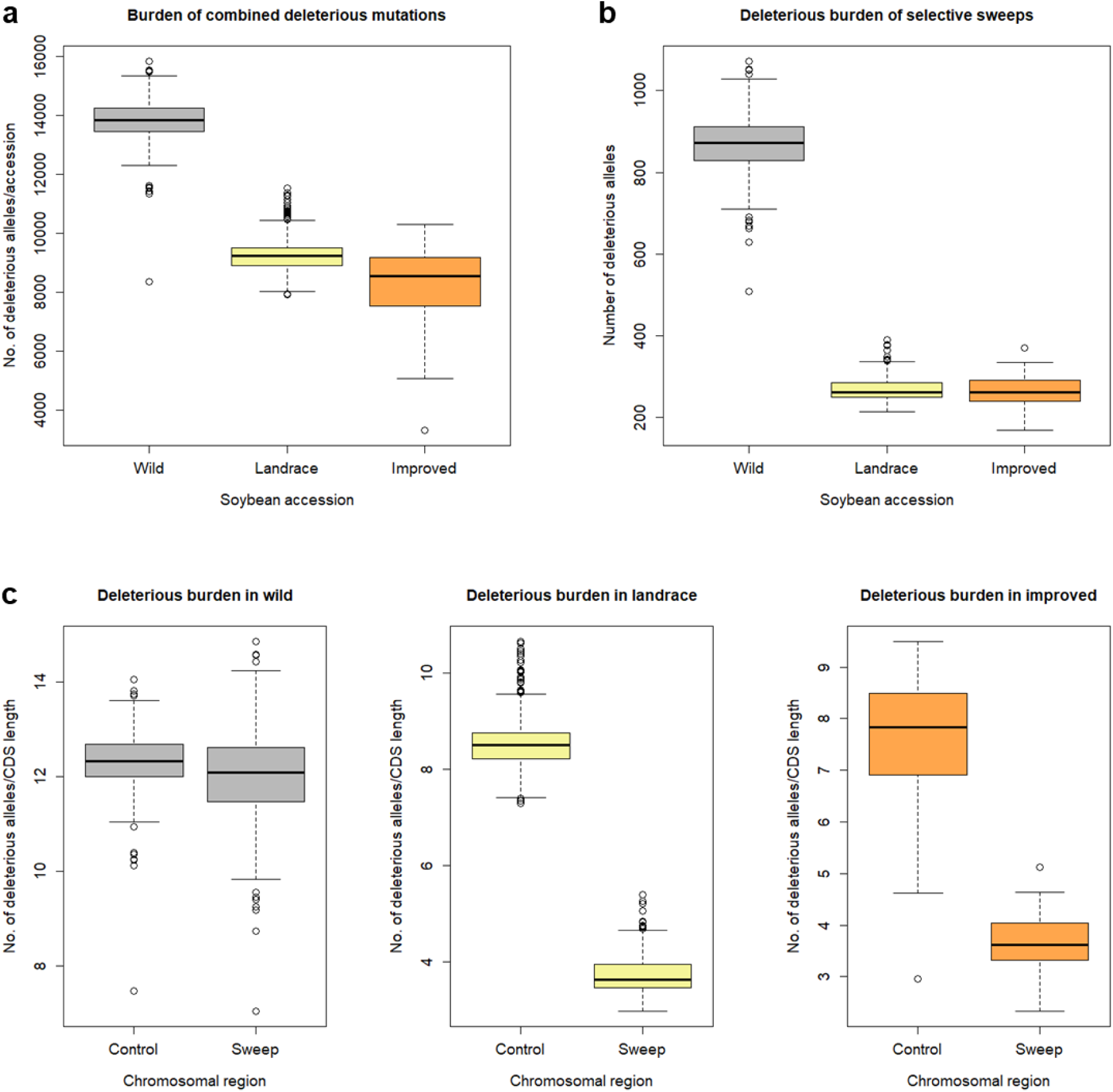
Box-and-whisker plot distributions of mutation burden in domesticated and wild soybean populations. **a** Total mutation burden in individual domesticated (*Glycine max*, landrace cultivars = 332 and improved lines = 86) and wild (*Glycine soja, n* = 345) soybean accessions. **b** Mutation burden among landrace, improved, and wild soybean accessions in domestication sweep regions. **c** Mutation burden in wild, landrace, and improved soybean accessions between domestication selective sweeps and control regions (rest of the genome). Vertical axis shows number of deleterious alleles per 100-kb CDS length. The subgroups in each of plots are significantly different between one another with *P* < 0.09e-5 in t-tests or Tukey multiple comparison tests except deleterious burdens between landrace and improved soybean accessions with *P* = 0.644.

We then compared selective sweep regions between the domesticated and wild soybean accessions. We found that domesticated soybean, compared with wild soybean, showed 69.7% (landraces) and 69.0% (improved lines) fewer (Fig. 4B) deleterious alleles in sweep regions. Thus, the decrease in deleterious alleles has likely been enhanced by artificial selection, suggesting the decreased mutation load we observe in soybean has been driven by the genome-wide effects of the domestication bottleneck as well as linkage associated with the selection of specific genes. However, total mutation burden between landraces and improved lines was significantly different while there is no significant difference between the two groups in selective sweeps, indicating artificial selection outside of selective sweeps during modern soybean breeding. In addition to the comparison between the domesticated and wild populations, within-population comparison of sweep regions with the rest of the genome in deleterious alleles showed that selective sweeps exhibited 56.5% (landraces) and 52.2% (improved lines) decreases (Fig. 4C) in deleterious alleles in domesticated soybean. As expected, nearly the same level of deleterious alleles (only 2.3% difference of means) was observed between sweep regions and the rest of the genome in wild soybean (Fig. 4C). Collectively, these results suggest that haplotypes containing fewer deleterious alleles have been favored during artificial selection. Unlike outcrossing species that maintained accumulated deleterious mutations in the heterozygous state, the predominantly selfing plant soybean has been less tolerable to accumulation of deleterious alleles. In other words, progeny that might more frequently inherit homozygous deleterious alleles from heterozygous deleterious allele parents by selfing rather than outcrossing have been naturally or artificially eliminated, thereby purging deleterious mutations from the domesticated soybean.

The selected mutations during domestication can be novel or standing genetic variation^38^. Novel domestication alleles such as those for the reduction of seed-shattering are deleterious in the wild and would be extremely rare in the wild plants. Standing domestication alleles such as those for seed size may be fixed in domesticated plants but are segregating in the wild. As the comparison between identified selective sweeps and domestication QTL supported their high correlation and the genetic load analysis in selective sweeps corroborated unique biological feature of soybean in domestication, we attempted to propose a list of 107 strong domestication candidate genes. Some of them are homologues of cloned canonical domestication genes including AP2 and PIF1 transcription factors^39^ that regulate plant architecture (Supplementary Table 4). However, in this approach, the trait or traits affected by the selected alleles may be ambiguous and novel mutation will generate a more conspicuous signature of a selective sweep than standing mutation. Thus, we highlighted 11 candidate novel mutation-containing genes whose deleterious alleles are almost fixed in the domesticated soybean and have low frequency in the wild soybean. Because none of them have been characterized with implication for domestication and three were annotated as proteins of unknown function, they are likely to be specific to soybean or eudicot crop plants.

### Uses of the haplotype data set for genomic association

A major objective for sequencing a large collection of accessions is to impute genotypes to improve existing GWAS by fine-mapping existing association signals and detecting new associations. We evaluated the usefulness of our haplotype data for GWAS by imputing our SNP data set into existing SoySNP50K genotyping and phenotyping data for seed protein and oil contents in a large population^7,13^. In soybean research, numerous linkage analysis and GWAS have been conducted for these two important traits^40,41^.

We re-analyzed the previous GWAS of seed protein and oil because of a substantial update of the soybean reference genome version and to eliminate many nearly identical accessions in the original 12,116 soybean accession set (Supplementary Figs. 11 to 12, Supplementary Table 5, and Supplementary Note). We then imputed 10.5 million SNPs into 36,647 SNPs from the SoySNP50K data of 8,844 non-redundant soybean accessions (Supplementary Fig. 13 and Supplementary Note). Imputation accuracies were approximately 97%. General patterns of GWAS results analyzed using a linear mixed model on imputed data on 8,844 accessions were quite similar to our re-analysis results on existing SoySNP50K array genotyping and phenotyping data (Fig. 5, Supplementary Fig. 13, and Supplementary Note). As expected, major peaks were clearly found for both seed oil and protein. Interestingly, more than 10 novel minor significant peaks such as those on chromosomes 2, 4, and 10 appeared for each of the oil and protein traits and their multivariate trait. Although they were clearly found, not a single SNP at these regions reached genome-wide significance in the previous GWAS. However, when we performed multi-locus mixed-model (MLMM) analysis on the same data set, none of the novel minor peaks were any longer significant (Supplementary Fig. 14), indicating that those minor peaks appeared likely to be due to the confounding partial LD effects of massively imputed SNPs. For the sake of simplicity for examining any improvement of our imputed GWAS, we focused on five significant major peaks on chromosomes 5, 8, 13, 15, and 20 from mvLMM (Fig. 5A), which were supported by both the MLM and MLMM models. Similar to the previous GWAS that used imputed data sets^12,42,43^, the general width and shape of the peaks detected from unimputed data remained largely the same as those from the imputed data with slightly longer tails as much as LD distances of boundary markers (Fig. 5B and Supplementary Fig. 15). The number of significant SNPs increased and the most significant SNPs showed improvement in signal strength and shifted in position in GWAS with imputed data. Among genes that have been reported as regulatory genes for oil content in soybean, the *GmSWEET39* (*Glyma*.*15g049200*) gene provided an opportunity to examine the improvement of our imputed GWAS because this was cloned as a gene controlling seed oil content by selection during soybean improvement and was suggested as the causal gene for the major association peak on chromosome 15^44^. The most significant SNP shifted 90.83 kb in position, from 3,937,899 to 3,847,069 bp (Fig. 5C), although the SNP is not located at the genic region of *GmSWEET39*. The most significant SNP does not necessarily correspond with variants from a causal gene of an association peak, as notably shown by a rice GWAS^45^. Interestingly, the *GmSWEET39* gene is located in a newly observed association region (approximately 85 kb) with 49 significant imputed SNPs (higher than –log_10_(*p*-value) of 20.09) in the middle of the chromosome 15 peak, which was non-significant valley in the GWAS with the original unimputed data (Fig. 5C). Because the *GmSWEET39* gene would not have been regarded as a candidate causal gene in the GWAS with unimputed data, this observation serves as apparent evidence that GWAS with imputed data had the potential benefit of resolving associations in soybean.

**Fig. 5.**
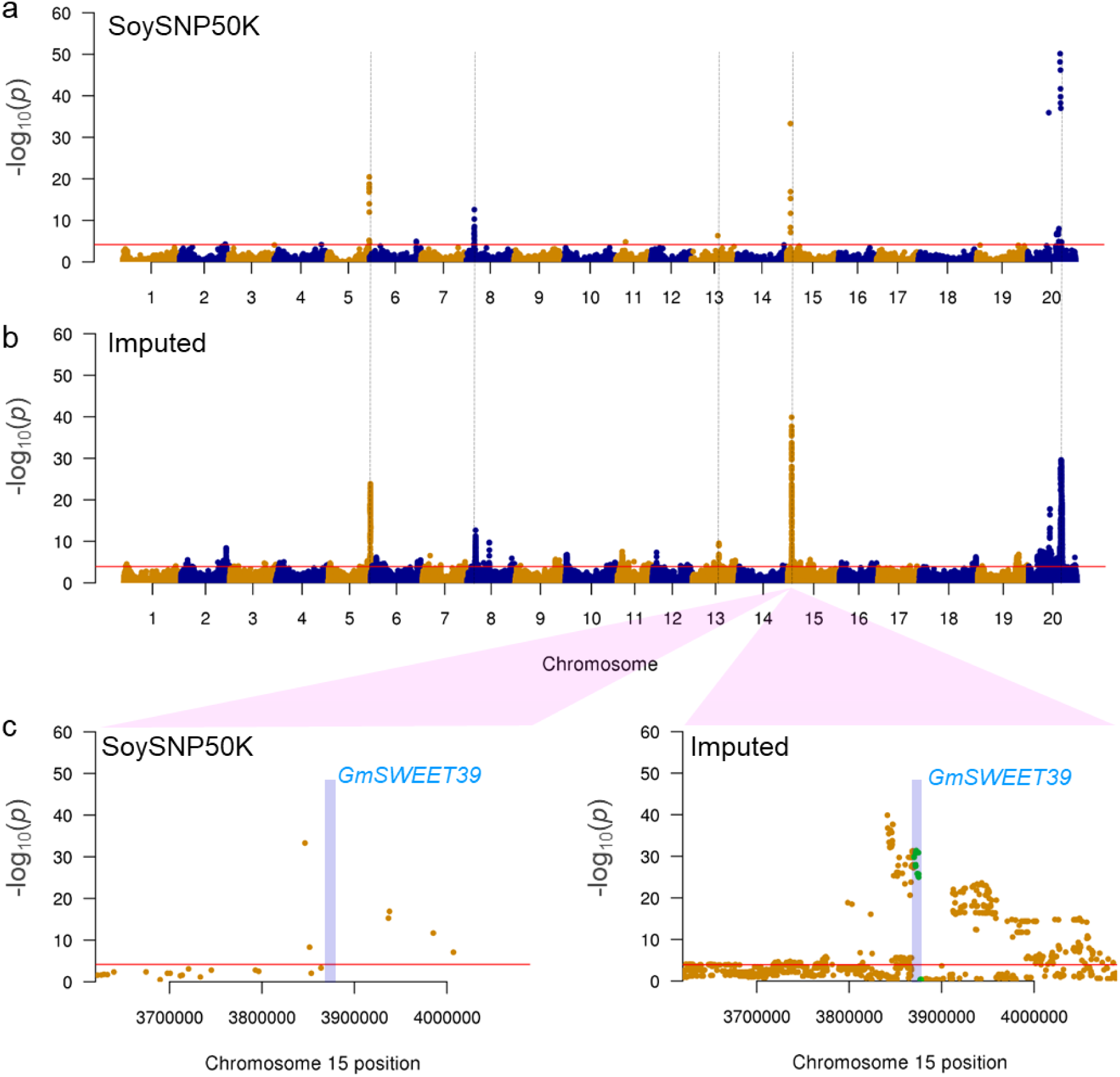
Comparison of mvMLM-based GWAS for oil and protein contents using unimputed and imputed genotype data. **a** Results using the original genotype data from SoySNP50K array. Horizontal line represent 5% significance thresholds corrected for multiple testing using Benjamini-Hochberg. Five major peaks are indicated by dashed vertical lines for comparison. **b** Results using imputed data that imputed 10.5 million SNP data from 781 soybean genomes into SoySNP50K data. **c** Comparison of mvMLM-based GWAS results using unimputed (SoySNP50K) and imputed genotype data at a major peak on chromosome 15. A pale blue box indicates a chromosomal region of oil content regulator GmSWEET39 that includes its genic region and 5 kb of each of its 5’ upstream and 3’ downstream regions. SNPs located in the GmSWEET39 region are also highlighted by green dots.

## Discussion

The discovery and characterization of extensive genome-wide genetic variation in the 781 diverse soybean accessions containing an unprecedented number of wild soybean accessions provided us with an opportunity to find unique features of plant genomes that were largely due to both wild and domesticated species being predominantly self-pollinating. The most striking feature is that more deleterious alleles were purged from domesticated than wild soybean accessions. During the past decade, genome-wide fine genetic variation of major crops including rice, maize, sorghum, cassava, and grape have been revealed^46^. However, those well-characterized major crops have different reproduction modes from soybean. Both wild and domesticated species of maize are predominantly outcrossing. Domesticated species of rice and sorghum tends to be selfing while their wild types are predominantly outcrossing. Both wild and domesticated species of cassava and grape are outcrossing, however cultivated types are predominantly clonally propagated. Therefore, the findings from this study may be extended to the characterization of wheat and barley^47,48^, which have the same reproduction mode as soybean but whose genome analyses have lagged behind due to their huge genome sizes.

Of the originally resequenced 855 samples, we excluded 74 samples (8.65%), which showed high heterozygosity and low inbreeding coefficient, based on presumed reproduction mode of soybean. The 781 soybean accessions were clearly divided into domesticated and wild accession groups with a distinct subgrouping of wild accessions according to geographic collection sites, in similar fashion to other major crops. However, compared to maize landraces that showed only 17% diversity reduction from their wild progenitor^24^, a drastic reduction in nucleotide diversity (∼ 48%) was observed during the transition from wild to domesticated soybean. This likely reflects different reproduction modes between selfing soybean and outcrossing maize. The overall chromosomal distribution patterns of variation of several variables were also quite similar to those observed in other major crops. Nevertheless, it is interesting to observe that the historical recombination hotspot in half of the 20 chromosomes corresponded with the densest indel region in each of those chromosomes. Discovery of the two highly frequent indel motifs suggested that there may be some relationship between indel mutagenesis and recombination.

A diverse collection of 345 wild soybean accessions were analyzed against 418 domesticated accessions to detect selective signals for soybean domestication. Although many canonical domestication genes have been cloned from major grass crop species, such knowledge has not been translated well to domestication research in eudicot seed crop species including soybean. In major crop species, cultivated species and their progenitors usually show distinct morphological and physiological differences in so-called domestication syndrome traits such as seed size, shattering, seed dormancy, flowering time, and viny growth habit. Soybean is not an exception. However, organs and tissues where several domestication traits are expressed differ between soybean and grasses. For example, shattering is related to the pod in soybean, but to pedicel in rice. Unfortunately, our analysis show that none of the soybean domestication genes cloned thus far should be regarded as a canonical domestication gene. However, in this study, we reported many candidate canonical domestication genes whose alleles are almost fixed in domesticated soybean and have low frequency in wild soybean.

In this selfing species, overall deleterious alleles among landraces relative to wild soybean accessions have been drastically reduced by up to almost 35%, similar to the observation in sorghum^20^. Mutation burden was further decreased in improved lines from modern soybean breeding. The results are in contrast to the cost of domestication hypothesis that deleterious alleles (the genetic load) that happen to be present in the neighborhood background of the strongly selected allele in the presence of selective sweeps may become more prevalent than those in other neutral backgrounds^32,33^. Purging of deleterious alleles from the domesticated soybean has been further enhanced in selective sweep regions. Unlike outcrossing species that maintained accumulated deleterious mutations in the heterozygous state, this predominantly selfing plant may have been less tolerable to the accumulation of deleterious alleles, eventually leading to the reduction of diversity. In other words, progeny that might inherit homozygous recessive deleterious alleles from heterozygous deleterious allele parents by selfing have been naturally or artificially eliminated, thereby purging deleterious mutations from domesticated soybean. Interestingly, artificial selection during modern soybean breeding have occurred outside of selective sweeps. Introgression of untapped variation in wild soybean should be an important objective for the future breeding of soybean. Most of the deleterious alleles have to be eliminated during breeding, although some deleterious alleles may provide beneficial effects for soybean growth or yield in crop fields. Information obtained here should help better design crossing and selfing efforts to efficiently eliminate unwanted deleterious alleles in a breeding program to select agronomically important untapped genes from wild soybean.

Finally, we have shown that our high-quality map of genome variation in soybean could be used as a reference panel for the imputation of genotypes to improve the existing GWAS for oil and protein traits. In addition to those unique genome variation features due to selfing and being a eudicot seed crop species that suggest soybean as a model for other such crops, our imputation results suggest that the soybean variation map and methods developed here can be used in a direct manner to accelerate genetic variation discovery in this economically important crop.

## Methods

### Plant materials and sequencing

We initially selected 818 accessions based on the 180 K Axiom^®^ SoyaSNP array genotyping data of ∼ 4,400 diverse soybean accessions, most of which were collected from South Korea. These diverse soybean accessions also contained representatives from the worldwide distribution of soybean^49^. Soybean plants were grown in the Ochang field of the Korea Research Institute of Bioscience and Biotechnology, Cheongju, Korea. Although more than two times of single plant selection for the SoyaSNP array genotyping had been performed^50^, we collected young leaves from a single plant of each accession and then extracted genomic DNA using the cetyltrimethylammonium bromide (CTAB) method^51^. DNA sequencing was performed at LabGenomics (Seongnam) or Macrogen (Seoul) companies in Korea. Paired-end sequencing libraries were constructed with an insert size of 500 bp using a TruSeq DNA PCR-Free kit (Illumina, San Diego, CA, USA) according to Illumina library preparation protocols. Libraries were then sequenced using Illumina HiSeq 2500 or 4000 platforms with 2 × 151-bp paired reads to a target coverage of 10X. Some accessions that showed high heterozygous variant levels were sequenced multiple times. We also added resequencing data of 16 accessions determined in our previous studies^8,52^ except IT182932 that was newly sequenced in the present study. Consequently, resequencing data from a total of 855 samples were used for initial variant calling in this study (Supplementary Table 1).

### Read mapping and variant calling

Short paired-end reads of 855 samples were quality checked using FastQC (http://www.bioinformatics.babraham.ac.uk/projects/fastqc/). We then essentially followed procedures described in the Genome Analysis Toolkit (GATK) Best Practices for data pre-processing and variant calling^53,54^. We used BWA (version 0.1.12) with default parameters except for –M option^55^ to map genomic reads from each accession against soybean Wm82.a2.v1 reference genome assembly^6^. Alignments were further checked for PCR duplicates using Picard tools (version 1.134) (http://picard.sourceforge.net/). We performed sorting operation, base recalibration, per-sample and joint variant callings, and variant filtration using GATK (version 4.0.1.2). Known variant sites for soybean extracted from NCBI dbSNP Build 144 (https://www.ncbi.nlm.nih.gov/projects/SNP/snp_summary.cgi?build_id=144) were used for base recalibration. Raw variant calling data were divided into SNPs and indels with SelectVariants function of GATK (v. 4.0.1.2). A total of 62,987,283 SNPs and 8,567,041 indels were identified from the analyses of the genomes of 855 samples. Quality filtering of raw SNP calls was performed using VariantFiltration in GATK according to the following criteria: ReadPosRankSum of < −2.0, MQRankSum < −2.0, polymorphism confidence scores (QUAL) < 30.0, genotype call quality divided by depth (QD) < 3.0, Phred-scaled P-value of Fisher exact test for strand (FS) > 30.0, mapping quality (MQ) < 30.0, total depth of coverage (DP) < 100, genotype-filter-expression depth of coverage (DP) < 5, and genotype-filter-expression genotype call quality (GQ) < 10.0. Bi-allelic variants were then selected using VCFtools (version 0.1.15)^56^. To exclude erroneous variants in repetitive regions, variants with high mapping depth (> 4X reads per sample, where X was mapping depth) in each sample were masked. Allele balance (AB) was calculated and variants with AB < 30 in heterozygous genotypes were masked^57^. The VCFtools was then used to remove markers that were monomorphic and markers with call rates < 50%. Up to this stage of filtration, 36.8 millions of SNPs were defined as candidate variants. In the 62.9 million raw SNP calls, some samples showed more heterozygous than homozygous non-reference alleles. Those samples still showed high heterozygous rates in the 36.8 million candidate SNP set. Thus, 66 samples that contained higher than two-third heterozygous to homozygous non-reference SNPs ratios among the samples with more than 0.5 million heterozygous SNPs in the raw SNP call set were excluded from further analyses. The inbreeding coefficient per individual was then calculated as the difference between the expected and the observed heterozygosity standardized by the expected heterozygosity under Hardy-Weinberg. Based on the assumption that pure inbred lines would show inbreeding coefficients of near 1.0, we additionally excluded eight wild samples that had < 0.8 inbreeding coefficients per individual in the 36.8 million candidate SNP set. Finally, 781 accessions were determined as our soybean genome variation study set. To perform population analyses using a set of 781 accessions, we further filtered these candidate SNPs using the VCFtools according to the following criteria: --non-ref-ac 1 --maf 0.01 --max-missing 0.8. Finally, we retained 10,597,683 high-quality SNPs from analyses of the genomes of 781 accessions. Filtering of raw indel calls was performed according to the following threshold criteria: ReadPosRankSum of < −20.0, QUAL < 30, QD < 2.0, and FS > 200. Bi-allelic variants were then retained. From this analysis of the genomes of 781 accessions, a filtered set of 5,717,052 indels were defined. The indels were then divided into small indels and structural variants (SV) with a cut-off of sequence length of 50 bp.

### Population structure and diversity pattern inference

Principal component analysis (PCA) was conducted to summarize the genetic structure and variation present in the 781 accessions using smartpca function in Eigensoft v7.2^58,59^. We plotted the first three PCs. We further used the model-based, Bayesian clustering software FastStructure v 1.0^60^ to estimate the population structure. FastStructure was run on default settings with 10-fold cross-validation for subpopulations (K) ranging from K = 2 to 12. Numbers of subpopulations were defined using the marginal likelihood function. We plotted the membership coefficient using DISTRUCT^61^. A neighbor-joining tree was constructed by MEGA7^62^ under the *p*-distances model.

Nucleotide diversity (π)^63^, SNP density, and Tajimas’ s D^64^ for 100 kb were calculated with the 10.5 million SNPs using vcftools --window-pi 100000, --SNPdensity 100000, and -- TajimaD 100000, respectively^56^. Indel and SV densities for the bi-allelic variants were calculated using vcftools --SNPdensity 100000. Population recombination rates (Rho, ρ) were calculated in the entire panel using the software FastEPRR^65^ and also to calculate the recombination rate for each of the gene windows used to build the evolutionary model (FastEPRR_segments.R). Linkage disequilibrium (LD) decay was calculated using PopLDdecay v3.31 with -MaxDist 1000 -MAF 0.05 -Miss 0.1 parameters^66^. Measures of LD (*r* ^*2*^) were calculated for the entire population, but also for each chromosome and subpopulation. Pairwise *r* ^*2*^ estimates were calculated from the unimputed SNP dataset with MAF > 0.1 and maximum missing rate < 0.1. LD scores were calculated using the Genome-wide Complex Trait Analysis (GCTA) suite with default settings^67,68^. Circos^69^ was used to display distributions of estimated variables on the Williams 82 reference genome ver. Wm82.a2.v1^6^.

To search indel motifs from high indel density regions that were highly correlated with recombination hotspots, we first designated regions with historical recombination rate (Rho) over 478.265, which was calculated by 3rd Quartile+1.5× IQR (Interquartile range): 1.5*(251.24-99.89)+251. Next, in order to see whether or not there are significant patterns in those indel sequences, we retrieved insertion or deletion sequences from 781 soybean resequencing matrix in these genome regions. We trimmed out any indel sequences with lengths below 7 bp and only used indel sequences with lengths between 8 and 50 bp. All 66,549 indel sequences were submitted as primary sequences for DREME analysis in the web-based motif search program^70^.

### Genome scan for selective signals

To scan selective signals over the soybean genome, we used a widely used cross-population composite likelihood ratio test (XP-CLR)^27^ updated by Hufford, et al.^24^. XP-CLR uses allele frequency differentiation between populations. A total of 763 soybean accessions consisting of 418 domesticated and 345 wild accessions were used for detecting selective sweep regions. Missing variants in our haplotype map data were imputed using the BEAGLE v5.0^71^ with the default option. Evidence for selection of domestication across the genome was evaluated by comparing domesticated versus wild soybean genomes. Individual SNPs were assigned at positions along with a recombinant inbred genetic map derived from a cross between *G. max* “Williams 82 K” and *G. soja* “IT182932”^19^. Markers located on the insertion of unanchored scaffolds or different chromosome segments as well as on chromosome segments whose physical or genetic orders were not collinear between the reference genome and our genetic maps were excluded from the genetic map. Coordinates of the soybean reference genome assembly Wm82.a2.v1 were applied to calculate genetic per physical distance between markers in the genetic map. XP-CLR was performed with the following criteria: -w1 0.0005 200 100 –p1 0.7. In other words, XP-CLR scores of 100 bp windows were calculated for a maximum of 200 SNPs per 0.05 cM genetic window. Markers with a correlation level > 0.7 were down-weighted. Manhattan plots of XP-CLR scores were constructed using qqman^72^ in R package or using Circos^69^. Windows with > 89.4 of XP-CLR values, accounting for 5% of the genome, were considered as selective sweep regions. Groups of adjacent windows with XP-CLR values not containing more than one window below this threshold were defined as a single sweep region. We assigned the gene closest to the window with the maximum XP-CLR score in each selective sweep region as the most likely candidate.

### Determination of effects of nucleotide variants

To predict functional effects of variants, we used Sorting Intolerant From Tolerant 4G (SIFT)^34^ to annotate the high-quality 10.5 million SNPs set. To create a soybean database, uniref90 (https://www.uniprot.org/, download date: Feb 9th, 2019) was used as a reference protein set. Annotation of *G. max* Wm82.a2.v1 was downloaded from EnsemblPlants (ftp://ftp.ensemblgenomes.org/pub/plants/release-44/gff3/glycine_max). Gff3 format was converted to Ensembl GTF format. Soybean SIFT4G database was constructed using SIFT4G_Create_Genomic_DB implemented in SIFT4G. SIFT scores ranged from 0 to 1, and any position with a SIFT score <0.05 was considered to be putatively deleterious.

### Genomic evolutionary rate profiling (GERP)

We estimated the individual burden of deleterious alleles based on the genomic evolutionary rate profiling (GERP) scores^35^ for each site in the soybean genome. GERP score reflects the strength of purifying selection based on constraint in a whole genome alignment of multiple plant species. For the whole genome alignment, we used the LASTz/MULTIz approach previously described for the alignment of 20 angiosperm genomes to *A. thaliana* reference^73^ with minor modifications. We aligned 12 soft repeat-masked genomes of *Arabidopsis thaliana* (TAIR10.1), *Cajanus cajan* (V1.0), *Lupinus angustifolius* (v1.0), *Medicago truncatula* (4.0), *Oryza sativa* (IRGSP_1.0), *Phaseolus vulgaris* (1_0), *Populus trichocarpa* (v3), *Prunus persica* (v2), *Vigna radiata* (ver6), *Zea mays* (v4) from RefSeq database (https://www.ncbi.nlm.nih.gov/refseq/), and *Vitis vinifera* (V2) from URGI database (https://urgi.versailles.inra.fr/Species/Vitis) to the *G. max (Wm82*.*a2*.*v1)* genome. Topology of the 12 species of interest was extracted from the whole phylogenetic tree using ete3 toolkit^74^. The phylogenetic tree was downloaded from Phylogenetic Resources files on Dryad database (https://datadryad.org/resource/doi:10.5061/dryad.63q27.2)^75^. The branch length (substitution per site) of the phylogenetic tree was calculated using phyloFit^76^ with four-fold degenerated sites of chromosome 1 in *G. max*. All alignment files (maf files) were merged using Multiz and converted to fasta format using maf2fasta program. Alignment gaps (-) in the reference genome (*G. max*) and sequences of the same position in other genomes were removed. Finally, we calculated GERP scores using gerpcol with –j option from GERP++^35^. Uncalculated positions were filled with “0” because neither GERP score of N nor n sequence position in *G. max* genome was calculated.

### Mutation load estimation

We estimated genome-wide mutation load using numbers of derived deleterious alleles identified in soybean accessions based on SIFT and GERP scores. From ∼ 10.5 million SNPs, we extracted 397,869 variants located inside the coding regions of soybean genes (CDS). We categorized these mutations into three categories of deleterious variants: stop (either gain or loss of stop codon) mutations, moderately-conserved deleterious mutations (SIFT < 0.05, 0 < GERP ≤ 0), and conserved deleterious mutations (SIFT < 0.05, GERP > 2). The criterion of GERP >2 to determine conservative site was proposed by Ramu, et al.^21^ based on the distribution of cassava GERP scores. For most of the mutation load analysis, a combined data set containing all three groups of deleterious mutations was used. We polarized derived and ancestral alleles for the 1,187,830 CDS SNPs using *Phaseolus vulgaris* (1_0) and *Vigna radiata* (ver6) genomes as outgroups. For each variant, the corresponding nucleotides in both the outgroup genomes were identified based on the whole-genome alignment for the GERP score calculation above. We then used the est-sfs software^77^ to infer the probability of the derived versus ancestral allelic state at a polymorphic site. We summarized the mutation load as the number of derived deleterious alleles in an accession^21,78^.

### Filtration and imputation of soybean data genotyped using SoySNP50K array

Genotype data in soysnp50k_wm82.a2_41317.vcf that consisted of 42,291 SNPs scored on 20,087 germplasm accessions using the Illumina Infinium SoySNP50K BeadChip^7^ were downloaded from SoyBase as of June 10, 2019^79^. In this data set, we corrected the genotypes of 3,494 reverse-oriented SNP sites in Glyma.Wm82.a2. We removed 96 SNPs presumed to be absent in Glyma.Wm82.a2 because they showed a base that was different from both reference and non-reference bases. We also removed 2 mitochondrial DNA SNPs. The resultant 42,193 then underwent further filtration. From the whole set, we selected a total of 12,116 accessions for GWAS of seed protein and oil content by Bandillo, et al.^13^. Of the 12,116 accessions, 559 with heterozygous rate > 0.05 or missing rate > 0.05 were removed. We calculated identical-by-descent (IBD) values for all pairwise comparisons among 11,557 *G. max* accessions using PLINK^80^. We considered pairs of accessions to be duplicates if they had an IBD > 0.98^81^. As a result, 3,272 duplicates were removed, leaving 8,844 non-duplicated accessions with high-quality genotype data. In this set of 8,844 accessions, SNPs with heterozygous rate > 0.02, minor allele frequency < 0.02, and missing rate > 0.10 were discarded from the genotype data, leaving a total of 36,647 high-quality SNPs for the imputation of soybean haplotype data and GWAS. Beagle (v5.0) was used for imputing genotypes at sites not on the 36,647 SoySNP50K data using 10.5 million SNP data from 781 soybean genomes. A genetic map constructed from a population of 233 recombinant inbred individuals derived from a cross between Williams 82K and IT182932^19^ was used as the fine-scale recombination map input for imputation.

### Genome-wide association analyses for oil and protein contents

For GWAS for seed protein and oil content on the 8,844 accessions using the original SoySNP50K data, of the 36,647 high-quality SNPs, 36,498 SNPs located on 20 soybean chromosomes were used. GWAS on the 12,116 accessions originally used by Bandillo, et al.^13^ was also conducted for a comparison using 37,142 SNPs located on the 20 soybean chromosomes with frequency > 2%. Missing variants were imputed using BEAGLE v5.0^82^ with default option. We used GEMMA^83^ to infer the correlation between each variant and seed oil and protein content. We first estimated a relatedness matrix from genotypes using the ‘-gk 1’ option in GEMMA. Then, we assessed evidence for correlation in a univariate linear mixed model (LMM) framework using the ‘-lmm 4’ option. We also assessed evidence for testing marker associations between oil and protein content as well as for estimating genetic correlations between oil and protein content in a multivariate linear mixed model (mvLMM). The Benjamini–Hochberg procedure^84^ was used to account for multiple testing by controlling the false discovery rate (FDR) at 5%. Manhattan plots were constructed to display GWAS results qqman^72^ in R package. The GWAS procedure for seed protein and oil content on the 8,844 accessions using genotype data that imputed 10.5 million SNPs into SoySNP50K array data of 8,844 accessions were essentially the same as that for unimputed SoySNP50K data.

A modified genome-wide approach^85^ for implementing a multi-locus mixed-model (MLMM) (Segura et al., 2012) to resolve association signals involving large-effect genes was used to further identify SNPs potentially associated with the oil and protein traits. The MLMM method relies on a simple, stepwise mixed-model regression procedure with forward selection and backward elimination while re-estimating the genetic and error variances at each step of the regression. This method may well lead to higher detection power and a lower FDR relative to traditional single-locus approaches. Because the imputed data appeared to exceed the computing power available, we reduced the number of markers by linkage disequilibrium (LD)-based marker pruning in PLINK software^80^. Briefly, we pruned markers from imputed data using the ‘--indep-pairwise 100 25 0.99’ option in PLINK. This option considers a window of 100 SNPs. calculates LD between each pair of SNPs in the window, and finally removes one of a pair of SNPs if the LD is greater than 0.99. Next, overlapping SNPs between the imputed data and SoySNP50K data that were deleted during pruning were added back to the pruned data, resulting into 804,281 markers for MLMM models.

### Data and materials availability

Although 16 of the original data have been released in conjunction with prior publications^8,52^, we uploaded raw reads in fastq format for all 855 final accessions to NCBI SRA with SRA accession number PRJNA555366. Large datasets including SNP, indel, and SV calls and SIFT scores, GERP scores, ancestral state of CDS SNP variants are available from figshare repository (https://figshare.com/projects/Soybean_haplotype_map_project/76110).

## Acknowledgments

This work was supported by the Next-Generation BioGreen 21 Program (PJ01321304), Rural Development Administration, and partly by the National Research Foundation grant (NRF-2018R1A2A2A05021904) funded by the Korea government and by the Korea Research Institute of Bioscience and Biotechnology Research Initiative Program. The work at Rural Development Administration was funded in part by Rural Development Administration Project No. PJ01155401. We thank S.H. Lee and S.T. Kang for assistance with selection of soybean samples and N. Kim for assistance with quality assessment of a part of raw data.

## Author contributions

S.C.J. and J.K.M. designed the research and generated the data from assistance from J.H.K., D.N.B., M.S.C., J.K., H.C.O., S.K.P. S.C.J. and M.A.G. analyzed the data with assistance from M.S.K, R.L., S.T.K, and J.H.P. S.C.J. wrote the manuscript with inputs from M.S.K., R.L., M.A.G, and J.K.M.

## Additional information

**Supplementary information** is available for this paper

**Correspondence and requests for materials** should be addressed to M.A.G., J.K.M. or S.C.J.

**Reprints and permissions information** is available.

## References

1. Lee, G.A., Crawford, G.W., Liu, L., Sasaki, Y. & Chen, X. Archaeological soybean (*Glycine max*) in East Asia: does size matter? PLoS One 6, e26720 (2011).

2. Jeong, S.C. et al. Genetic diversity patterns and domestication origin of soybean. Theor Appl Genet 132, 1179–1193 (2019).

3. Foyer, C.H. et al. Neglecting legumes has compromised human health and sustainable food production. Nat Plants 2, 16112 (2016).

4. Carlson, J.B. & Lersten, N.R. Reproductive morphology. in Soybeans: Improvement, production, and uses, 3rd edn (eds. Boerma, H.R. & Specht, J.E.) 59–95 (ASA, CSSA, and SSSA, Madison, 2004).

5. Charlesworth, D. & Willis, J.H. The genetics of inbreeding depression. Nat Rev Genet 10, 783–96 (2009).

6. Schmutz, J. et al. Genome sequence of the palaeopolyploid soybean. Nature 463, 178–83 (2010).

7. Song, Q. et al. Fingerprinting Soybean Germplasm and Its Utility in Genomic Research. G3 (Bethesda) 5, 1999–2006 (2015).

8. Chung, W.H. et al. Population structure and domestication revealed by high-depth resequencing of Korean cultivated and wild soybean genomes. DNA Res 21, 153–67 (2014).

9. Fang, C. et al. Genome-wide association studies dissect the genetic networks underlying agronomical traits in soybean. Genome Biol 18, 161 (2017).

10. Zhou, Z. et al. Resequencing 302 wild and cultivated accessions identifies genes related to domestication and improvement in soybean. Nat Biotechnol 33, 408–14 (2015).

11. Valliyodan, B. et al. Landscape of genomic diversity and trait discovery in soybean. Sci Rep 6, 23598 (2016).

12. Wang, D.R. et al. An imputation platform to enhance integration of rice genetic resources. Nat Commun 9, 3519 (2018).

13. Bandillo, N. et al. A population structure and genome-wide association analysis on the USDA soybean germplasm collection. The Plant Genome 8 (2015).

14. Lee, Y.G. et al. Development, validation and genetic analysis of a large soybean SNP genotyping array. Plant J 81, 625–36 (2015).

15. Lam, H.M. et al. Resequencing of 31 wild and cultivated soybean genomes identifies patterns of genetic diversity and selection. Nat Genet 42, 1053–9 (2010).

16. Gore, M.A. et al. A first-generation haplotype map of maize. Science 326, 1115–7 (2009).

17. Marand, A.P. et al. Historical meiotic crossover hotspots fueled patterns of evolutionary divergence in rice. Plant Cell 31, 645–662 (2019).

18. Rodgers-Melnick, E. et al. Recombination in diverse maize is stable, predictable, and associated with genetic load. Proc Natl Acad Sci U S A 112, 3823–8 (2015).

19. Lee, K. et al. Chromosomal features revealed by comparison of genetic maps of *Glycine max* and *Glycine soja*. Genomics 112, 1481–1489 (2020).

20. Lozano, R. et al. Comparative evolutionary analysis and prediction of deleterious mutation patterns between sorghum and maize. bioRxiv (2019).

21. Ramu, P. et al. Cassava haplotype map highlights fixation of deleterious mutations during clonal propagation. Nat Genet 49, 959–963 (2017).

22. Rowan, B.A. et al. An Ultra High-Density Arabidopsis thaliana Crossover Map That Refines the Influences of Structural Variation and Epigenetic Features. Genetics 213, 771–787 (2019).

23. Huang, X. et al. A map of rice genome variation reveals the origin of cultivated rice. Nature 490, 497–501 (2012).

24. Hufford, M.B. et al. Comparative population genomics of maize domestication and improvement. Nat Genet 44, 808–11 (2012).

25. Huang, X. et al. Genome-wide association studies of 14 agronomic traits in rice landraces. Nat Genet 42, 961–7 (2010).

26. Qi, J. et al. A genomic variation map provides insights into the genetic basis of cucumber domestication and diversity. Nat Genet 45, 1510–5 (2013).

27. Chen, H., Patterson, N. & Reich, D. Population differentiation as a test for selective sweeps. Genome Res 20, 393–402 (2010).

28. Swarm, S.A. et al. Genetic dissection of domestication-related traits in soybean through genotyping-by-sequencing of two interspecific mapping populations. Theor Appl Genet 132, 1195–1209 (2019).

29. Wang, L. et al. The interplay of demography and selection during maize domestication and expansion. Genome Biol 18, 215 (2017).

30. Marsden, C.D. et al. Bottlenecks and selective sweeps during domestication have increased deleterious genetic variation in dogs. Proc Natl Acad Sci U S A 113, 152–7 (2016).

31. Zhou, Y., Massonnet, M., Sanjak, J.S., Cantu, D. & Gaut, B.S. Evolutionary genomics of grape *(Vitis vinifera* ssp. vinifera) domestication. Proc Natl Acad Sci U S A 114, 11715–11720 (2017).

32. Moyers, B.T., Morrell, P.L. & McKay, J.K. Genetic costs of domestication and improvement. J Hered 109, 103–116 (2018).

33. Lu, J. et al. The accumulation of deleterious mutations in rice genomes: a hypothesis on the cost of domestication. Trends Genet 22, 126–31 (2006).

34. Vaser, R., Adusumalli, S., Leng, S.N., Sikic, M. & Ng, P.C. SIFT missense predictions for genomes. Nat Protoc 11, 1–9 (2016).

35. Davydov, E.V. et al. Identifying a high fraction of the human genome to be under selective constraint using GERP++. PLoS Comput Biol 6, e1001025 (2010).

36. Renaut, S. & Rieseberg, L.H. The Accumulation of Deleterious Mutations as a Consequence of Domestication and Improvement in Sunflowers and Other Compositae Crops. Mol Biol Evol 32, 2273–83 (2015).

37. Yang, J. et al. Incomplete dominance of deleterious alleles contributes substantially to trait variation and heterosis in maize. PLoS Genet 13, e1007019 (2017).

38. Larson, G. et al. Current perspectives and the future of domestication studies. Proc Natl Acad Sci U S A 111, 6139–46 (2014).

39. Dong, Z., Alexander, M. & Chuck, G. Understanding grass domestication through maize mutants. Trends Genet 35, 118–128 (2019).

40. Patil, G. et al. Molecular mapping and genomics of soybean seed protein: a review and perspective for the future. Theor Appl Genet 130, 1975–1991 (2017).

41. Lee, S. et al. Genome-wide association study of seed protein, oil and amino acid contents in soybean from maturity groups I to IV. Theor Appl Genet 132, 1639–1659 (2019).

42. Tian, F. et al. Genome-wide association study of leaf architecture in the maize nested association mapping population. Nat Genet 43, 159–62 (2011).

43. Genomes Project, C. et al. An integrated map of genetic variation from 1,092 human genomes. Nature 491, 56–65 (2012).

44. Miao, L. et al. Natural variation and selection in GmSWEET39 affect soybean seed oil content. New Phytol 225, 1651–1666 (2020).

45. Yano, K. et al. Genome-wide association study using whole-genome sequencing rapidly identifies new genes influencing agronomic traits in rice. Nat Genet 48, 927–34 (2016).

46. Gaut, B.S., Seymour, D.K., Liu, Q. & Zhou, Y. Demography and its effects on genomic variation in crop domestication. Nat Plants 4, 512–520 (2018).

47. International Wheat Genome Sequencing, C. et al. Shifting the limits in wheat research and breeding using a fully annotated reference genome. Science 361 (2018).

## References

48. Mascher, M. et al. A chromosome conformation capture ordered sequence of the barley genome. Nature 544, 427–433 (2017).

49. Jeong, S.C. et al. Genetic diversity patterns and domestication origin of soybean. Theor Appl Genet (2018).

50. Jeong, N., Moon, J.K., Kim, H.S., Kim, C.G. & Jeong, S.C. Fine genetic mapping of the genomic region controlling leaflet shape and number of seeds per pod in the soybean. Theor Appl Genet 122, 865–74 (2011).

51. Saghai-Maroof, M.A., Soliman, K.M., Jorgensen, R.A. & Allard, R.W. Ribosomal DNA spacer-length polymorphisms in barley: mendelian inheritance, chromosomal location, and population dynamics. Proc Natl Acad Sci U S A 81, 8014–8 (1984).

52. Ilut, D.C. et al. Identification of haplotypes at the Rsv4 genomic region in soybean associated with durable resistance to soybean mosaic virus. Theor Appl Genet 129, 453–68 (2016).

53. DePristo, M.A. et al. A framework for variation discovery and genotyping using next-generation DNA sequencing data. Nat Genet 43, 491–8 (2011).

54. Van der Auwera, G.A. et al. From FastQ data to high confidence variant calls: the Genome Analysis Toolkit best practices pipeline. Curr Protoc Bioinformatics 43, 11 10 1–33 (2013).

55. Li, H. & Durbin, R. Fast and accurate short read alignment with Burrows-Wheeler transform. Bioinformatics 25, 1754–60 (2009).

56. Danecek, P. et al. The variant call format and VCFtools. Bioinformatics 27, 2156–8 (2011).

57. Krumm, N. et al. Excess of rare, inherited truncating mutations in autism. Nat Genet 47, 582–8 (2015).

58. Patterson, N., Price, A.L. & Reich, D. Population structure and eigenanalysis. PLoS Genet 2, e190 (2006).

59. Price, A.L. et al. Principal components analysis corrects for stratification in genome-wide association studies. Nat Genet 38, 904–9 (2006).

60. Raj, A., Stephens, M. & Pritchard, J.K. fastSTRUCTURE: variational inference of population structure in large SNP data sets. Genetics 197, 573–89 (2014).

61. Rosenberg, N.A. DISTRUCT: a program for the graphical display of population structure. Molecular Ecology Notes 4, 137–138 (2004).

62. Kumar, S., Stecher, G. & Tamura, K. MEGA7: Molecular Evolutionary Genetics Analysis Version 7.0 for Bigger Datasets. Mol Biol Evol 33, 1870–4 (2016).

63. Tajima, F. Evolutionary relationship of DNA sequences in finite populations. Genetics 105, 437–60 (1983).

64. Tajima, F. Statistical method for testing the neutral mutation hypothesis by DNA polymorphism. Genetics 123, 585–95 (1989).

65. Gao, F., Ming, C., Hu, W. & Li, H. New Software for the Fast Estimation of Population Recombination Rates (FastEPRR) in the Genomic Era. G3 (Bethesda) 6, 1563–71 (2016).

66. Zhang, C., Dong, S.S., Xu, J.Y., He, W.M. & Yang, T.L. PopLDdecay: a fast and effective tool for linkage disequilibrium decay analysis based on variant call format files. Bioinformatics (2018).

67. Yang, J. et al. Genetic variance estimation with imputed variants finds negligible missing heritability for human height and body mass index. Nat Genet 47, 1114–20 (2015).

68. Yang, J., Lee, S.H., Goddard, M.E. & Visscher, P.M. GCTA: a tool for genome-wide complex trait analysis. Am J Hum Genet 88, 76–82 (2011).

69. Krzywinski, M. et al. Circos: an information aesthetic for comparative genomics. Genome Res 19, 1639–45 (2009).

70. Bailey, T.L. DREME: motif discovery in transcription factor ChIP-seq data. Bioinformatics 27, 1653–9 (2011).

71. Browning, S.R. & Browning, B.L. Rapid and accurate haplotype phasing and missing-data inference for whole-genome association studies by use of localized haplotype clustering. Am J Hum Genet 81, 1084–97 (2007).

72. Turner, S.D. qqman: an R package for visualizing GWAS results using Q-Q and manhattan plots. Journal of Open Source Software 3, 731 (2018).

73. Hupalo, D. & Kern, A.D. Conservation and functional element discovery in 20 angiosperm plant genomes. Mol Biol Evol 30, 1729–44 (2013).

74. Huerta-Cepas, J., Serra, F. & Bork, P. ETE 3: Reconstruction, Analysis, and Visualization of Phylogenomic Data. Mol Biol Evol 33, 1635–8 (2016).

75. Zanne, A.E. et al. Three keys to the radiation of angiosperms into freezing environments. Nature 506, 89–92 (2014).

76. Siepel, A. & Haussler, D. Phylogenetic estimation of context-dependent substitution rates by maximum likelihood. Mol Biol Evol 21, 468–88 (2004).

77. Keightley, P.D. & Jackson, B.C. Inferring the Probability of the Derived vs. the Ancestral Allelic State at a Polymorphic Site. Genetics 209, 897–906 (2018).

78. Fu, W. et al. Analysis of 6,515 exomes reveals the recent origin of most human protein-coding variants. Nature 493, 216–20 (2013).

79. Grant, D., Nelson, R.T., Cannon, S.B. & Shoemaker, R.C. SoyBase, the USDA-ARS soybean genetics and genomics database. Nucleic Acids Res 38, D843–6 (2010).

80. Purcell, S. et al. PLINK: a tool set for whole-genome association and population-based linkage analyses. Am J Hum Genet 81, 559–75 (2007).

81. Anderson, C.A. et al. Data quality control in genetic case-control association studies. Nat Protoc 5, 1564–73 (2010).

82. Browning, B.L. & Browning, S.R. Genotype Imputation with Millions of Reference Samples. Am J Hum Genet 98, 116–26 (2016).

83. Zhou, X. & Stephens, M. Genome-wide efficient mixed-model analysis for association studies. Nat Genet 44, 821–4 (2012).

84. Benjamini, Y. & Hochberg, Y. Controlling the False Discovery Rate: A Practical and Powerful Approach to Multiple Testing. J R Stat Soc Series B Methodol 57, 289–300 (1995).

85. Lipka, A.E. et al. Genome-wide association study and pathway-level analysis of tocochromanol levels in maize grain. G3 (Bethesda) 3, 1287–99 (2013).

