## Supplemental Note and Supplemental Figures and Tables for "Purging of deleterious mutations during domestication in the predominant selfing crop soybean"

### Supplementary Note

**Variation calling and exclusion of highly heterozygous genotype samples.** Accession information and statistics of genome sequencing analyses for a total of 855 samples from 833 soybean accessions are summarized in Supplementary Table S1. Analysis of sequencing data obtained at the initial stage in this study indicated that 17 soybean accessions including 12 wild accessions contained higher than 0.5 million heterozygous SNPs in the initial, unfiltered SNP call data (Supplementary Fig. 1). Seven of them contained more heterozygous than homozygous non-reference alleles, in contrast to a simple assumption that an inbred line that had gone through at least two generations of single-seed descent for the current study[1] would show a much lower number of heterozygous genotypes than homozygous genotypes. Such high heterozygous rate did not change after further filtering with the initial set of samples. To test whether such results might be due to experimental errors other than the natural conditions of plants, we repeated the sequencing data analysis using 22 samples prepared under different conditions from the 17 soybean accessions (Supplementary Fig. 1). However, the high heterozygous rate did not change except for one single plant and two filial plant bulk trials. The results suggested that those high heterozygous soybean samples resequenced were likely progeny from cross between different accessions, in contrast to the notion that soybean is predominantly selfing. In results, we analyzed a total of 855 samples including 22 repeated samples. The 855 genomes were aligned to the *G. max* cv. Williams 82 reference genome ver. Wm82.a2.v1[2]. Raw variant calling data from the Genome Analysis Toolkit (GATK) Best Practices[3] contained a total of 62,987,283 SNPs and 8,567,041 indels. In light of the observation of high heterozygosity, of the 855 samples that included duplicated samples, 66 samples from 54 accessions that contained higher than two-thirds of heterozygous to homozygous non-reference SNPs ratios were grouped as high heterozygous samples. GATK quality filtering and further filtration steps including read depth per samples and allele balance reduced the number of candidate SNPs to 36.8 millions. The relative heterozygosity rate in each sample tended to remain to be similar after these filtrations. Distributions of inbreeding coefficient per individuals in subgroups divided by species and heterozygosity indicated that most samples grouped by the high heterozygosity showed an inbreeding coefficient of less than 0.8 before and after filtration of SNPs with inbreeding coefficient per marker less than 0 (Supplementary Fig. 2). However, eight wild accessions, which were not excluded on this criterion, showed inbreeding coefficient per individual of less than 0.8. Thus, these eight accessions were additionally excluded from further population analyses. Thus, we ended up with the 781 non-redundant accessions consisting of 418 *G. max*, 345 *G. soja*, and 18 hybrid (*G. max* x *G. soja*) accessions.

We excluded 74 samples (8.65%) of the originally resequenced 855 samples. The exclusion rate is much higher than natural cross-pollination rates of soybean below 1% revealed by many well-designed experiments (for a review, see Lu[4,] Carlson and Lersten[5]). However, higher than 5% cross-pollination rates were reported based on a different way of calculation[6] or in a certain environment[7]. This, those highly heterozygous accessions may have higher cross-pollination rates than normal or have had higher cross-pollination rate in a habitat with frequent visits by pollinating insects.

To perform most of the population analyses using a set of 781 accessions, we further filtered SNPs with > 20% missing, > 10% heterozygosity, and < 1% minor allele frequency (Supplementary Fig. 3) to retain 10,597,683 high-quality SNPs (Supplementary Table 2). Raw indel calls underwent GATK quality filtering, and only bi-allelic variants were then retained. A resultant set of 5,717,052 indels, approximately two-thirds of raw calls, were defined for population analyses of the genomes of the 781 accessions. The indels were then divided into 5,578,041 small indels and 139,011 structural variants (SV) (> 50 bp) (Supplementary Fig. 4). The utility of our high-quality genome-wide variation set as a true soybean haplotype map was assessed by evaluating the improvement in the power of screening genetic variants known to control complex agronomic traits. Here we focus on the application of our genetic variant data to genomic regions that govern domestication and seed oil and protein traits, which are arguably the most frequently studied traits in soybean research.

**Population structure.** PCA[8] and fastSTRUCTURE[9] were used to infer population structure of the 781 soybean set using 10.5 million SNPs (Fig. 1). The population structure of the 781 soybean set was similar to that observed in our previous analysis of 3,016 non-redundant soybean accessions genotyped using the 180K SoyaSNP array[10]. As observed in the previous study, the 781 accessions were clearly divided into one *G. max* group and two *G. soja* groups (Supplementary Fig. 5) with a distinct subgroup of *G. soja* accessions collected from the middle region of the Yellow River basin. Both scree plot from the PCA and the estimated cross-validation (CV) error plot from ADMIXTURE that showed steep slopes up to $K = 3$ supported the presence of three distinct groups (Supplementary Fig. 6), although the slopes did not level off likely due to subgroupings within the large groups. While the majority of South Korean domesticated accessions formed a dense subgrouping due to recent overcollection, *G. max* accessions in general did not show distinct subgrouping on a geographic basis. Relative to landraces, the improved cultivars also appeared to be narrowly clustered within the landrace group. The groupings of wild soybean accessions were consistent with the geographic distributions of the collection sites, although several Korean accessions were grouped with a small number of Japanese accessions unlike the previous study. A wild accession from Taiwan was grouped together with Chinese accessions. Two hybrid accessions that had slightly higher than 30% *G. max* genomic fraction at $K = 2$, which was our threshold for the designation of admixture, appeared at the margin of the wild soybean group in the PCA plot. Thus, by defining a hybrid subpopulation, we show two distinctly separated soybean subpopulations, which are *G. max* and *G. soja*.

The population relationship inferred from Neighbor joining tree analysis is consistent with results obtained from our PCA and admixture analyses (Fig. 1 of the main text). The tree showed that, relative to *G. soja* accessions, *G. max* accessions formed a monophyletic cluster. Hybrid accessions appeared to cluster between the two large groups, while some accessions that have d*isproportionately high portions* of *G. soja* or *G. max* genomic fractions clustered within *G. soja* or *G. max* groups, respectively. The tree topology is also consistent with those obtained using the 180K SNP array data except the terminal branch lengths. The branch lengths of wild soybean accessions tended to be much longer than those of *G. max*, confirming that similar branch lengths between *G. max* and *G. soja* in our recent analysis of SNP array data[10] were due to ascertainment bias that more SNPs were selected from *G. max* than from *G. soja*[11].

**Comparison between selective sweeps and locations of cloned genes in soybean.** Although most of the major, canonical domestication genes have been cloned in grass crop species including maize and rice[12], only four genes have been cloned with implication of domestication selection in soybean[13-16]. Among the four, the chromosomal location of only one gene, *Bloom1*, corresponded with a XP-CLR peak. The other three genes, GmHs1-1, SHAT1-5, and *GmTfl1*, did not overlap with XP-CLR peaks but were located right next to those peaks. In fact, hard-seededness determined by *GmHs1-1*, seed coat shininess by *Bloom1*, and stem growth determinacy by *GmTfl1* are segregating in domesticated soybean population and should be regarded as improvement genes. Swarm, et al.[17] speculated that SHAT1-5, which was cloned on the basis of comparative homologue analyses to *Arabidopsis* genes, may not be a canonical domestication gene because of no detection of QTL in their population and, instead, *Pdh1* cloned from a cross between domesticated accessions[18] is likely a domestication gene. However, we did not detect any overlapping between these two genes and our XP-CLR peaks.

**Re-analysis of existing GWAS with SoySNP50K array genotyping and phenotyping data for oil and protein contents.** A recent study[19] reported an extensive protein and oil GWAS that analyzed the accumulated historical phenotypic data from USDA GRIN database (https://www.ars-grin.gov/) and the SoySNP50K data[20] from 12,116 soybean accessions . We re-analyzed the previous GWAS because the reference genome version was updated from Glyma1 to Wm82.a2.v1 and because we noticed that the 12,116 soybean accession set contained many nearly identical accessions. With a cut-off threshold of IBD > 0.98, we removed 3,272 of nearly identical accessions. We then performed PCA on the resultant 8,844 accessions. Scree plot from principal component (PC) analysis of the filtered 8,844 set steeply decreased up to PC number = 3 and did not level off up to PC number = 5 (Supplementary Fig. 11). The results are consistent with previous studies that large *G. max* germplasm collections were divided into three large subpopulations and two additional small subpopulations[10,19], indicating that updating and subsequent further filtration of markers and removal of redundant accessions did not change the structure of the original population.

To test whether the removal of redundant accessions improved GWAS resolution, we performed GWAS in both the 12,116 and the 8,844 accession sets using both univariate LMM and multivariate LMM (mvLMM) models. GWAS resulted in similar significant associations between both the sets in general (Supplementary Fig. 12). However, $-\log_{10}(P)$ values of the most significant SNPs in major peaks are higher in the original 12,116 set than the 8,844 set and more minor significant peaks appeared in in the original 12,116 set relative to the 8,844 set. The results indicated that filtering redundant samples improved the resolution of our GWAS likely due to reducing the exaggeration of association signals from repeated use of nearly duplicated samples[21].

Our GWAS results on the 8,844 accessions were quite similar to the previously reported[19] results on the 12,116 accessions with two notable exceptions. First, a minor peak on chromosome 8 for oil in the previous study, whose existence was not mentioned in the report likely because of being too minor, appeared to be a clearly visible major peak in our 8,844 set GWAS. Second, we observed a novel peak at 21 Mb on chromosome 20. We think that these novel peaks likely appeared due to the update of the soybean reference genome sequence. Notably, a sequence contig (~ 3.4 kb) containing the peak SNP (BARC_1.01_Gm_20_30930931_A_G) was displaced from the well-known major peak position at 30 Mb to the 21 Mb position during the version update from the soybean reference genome Glyma1 to Ws82.a2.v1 (Supplementary Table S5). Because no significant linkage or association signal for oil or protein contents has been reported in this region so far, we think that the additional major peak at chromosome 20 is likely to be an artifact related to the assembly problem of the soybean reference genome Ws82.a2.v1. Thus, in subsequent analyses, we did not pay attention to this peak. Interestingly, the major peak on chromosome 8 is likely true positive because this peak was recently detected as a major QTL for oil content from an interspecific soybean mapping population[22].

It is well known that seed protein content has a negative relationship with seed oil content in soybean[23], and Bandillo, et al.[19] also reported negative relationships between oil and protein association signals. Thus, we applied mvLMM model to identify genomic regions that exhibited pleiotropy for protein and oil contents. Distribution of significant SNPs from mvLMM model on the Manhattan plot looked like putting oil-LMM GWAS on protein-LMM GWAS. Among five major peaks, peaks on chromosomes 5 and 8 appear to be oil-specific peaks, a peak on chromosome 13 to be protein-specific, and peaks on chromosome 15 and 20 to be pleiotropic peaks. It should be noted that trait-specific peaks on chromosome 8 and 13 are novel peaks, which were missing in a recently reported mvLMM-GWAS analysis for seed oil and protein[24]. Because the mvLMM-GWAS results looked like the overlapping of the two univariate LMM-GWAS results and our objective for GWAS is to examine the possibility of improving GWAS resolution from the imputation of genome resequencing panel data into an existing GWAS, we focused on results from mvLMM results for a comparison between the existing GWAS and imputed GWAS.

**Imputation of existing SoySNP50K array genotyping data with genome sequencing panel data.** We imputed 10.5 million SNPs into 36,647 SNPs from SoySNP50K data of 8,844 soybean accessions. In the final imputed data, 1% (107,902) of SNPs from our genome resequencing panel were lost and ~15% (5,501) of SNPs from the SoySNP50K data were lost. The lost SNPs from our genome resequencing panel were SNPs from unanchored scaffolds and chloroplast. The lost 5,501 SNPs from the SNP array are those that are not overlapping with positions in the set of the 10.5 million SNPs. From the 62.9 million raw SNPs, we found 4,753 of the 5,501 SNP positions, however 1,541 positions (32.4%) of 4,753 were overlapping deletions and 1,385 (29.1%) of them were multi-allelic sites. The results suggested that probes of the lost 5,501 SNPs were likely designed from highly variable, low-quality SNP loci that were called from low-depth resequencing data obtained from eight soybean genotypes[20]. Imputation accuracy was evaluated by comparison between imputed genotypes and true genotypes in 97 soybean samples that were shared between our genome resequencing panel and the 8,844 oil-protein samples. The average concordance rate between imputed genotypes and resequencing panel data of the 97 soybean samples was 96.5%. When six potentially mislabeled accessions (PI 407736, PI 84669N, PI 567587A, PI 96354, PI 603910A, and PI 398874) during maintenance at germplasm banks based on PCA plots that showed different relative coordinates between our genome resequencing panel population and the 8,844 oil-protein samples (Supplementary Figs. 5 and 11) were excluded, the average concordance rate slightly increased to 97.0%. The estimated concordance rate is comparable to those of maize (97.2%)[25] and human (90 – 95%)[26].

### References


1. Jeong, N., Moon, J.K., Kim, H.S., Kim, C.G. & Jeong, S.C. Fine genetic mapping of the genomic region controlling leaflet shape and number of seeds per pod in the soybean. *Theor Appl Genet* **122**, 865-74 (2011).
2. Schmutz, J. *et al.* Genome sequence of the palaeopolyploid soybean. *Nature* **463**, 178-83 (2010).
3. DePristo, M.A. *et al.* A framework for variation discovery and genotyping using next-generation DNA sequencing data. *Nat Genet* **43**, 491-8 (2011).
4. Lu, B.R. Conserving biodiversity of soybean gene pool in the biotechnology era. *Plant Spec Biol* **19**, 115-25 (2004).

5. Carlson, J.B. & Lersten, N.R. Reproductive morphology. in *Soybeans: Improvement, production, and uses, 3rd edn* (eds. Boerma, H.R. & Specht, J.E.) 59-95 (ASA, CSSA, and SSSA, Madison, 2004).
6. Ray, J.D., Kilen, T.C., Abel, C.A. & Paris, R.L. Soybean natural cross-pollination rates under field conditions. *Environ Biosafety Res* **2**, 133-8 (2003).
7. Fujita, R., Ohara, M., Okazaki, K. & Shimamoto, Y. The extent of natural cross-pollination in wild soybean (*Glycine soja*). *J Hered* **88**, 124-28 (1997).
8. Patterson, N., Price, A.L. & Reich, D. Population structure and eigenanalysis. *PLoS Genet* **2**, e190 (2006).
9. Raj, A., Stephens, M. & Pritchard, J.K. fastSTRUCTURE: variational inference of population structure in large SNP data sets. *Genetics* **197**, 573-89 (2014).
10. Jeong, S.C. *et al.* Genetic diversity patterns and domestication origin of soybean. *Theor Appl Genet* **132**, 1179-1193 (2019).
11. Lee, Y.G. *et al.* Development, validation and genetic analysis of a large soybean SNP genotyping array. *Plant J* **81**, 625-36 (2015).
12. Dong, Z., Alexander, M. & Chuck, G. Understanding grass domestication through maize mutants. *Trends Genet* **35**, 118-128 (2019).
13. Zhang, D. *et al.* Elevation of soybean seed oil content through selection for seed coat shininess. *Nat Plants* **4**, 30-35 (2018).
14. Tian, Z. *et al.* Artificial selection for determinate growth habit in soybean. *Proc Natl Acad Sci U S A* **107**, 8563-8 (2010).
15. Sun, L. *et al.* *GmHs1-1*, encoding a calcineurin-like protein, controls hard-seededness in soybean. *Nat Genet* **47**, 939-43 (2015).
16. Dong, Y. *et al.* Pod shattering resistance associated with domestication is mediated by a NAC gene in soybean. *Nat Commun* **5**, 3352 (2014).
17. Swarm, S.A. *et al.* Genetic dissection of domestication-related traits in soybean through genotyping-by-sequencing of two interspecific mapping populations. *Theor Appl Genet* **132**, 1195-1209 (2019).
18. Funatsuki, H. *et al.* Molecular basis of a shattering resistance boosting global dissemination of soybean. *Proc Natl Acad Sci U S A* **111**, 17797-802 (2014).
19. Bandillo, N. *et al.* A population structure and genome-wide association analysis on the USDA soybean germplasm collection. *The Plant Genome* **8**(2015).
20. Song, Q. *et al.* Development and evaluation of SoySNP50K, a high-density genotyping array for soybean. *PLoS One* **8**, e54985 (2013).
21. Anderson, C.A. *et al.* Data quality control in genetic case-control association studies. *Nat Protoc* **5**, 1564-73 (2010).
22. Patil, G. *et al.* Dissecting genomic hotspots underlying seed protein, oil, and sucrose content in an interspecific mapping population of soybean using high-density linkage mapping. *Plant Biotechnol J* **16**, 1939-1953 (2018).
23. Burton, J.W. Quantitative genetics: Results relevant to soybean breeding. in *Soybeans: Improvement, production and uses, 2nd edn* (ed. Wilcox, J.R.) 211–247 (ASA, CSSA, and SSSA, Madison, 1987).
24. Lee, S. *et al.* Genome-wide association study of seed protein, oil and amino acid contents in soybean from maturity groups I to IV. *Theor Appl Genet* **132**, 1639-1659 (2019).
25. Wang, D.R. *et al.* An imputation platform to enhance integration of rice genetic resources. *Nat Commun* **9**, 3519 (2018).
26. Genomes Project, C. *et al.* An integrated map of genetic variation from 1,092 human genomes. *Nature* **491**, 56-65 (2012).

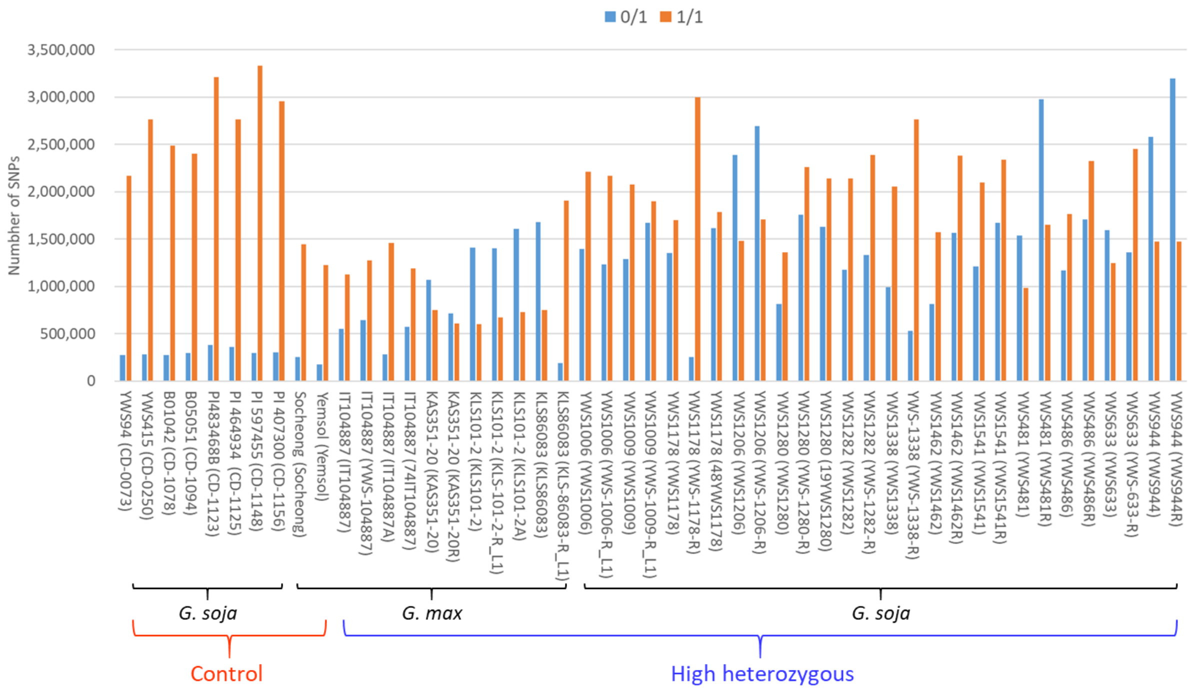


**Supplementary Figure 1. Distribution of heterozygous and homozygous SNP genotypes called from a set of soybean samples consisting of 27 soybean accessions and 49 samples.** Number of raw SNPs obtained from the GenotypeGVCF step of the Genome Analysis Toolkit (GATK) Best Practices are shown. Reference allele is indicated by 0, non-reference alleles is 1. Heterozygous SNPs (blue bars) and homozygous SNPs (red bars) are 0/1 and 1/1, respectively. Memberships to domesticated (*Glycine max*) and wild (*Glycine soja*) soybean are indicated under soybean accession and sample (in parenthesis) names. Control are those samples that have shown low heterozygosity in repeated SNP calling attempts and high heterozygous are those samples that have shown more heterozygous SNPs in the first analyzed samples shown leftmost of the list of each of soybean accessions. Genome resequencing of all 17 high heterozygous samples were repeated using new libraries prepared from filial bulks of the first sequenced samples.  In case of IT104887, additional resequencing data were obtained from a filial single plant (IT104887A) and using the same DNA (repeated sample, 74IT104887) as the first sample. In case of KLS101-2, additional resequencing data were obtained from a filial single plant (KLS101-2A). In cases of YWS1178 and YWS1280, additional resequencing data were obtained using the same DNAs (48YWS1178 and 19YWS1280, respectively) as the first samples.

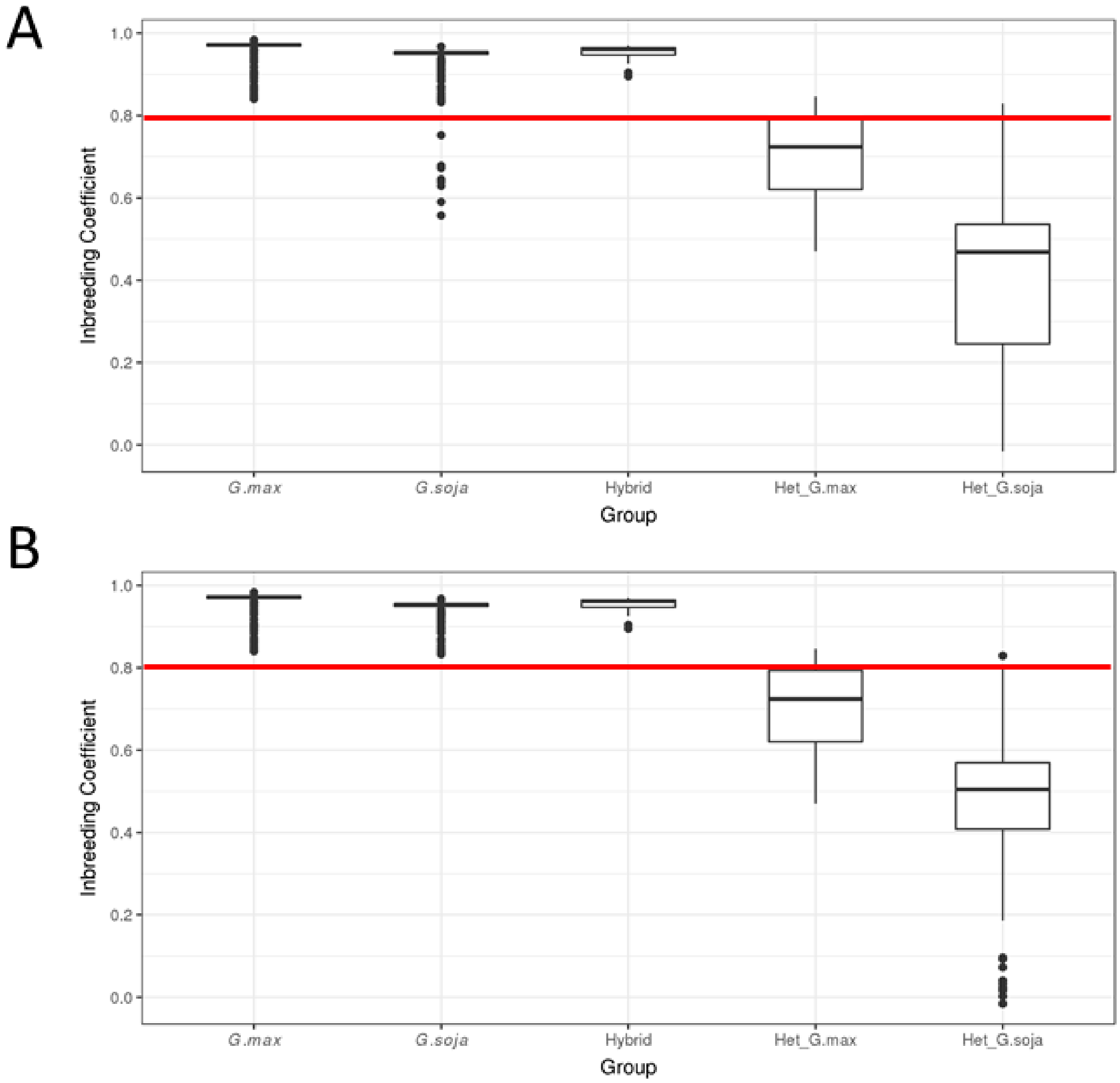


**Supplementary Figure 2. Distribution of inbreeding coefficients within subpopulations of a soybean population of 855 samples.** (**A)** Distribution of individual inbreeding coefficients in subpopulations. (**B)** Distribution of individual inbreeding coefficients in subpopulations after transferring eight wild accessions with inbreeding coefficient per individual of less than 0.8 from *G. soja* to high heterozygous *G. soja* subpopulation eight wild accessions. Subpopulations: *G. max*, domesticated; G. soja, wild; Hybrid, *G. max* x *G. soja*; Het_G.max, high heterozygous domesticated; and Het_G.soja, high heterozygous wild soybean subpopulation.

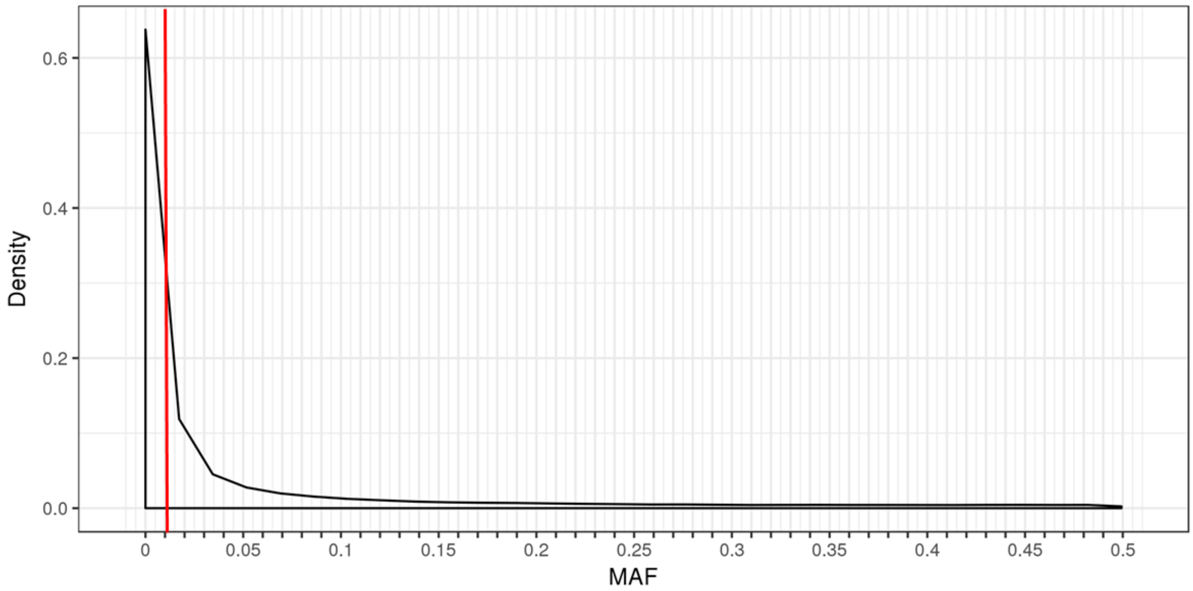


**Supplementary Figure** 3. **Distribution of minor allele frequency (MAF) for SNPs. A red line indicates 1% MAF.**

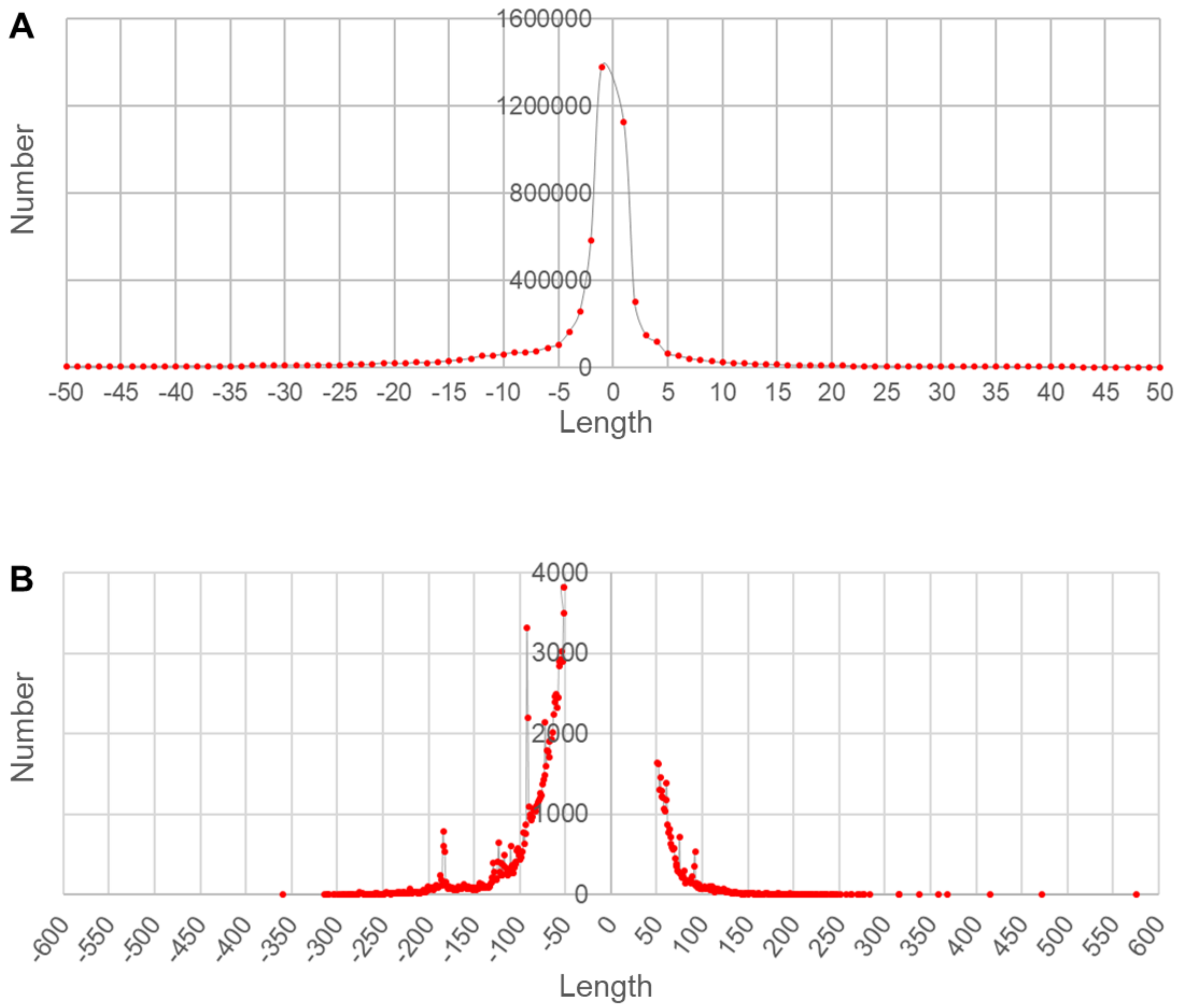


**Supplementary Figure 4. Distribution of sizes of insertion/deletion (indel) variants and structural variants (SV).** (**A**) Indel size distribution. (**B**) SV size distribution.

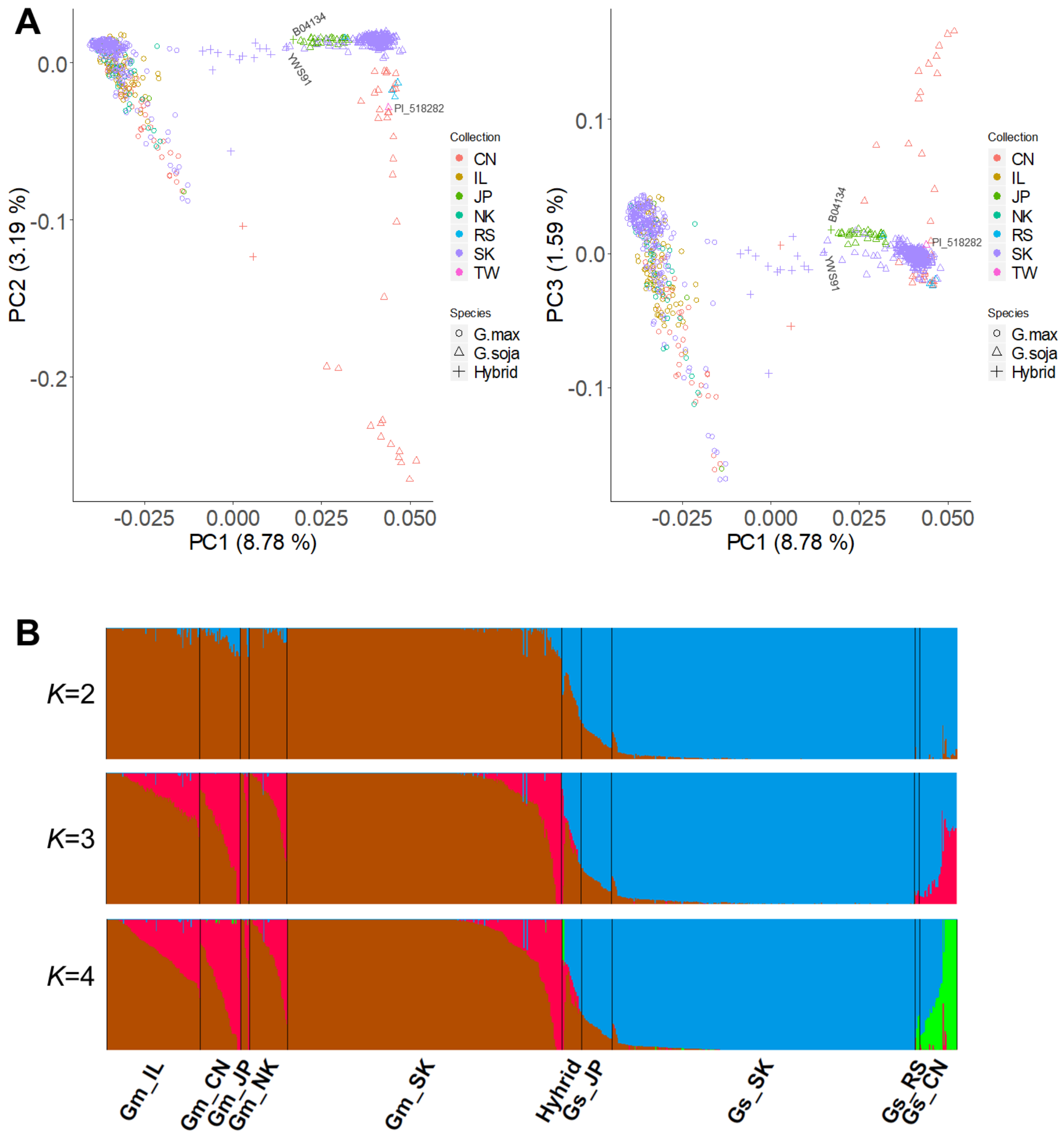


**Supplementary Figure 5. Population structure of the 781 haplotype soybean accession set.** (**A**) Principal components (PC) of SNP variation. The plots show the first three principal components. (**B**) fastSTRUCTURE plots. The accessions were divided into three groups: *Glycine max*, *G. soja*, and hybrids. The accessions were further indicated by the countries of collection or improvement status of the soybean accessions represented by two-letter codes—CN, China; IP, improved breeding line; JP, Japan; North Korea; RS, Russia; SK, South Korea; and TW, Taiwan. Two hybrid accessions JPN38 and YWS91 were located near G. soja accessions YWS204 and YWS1034 at the margin of a *G. soja* subpopulation. An only accession PI 549046 (G. soja) from Taiwan was labeled. All accessions from NK were *G. max*. In **B**, Gm and Gs are *G. max* and *G. soja*, respectively, and a *G. soja* accession from Taiwan was included in the Cs_CN group based on the PC analysis.

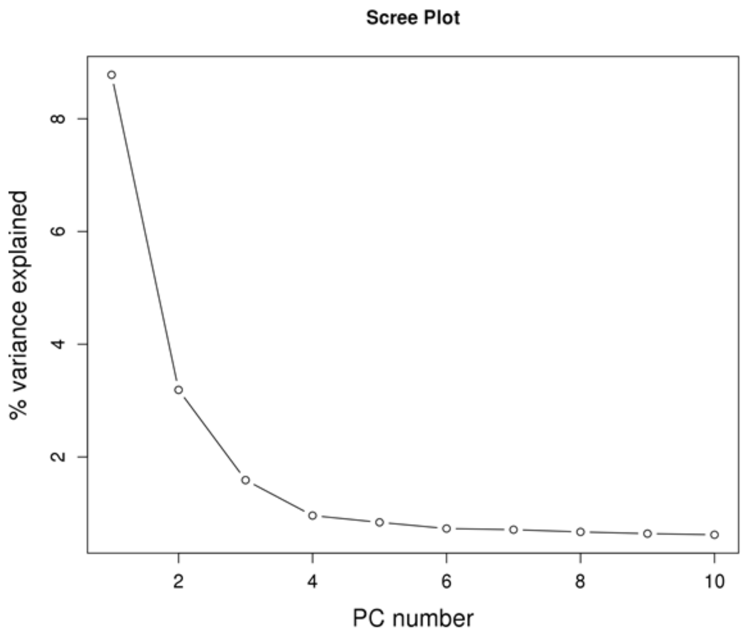


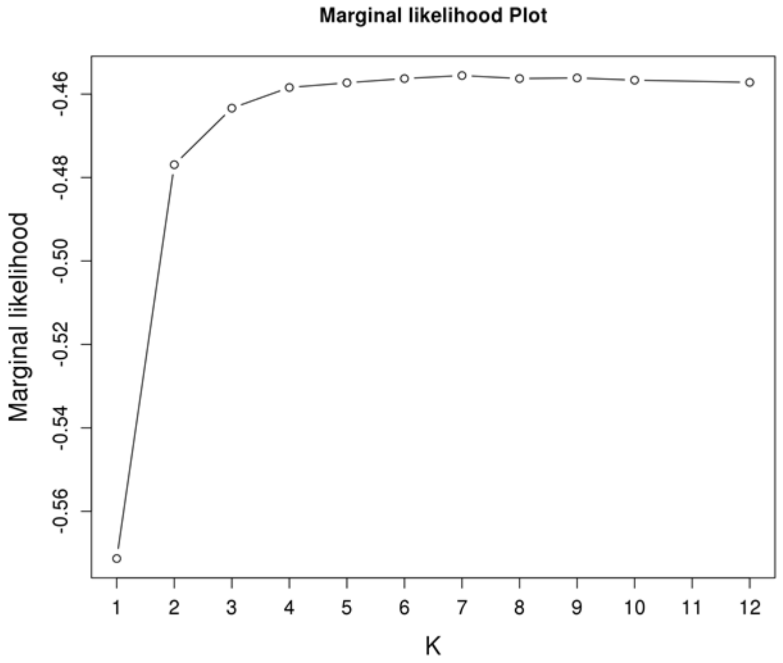


**Supplementary Figure 6. Scree plot and marginal likelihood plot from population structure analyses of the 781 haplotype soybean accession.** Scree plot shows the PC number and their contribution to variance from principal component analysis. Marginal likelihood plot shows the model complexity (*K*) and marginal likelihood from fastSTRUCTURE.

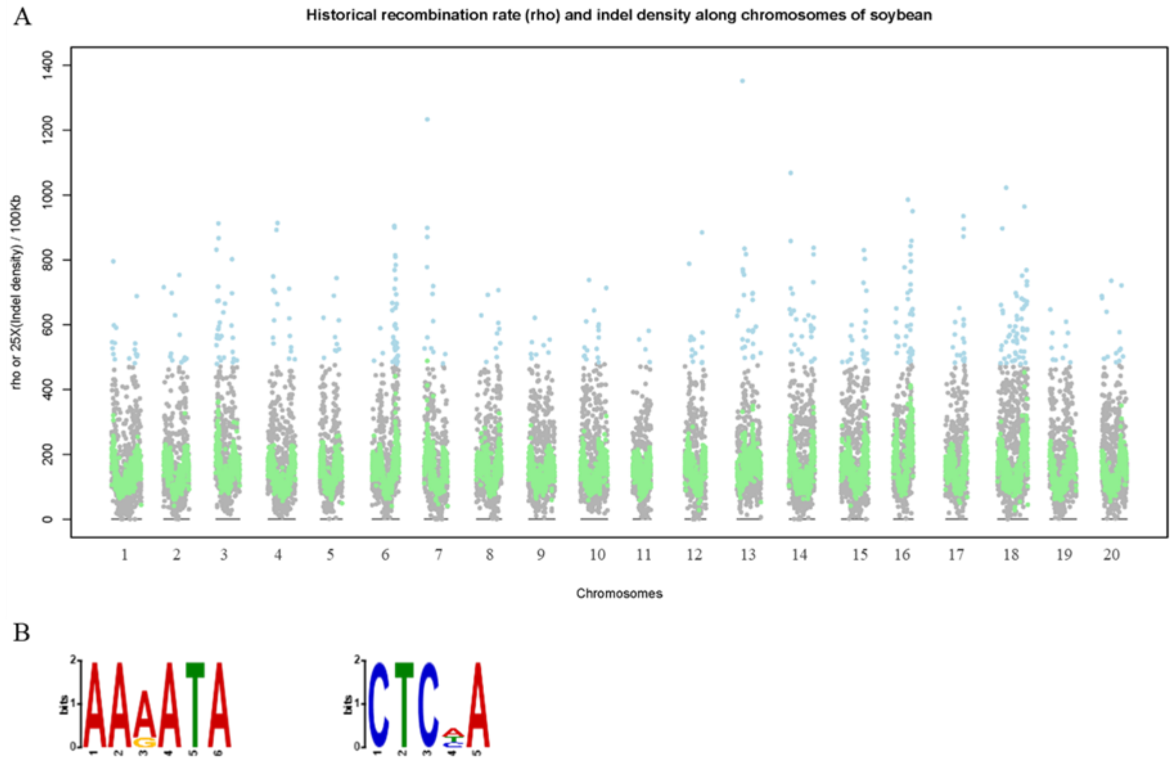


**Supplementary Figure 7. Relationship between historical recombination rate and enriched sequence motifs.** (**A**) The distribution of historical recombination rate (rho: gray dots and outliers were highlighted by blue color) and indel density (green dots) along the chromosomes of soybean. To visualize two values in a similar scale, indel density values were multiplied by 25. (**B**) Sequence logos of suggested short motifs from DREME analysis with selected 66,549 indel sequences.

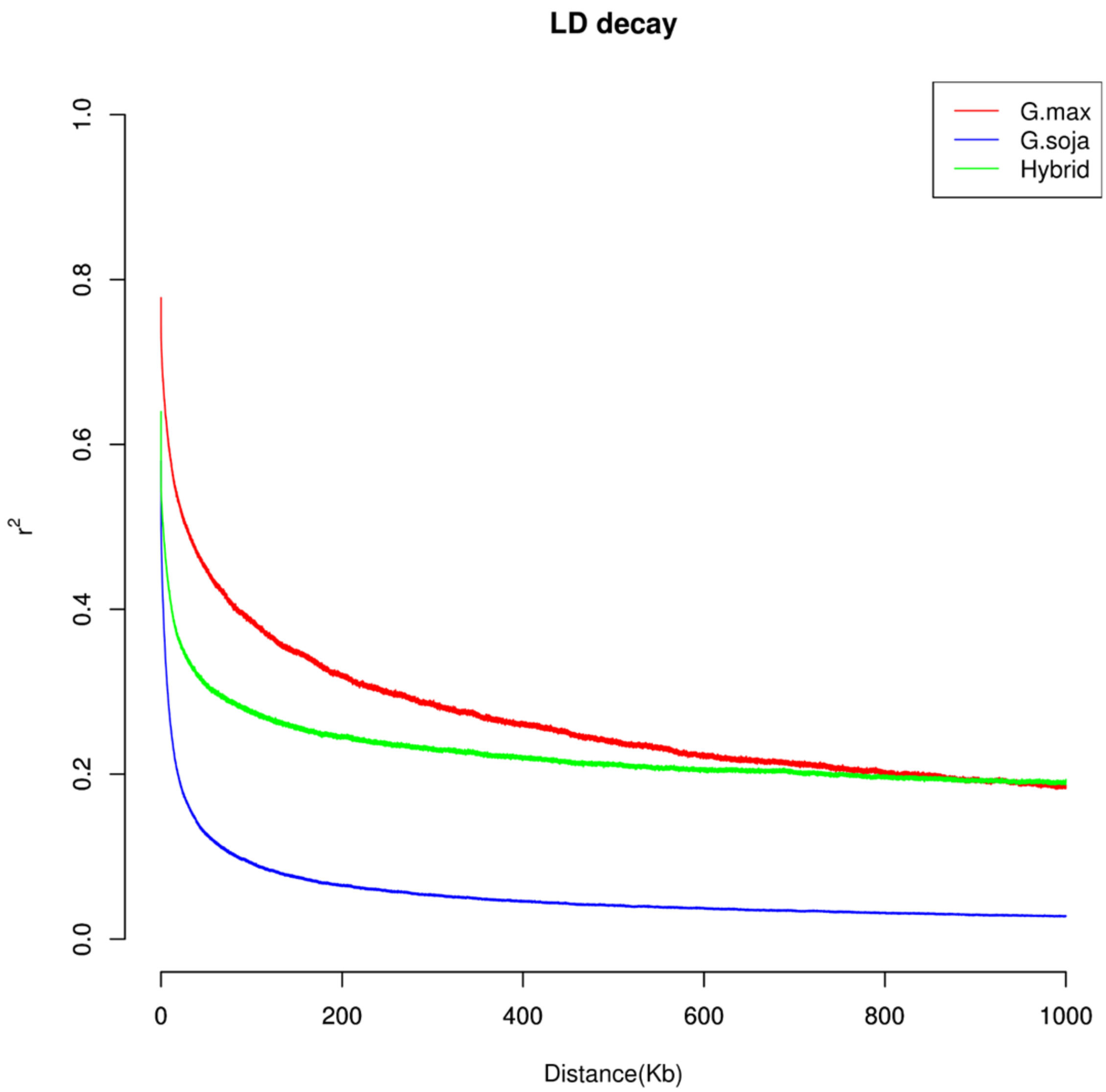


**Supplementary Figure 8. Linkage disequilibrium (LD) decay in soybean groups.** Decay rates of LD in domesticated (*Glycine max*), wild (*Glycine soja*), and their natural hybrids were determined by the squared correlations of allele frequencies ($r^2$) against physical distance (kb) between polymorphic SNP loci.

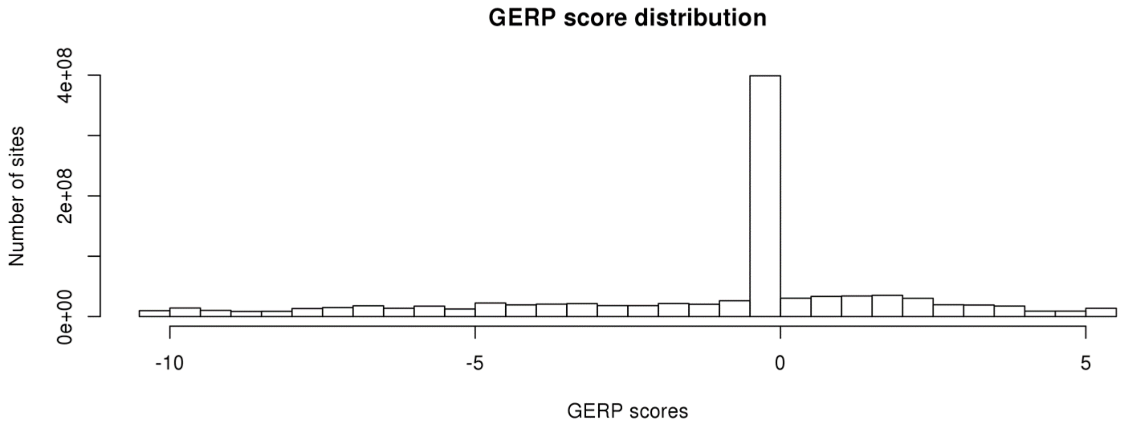


**Supplementary Figure 9. Distribution of GERP scores for soybean genome.**

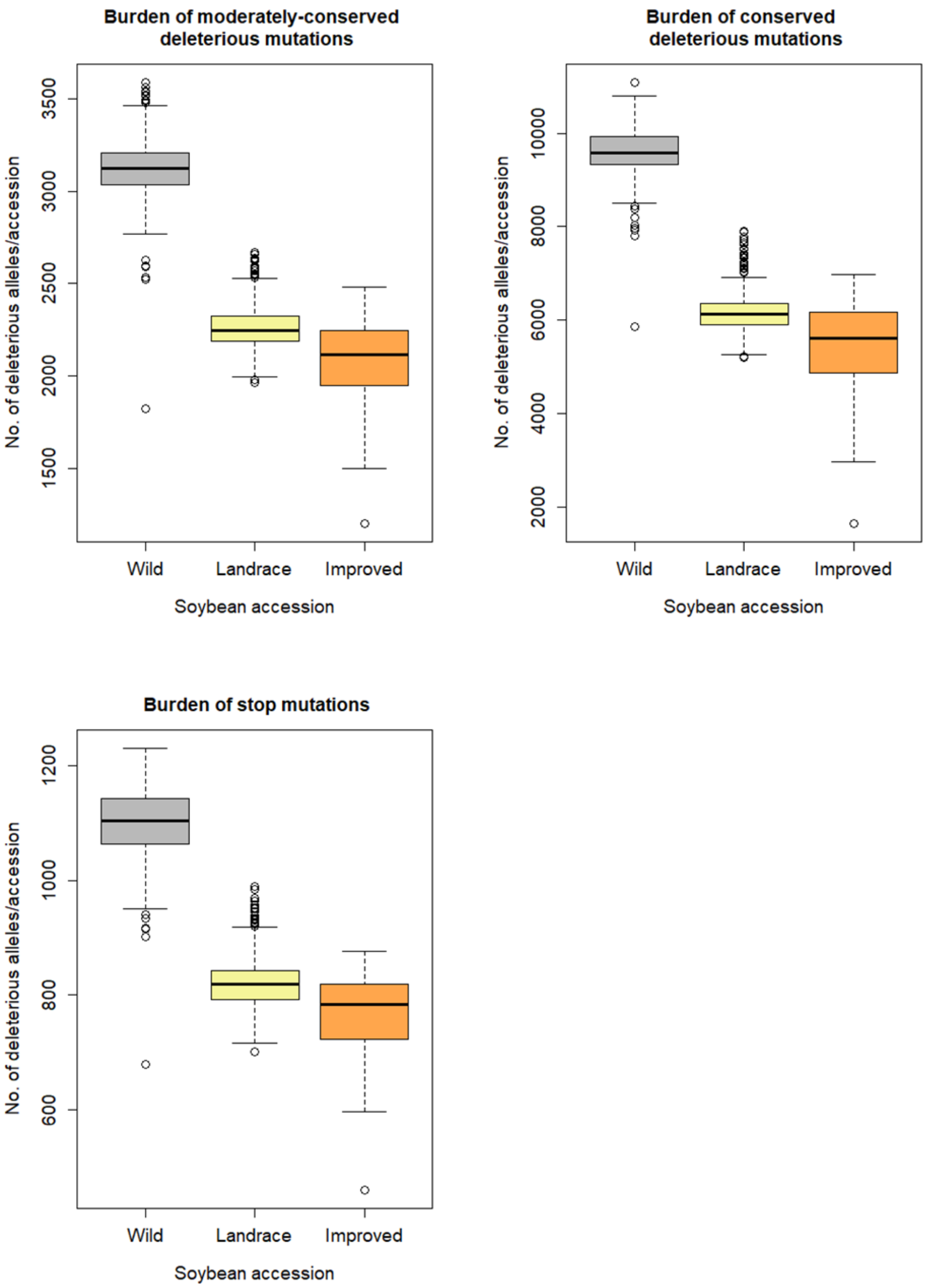


**Supplementary Figure 10. Box-and-whisker plot distributions of mutation burden in domesticated (*Glycine max*, landrace cultivars = 332 and improved lines = 86) and wild (*Glycine soja*, n = 345) soybean populations.** Distribution of each of three deleterious mutation categories is shown: conserved deleterious (nonsynonymous, SIFT $< 0.05$, GERP $\geq$ 2,), moderately-conserved deleterious (nonsynonymous, SIFT $< 0.05$, $0 <$ GERP $< 2$,), and stop mutations. The subgroups in each of plots are significantly different between one another with $P < 0.09e\text{-}5$ in Tukey multiple comparison tests.

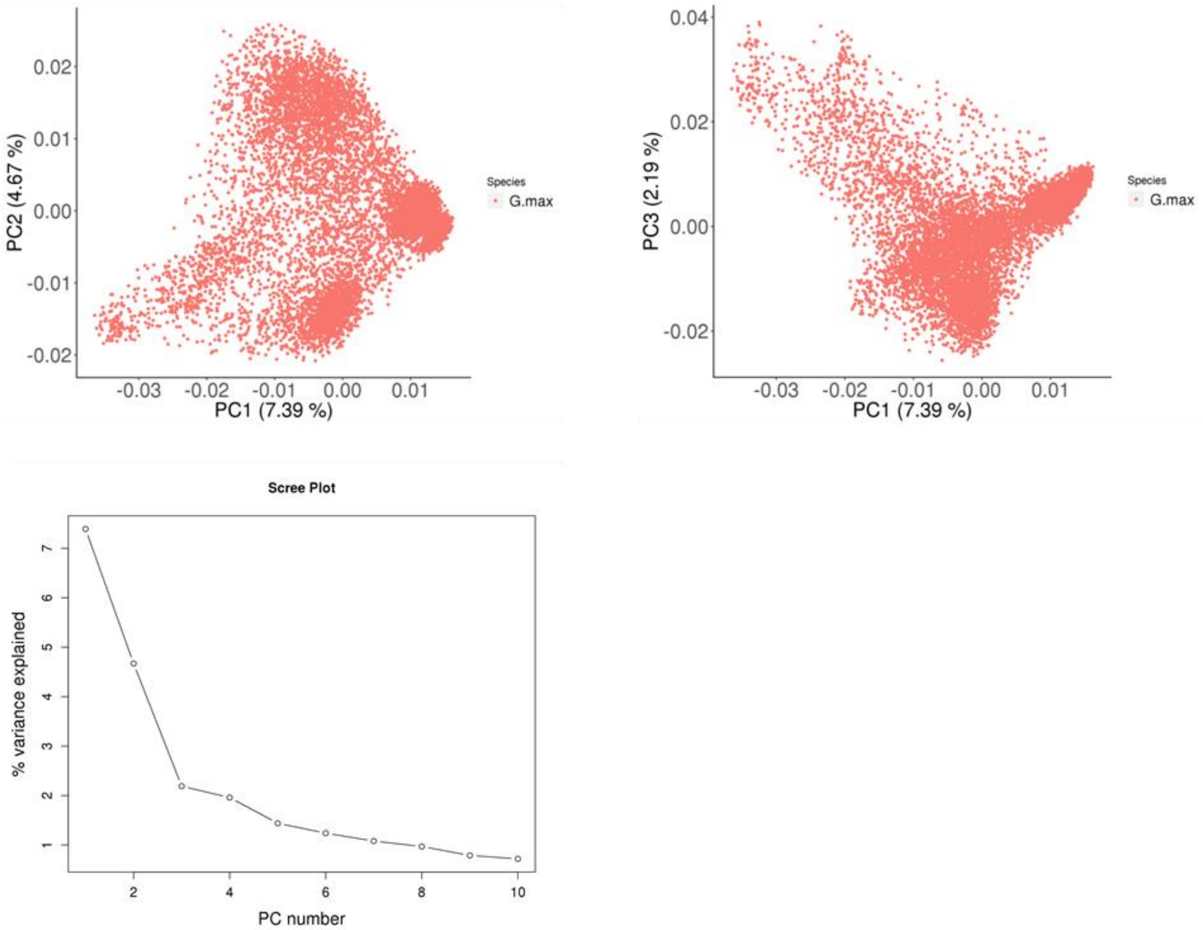


**Supplementary Figure 11. Principal component analysis of 8,844 non-redundant domesticated soybean (*G. max*) accessions.** The plots (top) show the first three principal components. Scree plot (bottom) of the PC number and their contribution to variance from principal component analysis.

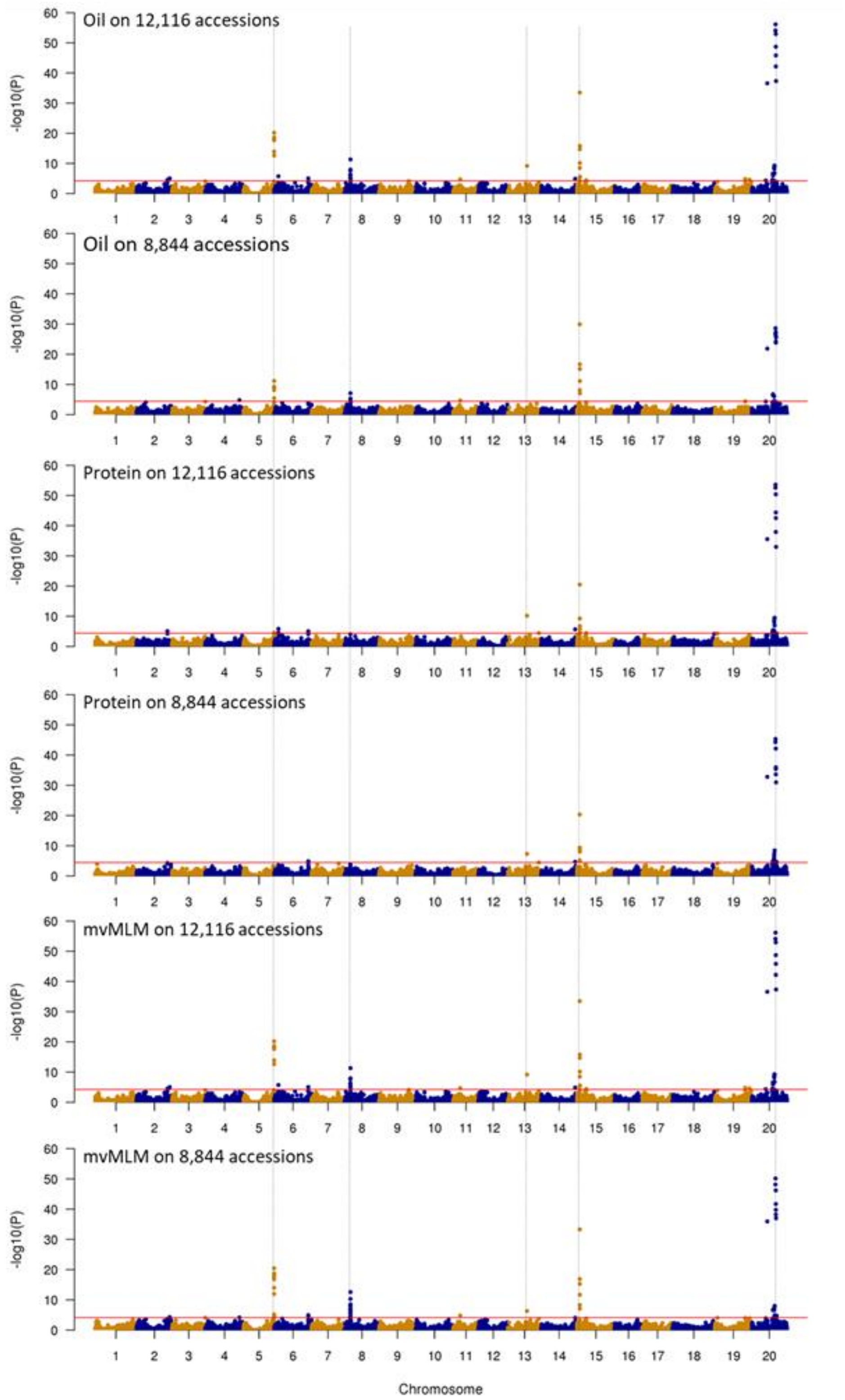


**Supplementary Figure 12. Comparison of genome-wide association studies for seed oil and protein contents in both the 12,116 and 8,844 soybean accession sets using both univariate LMM and multivariate LMM (mvLMM) models.** Horizontal line represent 5% significance thresholds corrected for multiple testing using Benjamini-Hochberg that ranged from 4.15 to 4.44. Chromosomal regions of five major peaks are indicated by dashed vertical line for comparison.

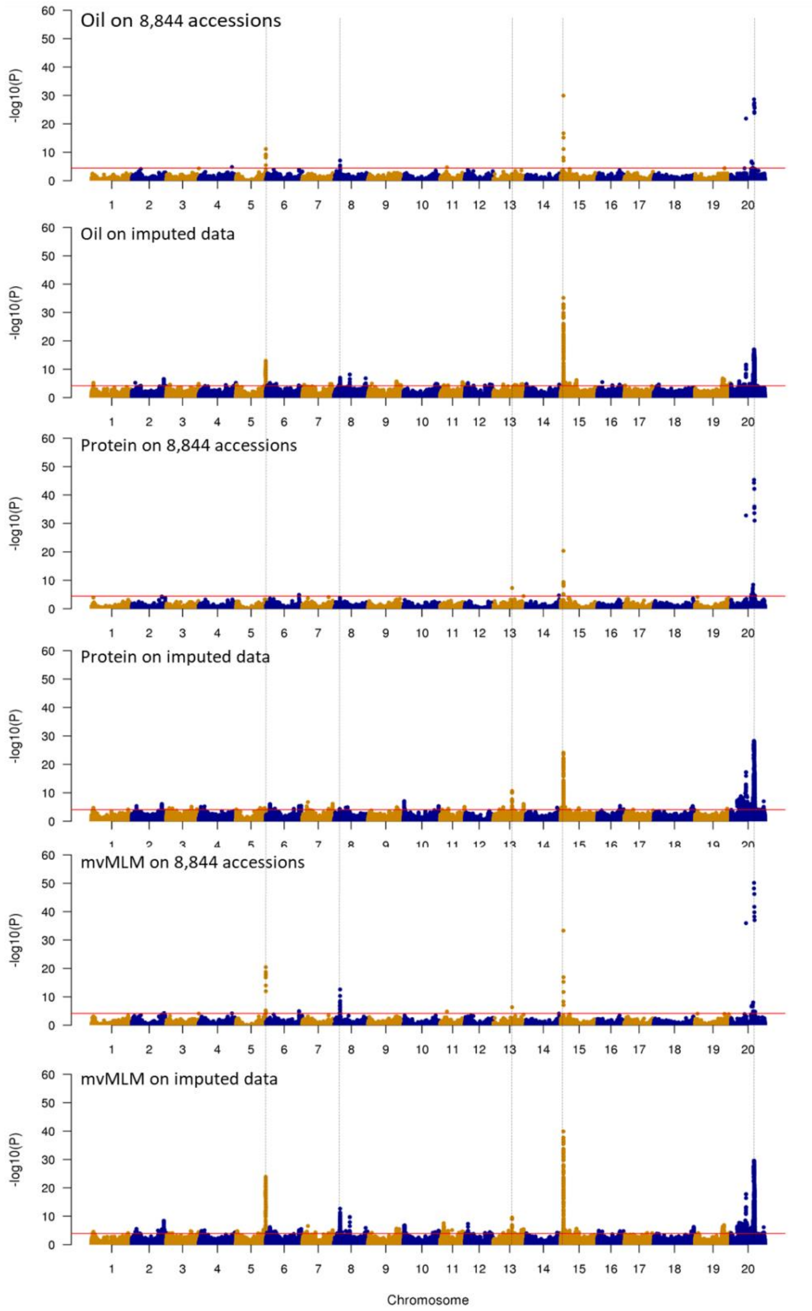


**Supplementary Figure 13. Comparison of genome-wide association scans for variants associated with seed oil and protein using SoySNP50K and imputed genotype data in soybean.** Six different trait-based and model-based Manhattan plots represent -log 10 (p-value) for SNPs distributed across all 20 chromosomes of soybean. Y-axis:-log 10 (p-value) and x-axis: soybean chromosomes. The red lines stand as FDR thresholds that ranged from 3.925 to 4.440. Univariate linear mixed model and multivariate linear mixed model (mvLMM) were used.

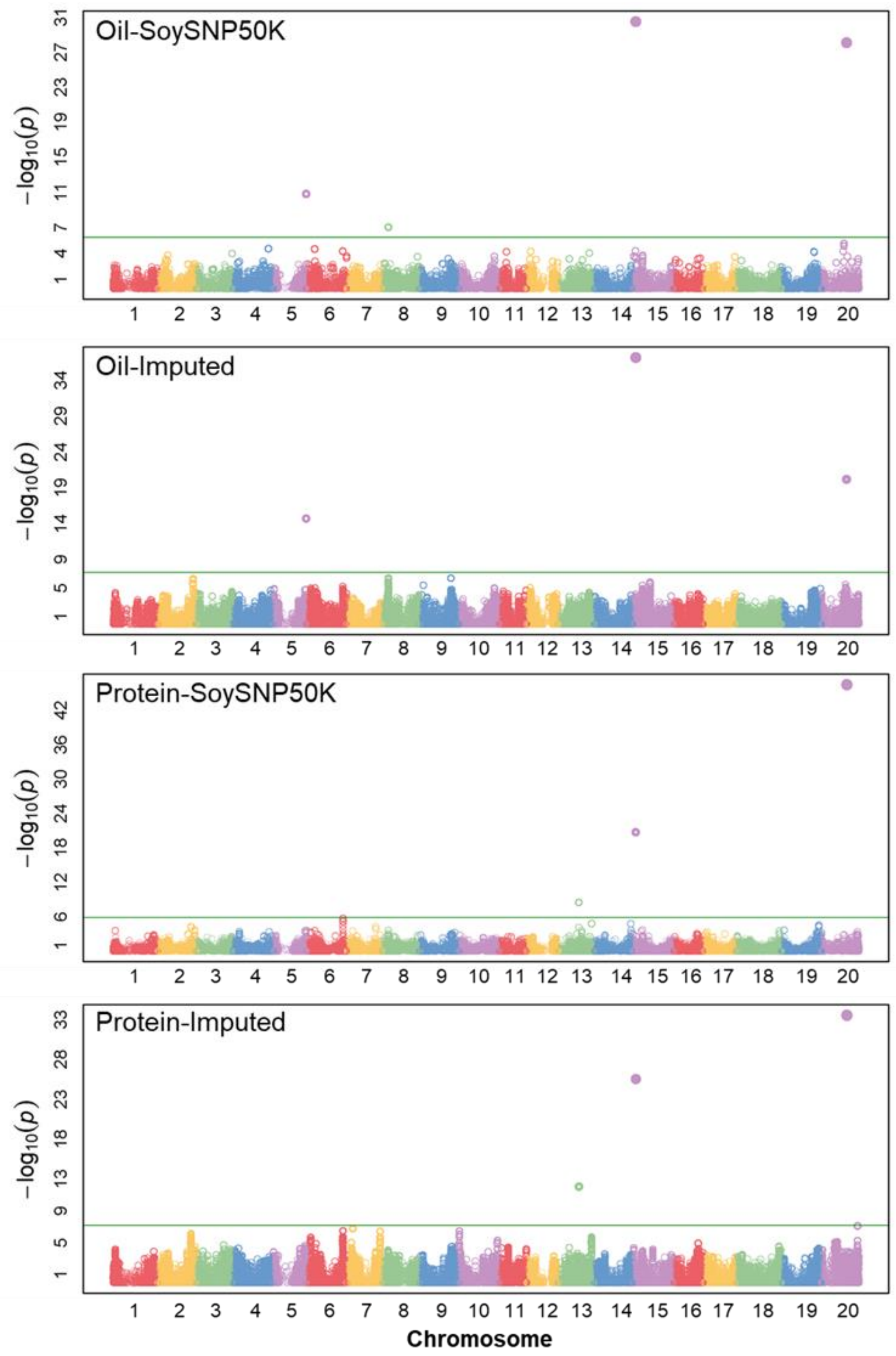


**Supplementary Figure 14. Comparison of genome-wide association studies using multi-locus mixed model (MLMM) for variants associated with oil and protein using SoySNP50K and imputed genotype data in 8,844 soybean accessions.** The green line stands as FDR thresholds that ranged from 5.17 to 6.74.

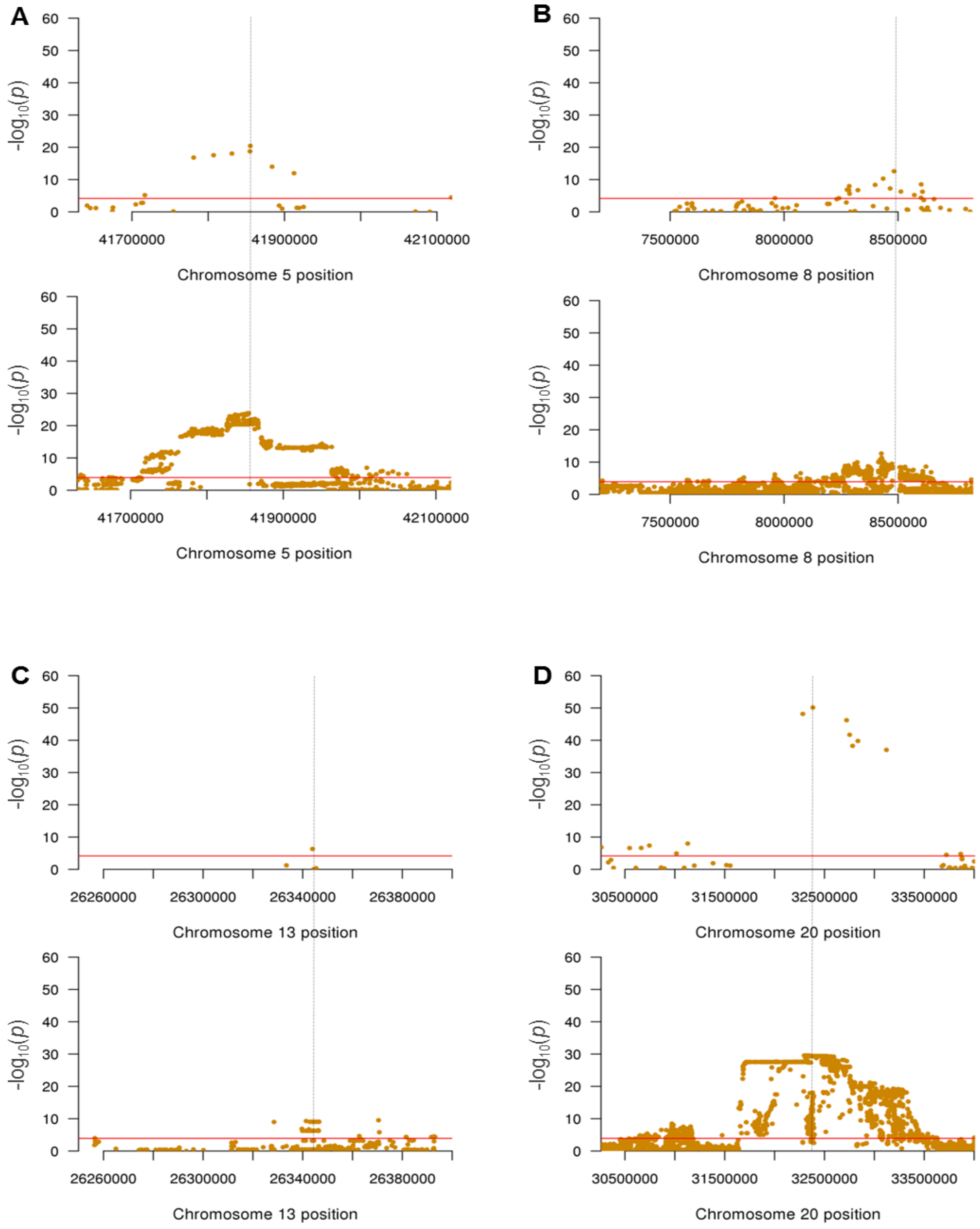


**Supplementary Figure 15. Comparison of mvMLM-based GWAS results using unimputed (SoySNP50K) and imputed 10.4 million SNP data at major peaks on chromosomes.** (**A**) A major peak on chromosome 5. The most significant SNP in the GWAS on unimputed data is indicated by dashed vertical lines for comparison. (**B**) A major peak on chromosome 8. (**C**) A major peak on chromosome 13. (**D**) A major peak on chromosome 20.

### Supplementary Table 1. List of soybean accessions sequences

| Number in haplotype map panel | SampleID | Type (landrace or improved line for G. max) | Name or accession code (IT, PI, and B number) used in this study | Other name or accession code | Collected country (*G. max* and *G. soja*) and province (*G. soja*) | Total raw sequences (bp) | Mean depth of non-duplicates | Genome coverage above 1 reads (%) | Genome coverage above 5 reads (%) |
|---|---|---|---|---|---|---|---|---|---|
| 1 | Ajuggali | Landrace | Ajuggali | IT180576 | South Korea | 26,194,459,240 | 25.35 | 98.5 | 97.3 |
| 2 | Anpyeong | Improved | Anpyeong | | South Korea | 16,409,936,476 | 15.61 | 96.7 | 90.1 |
| 3 | Babmit | Landrace | Babmit | IT175966 | South Korea | 18,787,686,666 | 17.39 | 98.5 | 97 |
| 4 | Back_Tac | Landrace | Back Tac | PI 567273 A | South Korea | 16,286,354,452 | 15.83 | 98.4 | 96.5 |
| 5 | Baegbong | Landrace | Baegbong | IT134332 | South Korea | 16,081,380,710 | 15.49 | 98.2 | 95.7 |
| 6 | Baekjung_42 | Landrace | Baekjung #42 | IT229365 | South Korea | 21,166,880,718 | 20.9 | 98.1 | 96.4 |
| 7 | Baekmo_9 | Landrace | Baekmo #9 | IT230778 | South Korea | 17,169,009,550 | 16.82 | 98.3 | 96.3 |
| 8 | Bancheongdu | Landrace | Bancheongdu | IT230769 | South Korea | 17,201,297,578 | 16.39 | 96.7 | 90.6 |
| 9 | Bangsa | Improved | Bangsa | | South Korea | 16,777,601,846 | 15.54 | 97.1 | 90.2 |
| 10 | Blackhawk | Improved | Blackhawk | PI 548516 | United States | 15,715,503,482 | 15.05 | 96.9 | 89.9 |
| 11 | Boone | Improved | Boone | PI 548309 | China | 22,868,391,300 | 22.19 | 98.2 | 96.7 |
| 12 | Boseok | Improved | Bosug | | South Korea | 16,025,387,796 | 15.27 | 96.8 | 90.5 |
| 13 | Browngilgun | Landrace | Browngilgun | PI 612611 | South Korea | 22,342,411,792 | 20.05 | 98.1 | 96.6 |
| 14 | Bukwang | Improved | Bukwang | | South Korea | 16,585,723,428 | 15.79 | 96.6 | 90.4 |
| 15 | Cai_dou | Landrace | Cai dou | PI 567297 | China | 17,326,340,074 | 16.81 | 97.8 | 95.5 |
| 16 | Cheonangun | Landrace | Cheonangun | IT162681 | South Korea | 21,042,587,786 | 20.33 | 98.2 | 96.7 |
| 17 | Cheongja | Improved | Cheongja | | South Korea | 16,762,904,110 | 15.09 | 97.2 | 89.8 |
| 18 | Cheongja3 | Improved | Cheongja3 | | South Korea | 16,984,628,282 | 15.34 | 96.9 | 89.4 |
| 19 | Cheongsong-1 | Landrace | Cheongsong sujib-1 | IT221947 | South Korea | 18,892,831,590 | 17.1 | 98.2 | 96.4 |
| 20 | Cheongtae | Landrace | Cheongtae | IT175994 | South Korea | 28,786,142,908 | 28.44 | 98.5 | 97.4 |
| 21 | Chonggok | Landrace | Chonggok | IT230808 | South Korea | 20,919,008,178 | 19.92 | 98.2 | 96.4 |
| 22 | Cin | Landrace | Cin | PI 603910 A | North Korea | 20,651,905,788 | 18.92 | 98.2 | 96.7 |
| 23 | Clark | Improved | Clark | PI 548533 | United States | 49,478,480,758 | 43.24 | 99.2 | 98.8 |

| | | | | | | | | | |
|---|---|---|---|---|---|---|---|---|---|
| 24 | CNS | Landrace | CNS | PI 548445 | China | 21,672,975,338 | 21.26 | 98.2 | 96.7 |
| 25 | CS_00829 | Landrace | CS 00829 | IT221942 | South Korea | 32,227,154,766 | 30.7 | 98.3 | 97.1 |
| 26 | CS_01964 | Landrace | CS 01964 | IT229073 | South Korea | 24,182,085,562 | 23.68 | 98.4 | 97 |
| 27 | CS_02038 | Landrace | CS 02038 | IT237763 | South Korea | 31,091,710,266 | 28.97 | 98.5 | 97.4 |
| 28 | Daechu | Landrace | Daechu | IT224516 | South Korea | 27,152,216,906 | 25.9 | 98.4 | 97.2 |
| 29 | Daeheug | Improved | Daeheug | | South Korea | 15,661,014,528 | 15.03 | 96.8 | 89.9 |
| 30 | Daepung | Improved | Daepung | | South Korea | 15,929,011,140 | 14.99 | 97 | 90.1 |
| 31 | Daewon | Improved | Daewon | | South Korea | 17,147,019,118 | 16.04 | 97 | 90.1 |
| 32 | Daeyang | Improved | Daeyang | | South Korea | 16,288,560,562 | 15.65 | 96.7 | 90 |
| 33 | Danbaeg | Improved | Danbaek | | South Korea | 17,193,738,216 | 15.81 | 97.1 | 90.5 |
| 34 | Danmi2 | Improved | Danmi2 | | South Korea | 15,504,692,080 | 14.87 | 96.5 | 89.3 |
| 35 | Dongsan-133 | Improved | Dongsan133 | IT154786 | Japan | 16,978,662,876 | 15.68 | 98.2 | 96.1 |
| 36 | Doremi | Improved | Doremi | | South Korea | 15,982,013,348 | 15.31 | 96.7 | 90.5 |
| 37 | Dowling | Improved | Dowling | PI 548663 | United States | 33,340,483,806 | 31.64 | 98.4 | 97.4 |
| 38 | Fen_dou_16 | Landrace | Fen dou 16 | PI 574476 A | China | 20,522,423,892 | 19.26 | 98 | 96 |
| 39 | Fiskeby | Improved | Fiskeby 840-7-3 | PI 438477 | Sweden | 15,543,431,130 | 14.83 | 96.5 | 88.8 |
| 40 | Fu_yang | Landrace | Fu yang (30) | PI 567709 | China | 45,474,782,198 | 41.48 | 98.4 | 97.3 |
| 41 | Galchae | Improved | Galchae | | South Korea | 15,420,136,610 | 14.77 | 96.6 | 90 |
| 42 | Galmi | Landrace | Galmi | | South Korea | 17,319,018,084 | 16.96 | 98.3 | 96.3 |
| 43 | Gangwon-72 | Landrace | Kangwon72 | IT186084 | South Korea | 19,028,727,060 | 17.32 | 98.4 | 96.9 |
| 44 | Geomeon | Landrace | Geomeun | IT109118 | South Korea | 15,174,804,192 | 14.09 | 96.7 | 90.2 |
| 45 | Geomeunbak | Landrace | Geomeunbak | IT178683 | South Korea | 16,946,856,538 | 15.7 | 98.4 | 96.6 |
| 46 | Geomjeong-1 | Landrace | Geomjeong-1 | IT177394 | South Korea | 15,397,899,444 | 14.65 | 96.6 | 89.3 |
| 47 | Geomjeong-2 | Landrace | Geomjeong-2 | IT177645 | South Korea | 15,406,115,354 | 14.62 | 96.8 | 90.7 |
| 48 | Geomjeong-3 | Landrace | Geomjeong-3 | IT177581 | South Korea | 15,553,814,494 | 14.83 | 96.6 | 90.1 |
| 49 | Geomjeongol | Landrace | Geomjeongol | A08 | South Korea | 16,177,266,556 | 15.95 | 98.3 | 95.5 |
| 50 | Giant | Landrace | Giant | PI 157422 | South Korea | 23,514,528,152 | 22.78 | 98.1 | 96.6 |
| 51 | GL_2624_96 | Landrace | GL 2624 /96 | PI 603156 | North Korea | 29,620,266,606 | 29.14 | 98.4 | 97.2 |
| 52 | GL_2626_96 | Landrace | GL 2626 /96 | PI 603158 | North Korea | 18,731,710,966 | 17.91 | 97.9 | 95.8 |
| 53 | GL_2631_96 | Landrace | GL 2631 /96 | PI 603162 | North Korea | 28,582,406,762 | 28.06 | 98 | 96.6 |

| | | | | | | | | | |
|---|---|---|---|---|---|---|---|---|---|
| 54 | GL_2684_95 | Landrace | GL 2684 /95 | PI 603171 | North Korea | 18,571,914,008 | 18.07 | 98.2 | 96.4 |
| 55 | GL_2687_A | Landrace | GL 2687 A | IT242452 | North Korea | 21,976,716,670 | 20.1 | 97.9 | 96.1 |
| 56 | GL2689 | Landrace | GL2689 | IT224194 | North Korea | 23,582,383,022 | 22.65 | 98.3 | 96.9 |
| 57 | Gyeongsan199754 | Landrace | Kyoungbuk Kyoungsan-1997-54 | IT219319 | South Korea | 18,190,802,994 | 16.44 | 98 | 96 |
| 58 | Hadaedu | Landrace | Hadaedu | IT104627 | South Korea | 26,684,893,348 | 25.3 | 98.3 | 97 |
| 59 | Haenam-1998-16 | Landrace | Jeonnam heanam-1998-16 | IT224190 | South Korea | 21,473,266,966 | 19.61 | 98.4 | 97 |
| 60 | Haman | Landrace | Haman | A07 | South Korea | 20,860,329,718 | 20.89 | 98.2 | 96.4 |
| 61 | Hannam | Improved | Hannam | | South Korea | 15,531,802,318 | 14.96 | 97.2 | 91.7 |
| 62 | Heihokuta | Landrace | Heihokuta | PI 88820 | North Korea | 17,175,555,702 | 15.63 | 98.4 | 96.6 |
| 63 | Heugcheong | Landrace | Heugcheong | | South Korea | 15,763,684,864 | 15.02 | 96.8 | 90.3 |
| 64 | Heugseagyuwoldu | Landrace | Heksaekyuwoldu | IT102712 | South Korea | 16,314,806,778 | 15.85 | 98.2 | 96.1 |
| 65 | Hojang | Improved | Hojang | | South Korea | 14,833,073,676 | 14.14 | 97.4 | 90.8 |
| 66 | Horangi | Landrace | Horangi | IT021842 | South Korea | 19,135,348,462 | 18.92 | 98.4 | 96.9 |
| 67 | Hoseo | Improved | Hoseo | | South Korea | 15,121,025,542 | 14.52 | 96.6 | 89.6 |
| 68 | Hwangkeum | Improved | Hwangkeum | A09 | South Korea | 19,751,497,784 | 19.61 | 98.5 | 96.7 |
| 69 | Hwinkong | Landrace | Heuin | IT175923 | South Korea | 16,485,446,140 | 15.33 | 98.4 | 96.5 |
| 70 | Iksan10 | Improved | Iksan10 | | South Korea | 16,668,273,920 | 15.98 | 98.2 | 95.7 |
| 71 | Ilpumgeomjeong | Improved | Ilpumgeomjeong | A04 | South Korea | 16,999,533,412 | 15.94 | 98.3 | 95.2 |
| 72 | IT102595 | Landrace | IT102595 | | South Korea | 21,881,591,502 | 21.04 | 98.2 | 96.6 |
| 73 | IT102668 | Landrace | IT102668 | | South Korea | 21,791,912,904 | 21.02 | 98.2 | 96.6 |
| 74 | IT103189 | Landrace | IT103189 | | South Korea | 21,694,861,882 | 21.38 | 98.3 | 96.8 |
| 75 | IT104334 | Landrace | IT104334 | | South Korea | 15,369,753,950 | 14.79 | 96.8 | 91 |
| 76 | IT104887A | Landrace | IT104887 | | South Korea | 19,211,584,436 | 18.73 | 98.4 | 96.8 |
| 77 | IT112859 | Landrace | IT112859 | | South Korea | 15,419,810,450 | 14.85 | 96.5 | 89.7 |
| 78 | IT115870 | Landrace | IT115870 | | South Korea | 45,365,911,500 | 42.72 | 98.5 | 97.5 |
| 79 | IT121464 | Landrace | Kongnamul | IT121464 | South Korea | 15,494,911,206 | 14.46 | 96.8 | 90 |
| 80 | IT121504 | Landrace | IT121504 | | South Korea | 15,927,486,040 | 14.99 | 96.9 | 90.6 |

| | | | | | | | | | |
|---|---|---|---|---|---|---|---|---|---|
| 81 | IT155162 | Landrace | IT155162 | | South Korea | 16,953,622,848 | 16.24 | 98.2 | 96.1 |
| 82 | IT177322 | Landrace | IT177322 | | South Korea | 24,925,137,234 | 24.08 | 98.4 | 97 |
| 83 | IT177327 | Landrace | IT177327 | | South Korea | 24,472,242,934 | 23.5 | 98.4 | 97.1 |
| 84 | IT177337 | Landrace | IT177337 | | South Korea | 29,042,124,148 | 28.14 | 98.6 | 97.4 |
| 85 | IT177388 | Landrace | IT177388 | | South Korea | 15,427,628,626 | 14.64 | 96.7 | 90.4 |
| 86 | IT177413 | Landrace | Geomjeong-4 | IT177413 | South Korea | 21,272,201,708 | 19.11 | 98.3 | 96.7 |
| 87 | IT177513 | Landrace | IT177513 | | South Korea | 18,276,109,840 | 17.81 | 98.2 | 96.3 |
| 88 | IT177518 | Landrace | IT177518 | | South Korea | 22,147,904,464 | 20.76 | 98.3 | 96.9 |
| 89 | IT177528 | Landrace | IT177528 | | South Korea | 21,539,644,754 | 21.19 | 98.3 | 97 |
| 90 | IT177627 | Landrace | IT177627 | | South Korea | 19,630,029,596 | 18 | 98.2 | 96.5 |
| 91 | IT177793 | Landrace | IT177793 | | South Korea | 16,077,237,572 | 15.73 | 97.9 | 95.2 |
| 92 | IT177951 | Landrace | IT177951 | | South Korea | 23,237,305,138 | 22.6 | 98.1 | 96.6 |
| 93 | IT177955 | Landrace | IT177955 | | South Korea | 25,624,083,014 | 25.07 | 98.5 | 97.2 |
| 94 | IT178024 | Landrace | IT178024 | | South Korea | 18,575,583,610 | 17.04 | 98.3 | 96.5 |
| 95 | IT178037 | Landrace | IT178037 | | South Korea | 25,579,148,736 | 24.64 | 98.1 | 96.7 |
| 96 | IT180313 | Landrace | IT180313 | | South Korea | 26,280,630,410 | 25.41 | 98.4 | 97 |
| 97 | IT180422 | Landrace | IT180422 | Beijing da qing don, PI 495017 B | China | 26,826,965,926 | 25.56 | 98.3 | 97 |
| 98 | IT208844 | Landrace | IT208844 | | South Korea | 22,751,357,542 | 21.91 | 98.4 | 97.1 |
| 99 | IT220684 | Landrace | IT220684 | | South Korea | 33,983,571,364 | 32.3 | 98.5 | 97.4 |
| 100 | IT224436 | Landrace | Jeonnam Wando-2000-57 | IT224436 | South Korea | 21,437,663,280 | 21.03 | 98.3 | 96.7 |
| 101 | IT224873 | Landrace | IT224873 | | China | 16,799,522,516 | 16.5 | 97.9 | 95.6 |
| 102 | IT224875 | Landrace | Nan zhao cao huang dou | PI 567642 A | China | 17,321,490,558 | 16.74 | 98.4 | 96.5 |
| 103 | IT228363 | Landrace | IT228363 | PI 603910 A, Cin | South Korea | 21,097,209,620 | 20.83 | 98 | 96.2 |
| 104 | IT234975 | Landrace | IT234975 | | South Korea | 20,982,161,512 | 20.79 | 98.2 | 96.6 |
| 105 | IT238217 | Landrace | Gong xian huang dou | PI 567618 A | China | 19,258,038,982 | 18.4 | 98.3 | 96.6 |

| | | | | | | | | | |
|---|---|---|---|---|---|---|---|---|---|
| 106 | Jinheung-NO_11 | Landrace | Jeonnam jinheungwon No.11 | IT178701 | South Korea | 22,834,595,084 | 21.14 | 98.3 | 96.8 |
| 107 | Josaengseori | Improved | Josaengseori | | South Korea | 15,794,063,044 | 15.16 | 97 | 91.4 |
| 108 | Juinuni | Landrace | Juinuni | IT224191 | South Korea | 20,626,768,516 | 19.91 | 98.4 | 97 |
| 109 | KAERI_590-6 | Landrace | KAERI 590-6 | PI 408342 | South Korea | 19,223,353,980 | 18.51 | 98.2 | 96.4 |
| 110 | Kangwon2-33 | Landrace | Kangwon sujib2-33 | IT156161 | South Korea | 15,196,384,206 | 14.48 | 96.5 | 89.2 |
| 111 | Kangwon2-4 | Landrace | Kangwon sujib2-4 | IT156134 | South Korea | 24,568,956,622 | 23.62 | 98.5 | 97.2 |
| 112 | Kangwon3-24 | Landrace | Kangwon sujib3-24 | IT156183 | South Korea | 17,545,021,294 | 17.62 | 98.2 | 96.1 |
| 113 | Kangwon3-25 | Landrace | Kangwon sujib3-25 | IT156184 | South Korea | 31,645,164,224 | 30.12 | 98.6 | 97.3 |
| 114 | Kangwon3-31 | Landrace | Kangwon sujib3-31 | IT156190 | South Korea | 18,086,769,430 | 17.88 | 98.1 | 96 |
| 115 | Kangwon5-19 | Landrace | IT156217 | Kangwon sujib5-19 | South Korea | 19,244,471,934 | 18.55 | 98.4 | 96.6 |
| 116 | kangwon5-26 | Landrace | IT156223 | Kangwon sujib5-26 | South Korea | 22,993,480,606 | 22.66 | 98.3 | 96.9 |
| 117 | Kantou_44 | Improved | Kantou #44 | IT022368 | Japan | 46,545,390,016 | 44.11 | 98.6 | 97.6 |
| 118 | KAS_102-2 | Landrace | KAS 102-2 | IT229952 | South Korea | 16,300,498,320 | 15.96 | 98 | 95.3 |
| 119 | KAS_205-22 | Landrace | KAS 205-22 | PI 424258 | South Korea | 17,240,516,204 | 16.79 | 98 | 95.6 |
| 120 | KAS-100-12 | Landrace | KAS 100-12 | PI 398187 | South Korea | 18,505,153,284 | 16.76 | 98.1 | 96.4 |
| 121 | KAS-100-8-2 | Landrace | KAS 100-8-2 | PI 423727 | South Korea | 20,928,768,214 | 18.94 | 98.3 | 96.8 |
| 122 | KAS150-22 | Landrace | KAS150-22 | IT115352 | South Korea | 17,217,672,622 | 16.91 | 98.2 | 96.2 |
| 123 | KAS-160-5 | Landrace | KAS 160-5 | IT219609 | South Korea | 17,039,904,852 | 15.6 | 98.2 | 96.3 |
| 124 | KAS210-22 | Landrace | KAS210-22 | PI 458043 | South Korea | 15,818,129,424 | 15.11 | 97 | 90.9 |
| 125 | KAS241-4 | Landrace | KAS241-4 | PI 458093 | South Korea | 20,771,508,962 | 20.24 | 98.2 | 96.6 |
| 126 | KAS302-19 | Landrace | KAS302-19 | IT219770 | South Korea | 16,311,776,812 | 16.13 | 98.1 | 95.6 |
| 127 | KAS304-12 | Landrace | KAS304-12 | PI 424305 | South Korea | 21,684,979,536 | 19.91 | 98.4 | 97 |
| 128 | KAS331-12 | Landrace | KAS331-12 | IT115548 | South Korea | 15,207,559,112 | 14.6 | 96.4 | 89.5 |
| 129 | KAS331-13 | Landrace | KAS331-13 | IT115549 | South Korea | 25,645,727,052 | 24.84 | 98.4 | 97 |
| 130 | KAS361-2 | Landrace | KAS361-2 | IT219783 | South Korea | 19,248,631,078 | 18.85 | 98.2 | 96.3 |

| | | | | | | | | | |
|---|---|---|---|---|---|---|---|---|---|
| 131 | KAS380-14 | Landrace | KAS380-14 | PI 398582 | South Korea | 15,828,021,736 | 15.29 | 96.7 | 90.8 |
| 132 | KAS502-6 | Landrace | KAS502-6 | IT115594 | South Korea | 20,577,563,958 | 20.63 | 98.1 | 96.3 |
| 133 | KAS505-1 | Landrace | KAS505-1 | IT115608 | South Korea | 15,352,938,892 | 14.82 | 96.3 | 89.8 |
| 134 | KAS524-4 | Landrace | KAS524-4 | IT154281 | South Korea | 23,606,981,526 | 22.93 | 98.5 | 97.3 |
| 135 | KAS531-5 | Landrace | KAS531-5 | IT115665 | South Korea | 15,381,197,334 | 14.9 | 96.9 | 91.3 |
| 136 | KAS544-2 | Landrace | KAS544-2 | PI 407996 | South Korea | 20,076,058,530 | 19.71 | 98.2 | 96.5 |
| 137 | KAS571-23 | Landrace | KAS571-23 | IT115727 | South Korea | 15,300,373,376 | 14.78 | 97 | 91.5 |
| 138 | KAS574-11 | Landrace | KAS574-11 | PI 458231 | South Korea | 16,628,144,160 | 16.02 | 97 | 91.7 |
| 139 | KAS575-1 | Landrace | KAS575-1 | IT154323 | South Korea | 18,177,585,964 | 17.55 | 98.2 | 96.2 |
| 140 | KAS590-6 | Landrace | KAS590-6 | IT181672 | South Korea | 19,723,625,738 | 18.28 | 98.3 | 96.8 |
| 141 | KAS611-1 | Landrace | KAS611-1 | IT141910 | South Korea | 15,575,003,720 | 14.95 | 97 | 91.2 |
| 142 | KAS622-8 | Landrace | KAS622-8 | PI 424525 | South Korea | 15,927,289,740 | 15.23 | 97 | 90 |
| 143 | KAS625-19 | Landrace | KAS625-19 | IT115766 | South Korea | 15,812,024,192 | 15.11 | 97.2 | 91.7 |
| 144 | KAS629-10 | Landrace | KAS629-10 | IT115768 | South Korea | 25,037,267,116 | 24.08 | 98.3 | 97.2 |
| 145 | KAS636-21 | Landrace | KAS636-21 | IT154540 | South Korea | 22,020,561,936 | 21.33 | 98.2 | 96.6 |
| 146 | KAS638-10 | Landrace | KAS638-10 | IT154569 | South Korea | 16,135,893,824 | 15.49 | 96.8 | 91.1 |
| 147 | KAS640-12 | Landrace | KAS640-12 | PI 424567 | South Korea | 18,675,476,452 | 17.96 | 98 | 96 |
| 148 | KAS640-46 | Landrace | KAS640-46 | IT154621 | South Korea | 17,560,448,360 | 17.18 | 98.2 | 96.2 |
| 149 | KAS640-49 | Landrace | KAS640-49 | IT154624 | South Korea | 19,211,143,818 | 18.46 | 98.5 | 97 |
| 150 | KAS641-8 | Landrace | KAS641-8 | IT154634 | South Korea | 35,689,407,190 | 34.35 | 98.7 | 97.7 |
| 151 | KAS651-37 | Landrace | KAS651-37 | IT154735 | South Korea | 18,593,856,120 | 17.98 | 98.2 | 96.2 |
| 152 | KAS660-12 | Landrace | KAS660-12 | PI 408170 | South Korea | 18,645,236,286 | 17.18 | 98.3 | 96.7 |
| 153 | KAS660-21 | Landrace | KAS660-21 | IT115872 | South Korea | 15,392,098,628 | 14.5 | 96.8 | 89.6 |
| 154 | KAS663-8 | Landrace | KAS663-8 | IT115880 | South Korea | 20,544,645,656 | 19.86 | 98.5 | 97.1 |
| 155 | Keonol | Landrace | Keunol | | South Korea | 15,821,031,040 | 15.16 | 96.8 | 90.7 |
| 156 | Kershaw | Improved | Kershaw | PI 548985 | United States | 28,005,027,156 | 27.16 | 98.4 | 97.2 |
| 157 | Keumkangdaelib | Improved | Keumkangdaelib | IT021889 | South Korea | 25,308,091,052 | 24.82 | 98.3 | 96.9 |
| 158 | KLG_16001 | Landrace | KLG 16001 | IT238318, PI 96978 | North Korea | 23,370,699,142 | 22.46 | 98 | 96.4 |

| | | | | | | | | | |
|---|---|---|---|---|---|---|---|---|---|
| 159 | KLS_808-1 | Landrace | KLS 808-1 | PI 399025 | South Korea | 30,810,652,456 | 29.35 | 98.4 | 97.1 |
| 160 | KLS087019 | Landrace | KLS087019 | IT153356 | South Korea | 21,060,772,414 | 20.49 | 98.2 | 96.4 |
| 161 | KLS087062 | Landrace | KLS087062 | IT153382 | South Korea | 15,722,106,108 | 15.09 | 97.2 | 91.9 |
| 162 | KLS087160 | Landrace | KLS087160 | IT153411 | South Korea | 16,113,499,316 | 14.75 | 97.9 | 94.6 |
| 163 | KLS105 | Landrace | KLS105 | IT022396 | South Korea | 18,670,237,054 | 17.86 | 97 | 92.5 |
| 164 | KLS-116-1 | Landrace | KLS 116-1 | PI 398874 | South Korea | 15,530,791,222 | 14.83 | 96.3 | 89.1 |
| 165 | KLS117 | Landrace | KLS117 | PI 398875 | South Korea | 16,283,184,056 | 15.51 | 96.1 | 88.7 |
| 166 | KLS123-1 | Landrace | KLS123-1 | PI 398879 | South Korea | 16,956,681,504 | 16.08 | 96.8 | 90.3 |
| 167 | KLS137-1 | Landrace | KLS137-1 | PI 398888 | South Korea | 26,476,233,394 | 25.76 | 98.5 | 97.3 |
| 168 | KLS419 | Landrace | KLS419 | IT022505 | South Korea | 21,126,697,504 | 20.63 | 98.5 | 97.1 |
| 169 | KLS606-2 | Landrace | KLS606-2 | IT022527 | South Korea | 19,256,066,922 | 19 | 98.3 | 96.6 |
| 170 | KLS714-2 | Landrace | KLS714-2 | IT022595 | South Korea | 30,812,031,388 | 29.51 | 98.3 | 97 |
| 171 | KLS72O-1 | Landrace | KLS72O-1 | IT022602 | South Korea | 23,485,785,000 | 22.7 | 98.3 | 97 |
| 172 | KLS739-2 | Landrace | K.L.S 739-2 | IT021695 | South Korea | 20,734,306,790 | 19.29 | 97.6 | 93.3 |
| 173 | KLS77013 | Landrace | KLS77013 | IT025215 | South Korea | 20,699,258,180 | 20.05 | 98.3 | 96.8 |
| 174 | KLS77048 | Landrace | KLS77048 | IT025253 | South Korea | 15,741,559,740 | 15.03 | 97 | 91.3 |
| 175 | KLS77114-1 | Landrace | KLS77114-1 | IT025335 | South Korea | 16,792,781,574 | 16.54 | 98.1 | 95.8 |
| 176 | KLS77131-2 | Landrace | KLS77131-2 | IT025370 | South Korea | 16,619,174,156 | 15.93 | 96.6 | 90.2 |
| 177 | KLS77170 | Landrace | KLS77170 | IT025435 | South Korea | 16,289,632,058 | 15.52 | 96.8 | 91.1 |
| 178 | KLS77196-1 | Landrace | KLS77196-1 | IT025484 | South Korea | 17,969,069,158 | 17.72 | 98.3 | 96.5 |
| 179 | KLS85044 | Landrace | KLS85044 | IT142986 | South Korea | 16,634,736,216 | 16.07 | 98.3 | 96.2 |
| 180 | KLS85109 | Landrace | KLS85109 | IT143050 | South Korea | 17,823,684,244 | 17.28 | 98.3 | 96.5 |
| 181 | KLS85250 | Landrace | KLS85250 | IT143167 | South Korea | 16,517,878,826 | 16.18 | 98.3 | 96.1 |
| 182 | KLS86037 | Landrace | KLS86037 | IT143237 | South Korea | 23,732,142,406 | 23.07 | 98.4 | 97 |
| 183 | KLS-86083-R_L1 | Landrace | KLS86083 | IT142839 | South Korea | 20,308,062,300 | 14.25 | 97.5 | 94.5 |
| 184 | KLS86094 | Landrace | KLS86094 | IT142878 | South Korea | 22,207,174,078 | 21.45 | 98.1 | 96.5 |
| 185 | KLS86097 | Landrace | KLS86097 | IT142891 | South Korea | 48,401,642,378 | 44.15 | 98.6 | 97.8 |
| 186 | KLS87005 | Landrace | KLS87005 | IT155984 | South Korea | 18,768,627,144 | 17.24 | 98.4 | 96.9 |
| 187 | KLS87096 | Landrace | KLS87096 | IT153718 | South Korea | 16,176,206,294 | 15.85 | 97.9 | 95.4 |

| | | | | | | | | | |
|---|---|---|---|---|---|---|---|---|---|
| 188 | KLS87113 | Landrace | KLS87113 | IT153730 | South Korea | 18,020,279,600 | 17.14 | 96.4 | 88.8 |
| 189 | KLS87277 | Landrace | KLS87277 | IT155954 | South Korea | 21,627,874,356 | 21.28 | 98.3 | 97 |
| 190 | KLS87323 | Landrace | KLS87323 | IT156001 | South Korea | 16,830,666,568 | 16.51 | 98.1 | 95.9 |
| 191 | KLS87345 | Landrace | KLS87345 | IT156261 | South Korea | 28,940,659,094 | 28.11 | 98.5 | 97.5 |
| 192 | KLS87348 | Landrace | KLS87348 | IT156003 | South Korea | 25,769,338,974 | 24.09 | 98.4 | 97.2 |
| 193 | KLS87352 | Landrace | KLS87352 | IT156006 | South Korea | 15,994,140,158 | 14.6 | 97.5 | 93.8 |
| 194 | KLS88035 | Landrace | KLS88035 | IT163579 | South Korea | 17,515,649,076 | 15.98 | 98.4 | 96.6 |
| 195 | KLS88046 | Landrace | KLS88046 | IT160109 | South Korea | 18,402,030,250 | 17.85 | 98.3 | 96.6 |
| 196 | KLS88048 | Landrace | KLS88048 | IT160111 | South Korea | 16,603,112,588 | 15.54 | 98.3 | 96.5 |
| 197 | KLS88066 | Landrace | KLS88066 | IT160129 | South Korea | 21,271,204,806 | 19.53 | 98.3 | 96.9 |
| 198 | KLS88068-2 | Landrace | KLS88068-2 | IT160132 | South Korea | 18,872,176,602 | 18.78 | 98.3 | 96.6 |
| 199 | Kongnamul | Landrace | Kongnamul | IT104023 | South Korea | 26,177,983,630 | 25.35 | 98.2 | 96.9 |
| 200 | Kwangan | Improved | Gwangan | | South Korea | 17,441,167,722 | 16.44 | 96.9 | 90.4 |
| 201 | kwangkyo | Improved | Kwangkyo | IT023515 | South Korea | 26,094,204,300 | 25.34 | 98 | 96.5 |
| 202 | KwangkyoV2 | Improved | KwangkyoV2 | A03 | South Korea | 21,598,195,318 | 19.8 | 98 | 95.7 |
| 203 | L62-667 | Improved | L62-667 | PI 547716 | United States | 15,656,967,124 | 14.97 | 96.7 | 89.8 |
| 204 | L-B | Landrace | L-B | IT228619 | South Korea | 17,969,665,306 | 17.37 | 98.3 | 96.2 |
| 205 | Lindarin_63 | Improved | Lindarin #63 | PI 548590 | United States | 17,054,765,970 | 16.46 | 98.4 | 96.4 |
| 206 | Lindou_9 | Improved | Lindou 9 | | China | 29,964,045,890 | 28.69 | 98.1 | 96.8 |
| 207 | Manpung | Improved | Manpung | | South Korea | 15,288,363,742 | 14.8 | 98.3 | 95.7 |
| 208 | Marshall | Improved | Marshall | PI 548693 | United States | 26,371,174,238 | 25.17 | 98.6 | 97.5 |
| 209 | Meju | Landrace | Meju | IT108810 | South Korea | 15,120,481,942 | 14.62 | 96.7 | 90.4 |
| 210 | Meju_Jeonbuk | Landrace | Meju_Jeonnam | IT103340 | South Korea | 17,438,438,850 | 16.73 | 98.1 | 96.2 |
| 211 | Meju_Jeonnam | Landrace | Meju_Jeonbuk | IT162760 | South Korea | 17,882,071,716 | 17.23 | 98 | 95.8 |
| 212 | Milyang_26 | Improved | Milyang26 | IT157944 | South Korea | 15,133,197,350 | 14.7 | 98.5 | 96.6 |
| 213 | Milyang12 | Improved | Milyang12 | IT24623 | South Korea | 16,136,661,508 | 15.4 | 97.1 | 90.6 |
| 214 | Milyang206 | Improved | Milyang206 | | South Korea | 31,008,134,182 | 28.36 | 98.8 | 97.9 |
| 215 | Mote | Landrace | Mote | PI 88814 | North Korea | 27,377,243,448 | 25.86 | 98.5 | 97.3 |

| | | | | | | | | | |
|---|---|---|---|---|---|---|---|---|---|
| 216 | Myeongjunamul | Improved | Myeongjunamul | | South Korea | 17,259,848,130 | 15.65 | 98.2 | 95.8 |
| 217 | Nampung | Improved | Nampung | | South Korea | 16,710,511,338 | 15.89 | 98.6 | 96.8 |
| 218 | Namul | Landrace | Namul | IT178511 | South Korea | 16,620,684,458 | 15.27 | 98.3 | 96.4 |
| 219 | Nezumi_Meta | Landrace | Nezumi Meta | PI 417193 | South Korea | 16,145,442,460 | 15.9 | 98 | 95.8 |
| 220 | No_39_Green | Landrace | No. 39 Green | PI 171430 | China | 45,535,515,304 | 41.09 | 98.5 | 97.6 |
| 221 | Nokwon | Improved | Nogwon | | South Korea | 16,091,118,096 | 15.34 | 96.6 | 89.8 |
| 222 | Nonduleong | Landrace | Nonduleong | IT109029 | South Korea | 16,880,672,030 | 16.59 | 98.2 | 96.2 |
| 223 | Nonglim73 | Improved | Nonglim73 | IT135700 | Japan | 22,789,591,648 | 21.71 | 98.3 | 96.9 |
| 224 | Ogden | Improved | Ogden | PI 548477 | United States | 37,871,339,372 | 34.88 | 98.6 | 97.7 |
| 225 | Oial | Landrace | Oial | IT162657 | South Korea | 41,148,156,850 | 37.91 | 98.5 | 97.6 |
| 226 | Ol | Landrace | Ol | IT103926 | South Korea | 27,812,247,570 | 26.93 | 98.3 | 97.1 |
| 227 | ORD_8139 | Landrace | ORD 8139 | PI 407803 | South Korea | 33,308,526,166 | 32.43 | 98.3 | 97 |
| 228 | OT89-06 | Improved | OT89-06 | PI 546044 | Canada | 23,055,101,894 | 22.61 | 98.5 | 97.2 |
| 229 | OT94-51 | Improved | OT94-51 | PI 591432 | Canada | 17,107,681,504 | 16.39 | 97.2 | 90.7 |
| 230 | Paldonamul | Improved | Paldonamul | | South Korea | 16,080,136,772 | 15.38 | 96.5 | 90.3 |
| 231 | Peking | Landrace | Peking | | China | 23,742,417,430 | 18.24 | 98.3 | 95.9 |
| 232 | PI_157430 | Landrace | I chu tau chow | PI 157430 | South Korea | 39,452,739,118 | 37.23 | 98.5 | 97.6 |
| 233 | PI_159764 | Landrace | PI159764 | | South Korea | 19,908,287,564 | 19.58 | 98.2 | 96.6 |
| 234 | PI_196175 | Landrace | PI 196175 | | South Korea | 19,398,055,544 | 18.94 | 98.2 | 96.4 |
| 235 | PI_339982 | Landrace | PI 339982 | | South Korea | 20,690,145,934 | 20.19 | 98.4 | 96.9 |
| 236 | PI_340003 | Landrace | PI 340003 | | South Korea | 17,649,503,932 | 17.19 | 98.2 | 96.4 |
| 237 | PI_391597 | Improved | Niu mao huang | IT238204, PI 391597 | China | 29,051,753,418 | 26.45 | 98.3 | 97 |
| 238 | PI_399062 | Landrace | PI 399062 | | South Korea | 17,578,386,858 | 16.87 | 98.3 | 96.3 |
| 239 | PI_399079 | Landrace | PI 399079 | | South Korea | 22,869,506,284 | 22.12 | 98.3 | 96.7 |
| 240 | PI_399089 | Landrace | PI 399089 | | South Korea | 17,775,163,112 | 17.32 | 98.2 | 96.3 |
| 241 | PI_399100 | Landrace | PI 399100 | | South Korea | 21,691,624,744 | 21.11 | 98.1 | 96.5 |
| 242 | PI_399108 | Landrace | PI 399108 | | South Korea | 17,360,119,378 | 17.28 | 98.4 | 96.6 |
| 243 | PI_399113 | Landrace | PI 399113 | | South Korea | 21,044,085,102 | 20.59 | 98.3 | 96.8 |

| | | | | | | | | | |
|---|---|---|---|---|---|---|---|---|---|
| 244 | PI_399126 | Landrace | PI 399126 | | South Korea | 17,961,128,672 | 16.8 | 98.1 | 95.9 |
| 245 | PI_407736 | Landrace | PI407736 | | China | 24,783,481,718 | 23.66 | 98.4 | 97.2 |
| 246 | PI_407810 | Landrace | PI 407810 | | South Korea | 30,579,092,144 | 28.88 | 98.4 | 97.2 |
| 247 | PI_458156 | Landrace | PI 458156 | | South Korea | 17,500,472,670 | 16.74 | 98.2 | 95.9 |
| 248 | PI_458184 | Landrace | PI 458184 | | South Korea | 22,330,143,646 | 22.03 | 98 | 96.3 |
| 249 | PI_458232 | Landrace | PI 458232 | | South Korea | 45,983,421,470 | 44.07 | 98.7 | 97.9 |
| 250 | PI_458260 | Landrace | PI 458260 | | South Korea | 19,921,563,786 | 19.14 | 98.2 | 96.6 |
| 251 | PI_458277 | Landrace | PI 458277 | | South Korea | 18,383,977,898 | 17.88 | 98 | 96.1 |
| 252 | PI_475824_A | Landrace | PI475824A | | China | 19,104,781,532 | 18.55 | 98 | 95.9 |
| 253 | PI_556949 | Landrace | PI 556949 | | China | 17,856,518,892 | 17.42 | 97.9 | 95.7 |
| 254 | PI_567317 | Landrace | Hua huang dou | IT238194 | China | 21,876,469,884 | 21.31 | 98.2 | 96.7 |
| 255 | PI_567366_A | Landrace | Wu tong shu huang dou | IT238326 | China | 20,397,930,734 | 19.86 | 98.1 | 96.4 |
| 256 | PI_567458 | Landrace | Ji li huang dou | PI 567458 | China | 42,774,947,196 | 38.64 | 98.3 | 97.1 |
| 257 | PI_567519 | Landrace | Bai hua chi | PI 567519 | China | 17,889,406,994 | 17.43 | 98.1 | 96.2 |
| 258 | PI_567591 | Landrace | PI 567591 | | China | 17,993,667,058 | 17.41 | 98.2 | 96.3 |
| 259 | PI_567601 | Landrace | Xiao tie jiao | PI 567601 | China | 20,068,268,742 | 19.24 | 98.3 | 96.7 |
| 260 | PI_567660_A | Landrace | PI 567660 A | | China | 36,914,966,074 | 34.62 | 98.7 | 97.7 |
| 261 | PI_567759 | Landrace | Pei xian xiao bai jian ke | PI 567759 | China | 17,663,692,194 | 17.35 | 97.9 | 95.4 |
| 262 | PI_567773 | Landrace | Tong shan huang da dou jia | PI 567773 | China | 24,681,309,682 | 23.63 | 98.3 | 97 |
| 263 | PI_587718 | Landrace | Huang pi feng zi wo | PI 587718 | China | 22,765,391,180 | 22.34 | 98.3 | 96.9 |
| 264 | PI_603174_A | Landrace | PI 603174 A | | North Korea | 17,151,916,350 | 16.36 | 98.1 | 95.9 |
| 265 | PI_603176_A | Landrace | PI 603176 A | | North Korea | 33,013,850,874 | 31.72 | 98.5 | 97.4 |
| 266 | PI_62203-8 | Landrace | PI 62203-8 | | China | 20,050,613,520 | 19.53 | 98 | 95.9 |
| 267 | PI_64698 | Landrace | PI 64698 | | South Korea | 19,695,196,062 | 19.06 | 98.3 | 96.5 |
| 268 | PI_72227 | Landrace | PI 72227 | | China | 36,924,408,104 | 35.42 | 98.5 | 97.4 |
| 269 | PI_81037 | Landrace | PI 81037 | | Japan | 25,932,264,048 | 25.15 | 98.3 | 96.9 |
| 270 | PI_82218 | Landrace | PI 82218 | | South Korea | 19,675,456,134 | 19.12 | 98.3 | 96.7 |

| | | | | | | | | |
|---|---|---|---|---|---|---|---|---|
| 271 | PI_82246 | Landrace | PI 82246 | South Korea | 17,815,353,574 | 17.27 | 98 | 95.9 |
| 272 | PI_82555 | Landrace | PI 82555 | South Korea | 18,321,967,332 | 17.78 | 98.1 | 96.1 |
| 273 | PI_83853 | Landrace | PI 83853 | South Korea | 23,196,655,334 | 22.74 | 98.4 | 97.1 |
| 274 | PI_84646 | Landrace | PI 84646 | South Korea | 19,763,142,136 | 19.15 | 98.2 | 96.5 |
| 275 | PI_84646-2 | Landrace | PI 84646-2 | South Korea | 24,574,816,932 | 23.91 | 98.5 | 97.3 |
| 276 | PI_84928 | Landrace | PI 84928 | North Korea | 26,052,165,900 | 25.03 | 98.2 | 96.9 |
| 277 | PI_87002 | Landrace | PI 87002 | South Korea | 21,527,738,706 | 20.85 | 98.2 | 96.7 |
| 278 | PI_87565 | Landrace | PI 87565 | North Korea | 30,317,310,692 | 27.58 | 98.3 | 96.9 |
| 279 | PI_87630 | Landrace | PI87630 | Japan | 35,014,983,508 | 32.64 | 98.6 | 97.6 |
| 280 | PI_87631-1 | Landrace | PI87631-1 | Japan | 37,665,855,250 | 35.12 | 98.2 | 97 |
| 281 | PI_87632 | Landrace | PI 87632 | Japan | 18,152,669,756 | 17.68 | 98.4 | 96.6 |
| 282 | PI_88816-S | Landrace | PI 88816-S | North Korea | 24,441,261,962 | 23 | 98.2 | 96.7 |
| 283 | PI_89138 | Landrace | PI 89138 | North Korea | 22,580,237,396 | 21.92 | 98.3 | 96.9 |
| 284 | PI_89143 | Landrace | PI 89143 | North Korea | 27,429,407,606 | 26.48 | 98.3 | 96.9 |
| 285 | PI_89154-S | Landrace | PI 89154-S | North Korea | 16,491,097,768 | 16.21 | 98.3 | 96.1 |
| 286 | PI_90763 | Landrace | PI 90763 | China | 27,381,626,676 | 26.7 | 98.2 | 96.8 |
| 287 | PI_91073 | Landrace | PI 91073 | South Korea | 19,840,677,918 | 19.37 | 98.2 | 96.5 |
| 288 | PI_91725 | Landrace | PI 91725 | North Korea | 30,647,125,798 | 29.46 | 98.4 | 97.1 |
| 289 | PI_96280 | Landrace | PI 96280 | North Korea | 18,764,987,742 | 18.3 | 98.3 | 96.5 |
| 290 | PI_96354 | Landrace | PI 96354 | North Korea | 19,626,376,604 | 19.11 | 98.2 | 96.4 |
| 291 | PI_96549 | Landrace | PI 96549 | North Korea | 31,447,217,720 | 29.15 | 98.5 | 97.4 |
| 292 | PI_96786 | Landrace | PI 96786 | North Korea | 64,967,740,034 | 61.27 | 98.8 | 98 |
| 293 | PI196168 | Landrace | PI 196168 | South Korea | 19,971,451,468 | 18.05 | 98.4 | 96.9 |
| 294 | PI227159 | Landrace | PI 227159 | South Korea | 25,949,446,036 | 23.82 | 98.5 | 97.4 |
| 295 | PI339736 | Landrace | PI339736 | South Korea | 18,914,502,808 | 17.48 | 98.1 | 96.2 |
| 296 | PI-399122 | Landrace | PI 399122 | South Korea | 16,667,131,454 | 15.71 | 98.1 | 96 |
| 297 | PI407795A | Landrace | PI 407795A | South Korea | 16,080,256,364 | 14.83 | 98.2 | 96.1 |
| 298 | PI417329 | Landrace | PI 417329 | Japan | 26,872,754,864 | 26.57 | 98.3 | 97.1 |
| 299 | PI-458175-C | Landrace | PI 458175 C | South Korea | 23,408,534,910 | 21.32 | 98.3 | 96.9 |
| 300 | PI-458269 | Landrace | PI 458269 | South Korea | 20,792,369,914 | 19.21 | 98.3 | 96.9 |

| | | | | | | | | |
|---|---|---|---|---|---|---|---|---|
| 301 | PI475812B | Landrace | PI 475812 B | China | 17,649,905,290 | 16.1 | 97.6 | 94.7 |
| 302 | PI475814 | Landrace | PI 475814 | China | 15,659,173,838 | 14.95 | 97.9 | 95.1 |
| 303 | PI54818 | Landrace | PI 54818 | China | 15,614,123,290 | 15.29 | 97.9 | 95.4 |
| 304 | PI60269-2 | Landrace | PI 60269-2 | South Korea | 16,541,815,346 | 15.34 | 97.9 | 95.5 |
| 305 | PI68484-4 | Landrace | PI 68484-4 | China | 16,676,716,632 | 16.09 | 97.9 | 95.8 |
| 306 | PI68696 | Landrace | PI 68696 | China | 21,404,616,628 | 20.57 | 98.4 | 96.8 |
| 307 | PI80470 | Landrace | PI 80470 | Japan | 23,850,938,032 | 22.99 | 98.5 | 97.2 |
| 308 | PI82183 | Landrace | PI 82183 | South Korea | 23,948,686,674 | 22.71 | 98.4 | 97 |
| 309 | PI82235 | Landrace | PI 82235 | South Korea | 22,113,270,198 | 20.44 | 98.3 | 96.6 |
| 310 | PI82278 | Landrace | PI 82278 | South Korea | 17,799,831,680 | 16.4 | 98.2 | 96.2 |
| 311 | PI82291 | Landrace | PI 82291 | South Korea | 19,889,359,714 | 17.92 | 98.3 | 96.8 |
| 312 | PI82295 | Landrace | PI 82295 | South Korea | 15,550,182,038 | 14.7 | 97.9 | 95.1 |
| 313 | PI82544 | Landrace | PI 82544 | North Korea | 17,811,309,492 | 16.56 | 98.4 | 96.8 |
| 314 | PI83868 | Landrace | PI 83868 | North Korea | 24,007,454,968 | 23.3 | 98.4 | 97.2 |
| 315 | PI83893 | Landrace | PI 83893 | South Korea | 24,460,411,178 | 23.54 | 98.4 | 97.2 |
| 316 | PI84581 | Landrace | PI 84581 | South Korea | 19,925,485,256 | 18.82 | 98.4 | 97 |
| 317 | PI84609 | Landrace | PI 84609 | South Korea | 20,568,972,662 | 19.73 | 97.9 | 96 |
| 318 | PI84611 | Landrace | PI 84611 | South Korea | 20,512,161,026 | 18.89 | 98.1 | 96.3 |
| 319 | PI84642 | Landrace | PI 84642 | South Korea | 19,012,237,256 | 17.47 | 98.1 | 96.2 |
| 320 | PI84644 | Landrace | PI 84644 | South Korea | 17,707,105,902 | 16.3 | 98.3 | 96.5 |
| 321 | PI84669 | Landrace | PI 84669 | South Korea | 17,533,848,502 | 15.51 | 97.1 | 90 |
| 322 | PI84669N | Landrace | PI 84669N | South Korea | 17,938,878,822 | 16.25 | 97.6 | 94.4 |
| 323 | PI84680 | Landrace | PI 84680 | South Korea | 16,237,821,844 | 15.08 | 97.6 | 94.2 |
| 324 | PI84734 | Landrace | PI 84734 | South Korea | 18,318,725,966 | 16.89 | 98.3 | 96.6 |
| 325 | PI84946-2 | Landrace | PI 84946-2 | South Korea | 16,518,023,484 | 15.92 | 98 | 95.5 |
| 326 | PI85089 | Landrace | PI 85089 | South Korea | 21,015,884,342 | 20.05 | 98.4 | 96.9 |
| 327 | PI86490 | Landrace | PI 86490 | Japan | 21,464,358,570 | 19.82 | 98.2 | 96.5 |
| 328 | PI86904-1 | Landrace | PI 86904-1 | South Korea | 18,565,376,614 | 17.06 | 98.4 | 96.7 |
| 329 | PI87574 | Landrace | PI 87574 | North Korea | 22,087,935,116 | 20.35 | 98.4 | 97 |
| 330 | PI87619-1 | Landrace | PI 87619-1 | North Korea | 15,705,718,380 | 15.1 | 97.9 | 95.1 |

| | | | | | | | | | |
|---|---|---|---|---|---|---|---|---|---|
| 331 | PI88306-1 | Landrace | PI 88306-1 | | China | 22,499,577,424 | 21.11 | 98.1 | 96.5 |
| 332 | PI88820 | Landrace | PI 88820 | | North Korea | 20,714,228,018 | 19.06 | 98.2 | 96.7 |
| 333 | PI89128 | Landrace | PI 89128 | | North Korea | 15,828,871,564 | 15.23 | 97.9 | 95.4 |
| 334 | PI91083 | Landrace | PI 91083 | | South Korea | 17,812,554,336 | 16.08 | 97.8 | 95 |
| 335 | PI92568 | Landrace | PI 92568 | | China | 15,474,602,914 | 14.99 | 97.9 | 95.1 |
| 336 | PI93559 | Landrace | PI 93559 | | China | 16,467,642,334 | 16.21 | 98 | 95.6 |
| 337 | PI95853 | Landrace | PI 95853 | | South Korea | 17,476,601,684 | 16.04 | 98.4 | 96.6 |
| 338 | PI96089-5 | Landrace | PI 96089-5 | | North Korea | 17,473,348,842 | 16.05 | 98.3 | 96.4 |
| 339 | PI96983 | Landrace | PI96983 | A06 | North Korea | 18,022,767,644 | 17.48 | 98.3 | 95.8 |
| 340 | Pu_Ru_Ragi | Landrace | IT196925 | | North Korea | 26,120,433,302 | 25.69 | 98.4 | 97.2 |
| 341 | Pungwon | Improved | Pungwon | | South Korea | 18,507,147,994 | 17.46 | 97.4 | 90.5 |
| 342 | Pureun | Improved | Pureun | A02 | South Korea | 27,962,663,050 | 26.67 | 99.2 | 98.5 |
| 343 | Pureun-3 | Improved | Pureun-3 | | South Korea | 15,820,917,488 | 15.47 | 98.8 | 96.9 |
| 344 | Pureundogseagi | Landrace | Pureundogseagi | IT229464 | South Korea | 24,866,265,958 | 24 | 98.4 | 96.9 |
| 345 | Qi_si_wa | Landrace | Qi si wa | IT238327 | China | 24,613,410,418 | 23.88 | 98.2 | 96.7 |
| 346 | RAIKO | Improved | RAIKO | IT120618 | Japan | 18,454,972,360 | 17.87 | 98.2 | 96.3 |
| 347 | Ryong_song | Landrace | Ryong song | PI 612612 A | South Korea | 24,695,314,630 | 24.04 | 98.3 | 96.9 |
| 348 | Saebyeol | Improved | Saebyeol | | South Korea | 16,647,354,078 | 15.93 | 96.9 | 91.7 |
| 349 | Saedanbaek | Improved | Saedanbaek | | South Korea | 17,254,799,596 | 15.69 | 97.9 | 94.5 |
| 350 | Saline | Improved | Saline | PI 578057 | United States | 17,257,157,008 | 16.98 | 98.7 | 97 |
| 351 | Sangjupureun | Landrace | Sangjupureun | IT220685 | South Korea | 41,682,127,580 | 38.94 | 98.6 | 97.6 |
| 352 | Savoy | Improved | Savoy | PI 597381 | United States | 17,094,827,780 | 16.54 | 98.2 | 96.2 |
| 353 | seonam | Improved | Sunam | | South Korea | 17,220,396,662 | 16.32 | 96.7 | 91.2 |
| 354 | Seonbi | Landrace | Seonbi | IT224183 | South Korea | 17,759,640,916 | 16.28 | 98.3 | 96.5 |
| 355 | Seoritae | Landrace | Seoritae | A05 | South Korea | 18,740,206,398 | 17.75 | 98.4 | 95.9 |
| 356 | Shin2 | Landrace | PI 507238 | Shin 2 | Japan | 31,506,184,730 | 30.19 | 98.5 | 97.4 |
| 357 | Shinhwa | Improved | Shinhwa | | South Korea | 15,538,838,616 | 14.59 | 96.8 | 90 |
| 358 | Si_li_da_dou | Landrace | Si li da dou | PI 567587 A | China | 30,977,356,456 | 28.8 | 98.4 | 97.2 |
| 359 | Sinpaldal | Improved | Sinpaldal | | South Korea | 21,942,741,670 | 20.83 | 97.9 | 93.1 |

| | | | | | | | | | |
|---|---|---|---|---|---|---|---|---|---|
| 360 | Sinpaldal2 | Improved | Sinpaldal2 | | South Korea | 15,286,611,236 | 14.65 | 97 | 90.4 |
| 361 | Sinrok | Improved | Shillog | | South Korea | 16,203,112,380 | 15.42 | 96.8 | 90.8 |
| 362 | SLS_B229-1 | Landrace | SLS B229-1 | IT158136 | South Korea | 16,866,462,930 | 15.37 | 97.7 | 94.5 |
| 363 | SLS_B30-2 | Landrace | SLS B30-2 | IT158125 | South Korea | 32,951,063,866 | 31.34 | 98.6 | 97.6 |
| 364 | SLS90-101 | Landrace | SLS90-101 | IT167739 | South Korea | 45,454,756,880 | 42.36 | 98.6 | 97.7 |
| 365 | SLS90-102 | Landrace | SLS90-102 | IT167740 | South Korea | 20,080,438,738 | 19.34 | 98.2 | 96.5 |
| 366 | SLS90-146 | Landrace | SLS90-146 | IT167784 | South Korea | 23,278,893,860 | 22.59 | 98.4 | 96.9 |
| 367 | SLSB322-3 | Landrace | SLSB322-3 | IT134413 | South Korea | 15,645,417,134 | 14.67 | 97.2 | 91.1 |
| 368 | SLSB397-1 | Landrace | SLSB397-1 | IT134454 | South Korea | 17,130,408,212 | 16.07 | 97.1 | 91.6 |
| 369 | SLSB406-2 | Landrace | SLSB406-2 | IT134433 | South Korea | 20,379,294,314 | 19.94 | 98.3 | 96.6 |
| 370 | SLSB-B-15 | Landrace | SLSB-B-15 | IT134470 | South Korea | 15,435,095,878 | 14.48 | 96.9 | 90.3 |
| 371 | SLSB-B-27 | Landrace | SLSB-B-27 | IT134477 | South Korea | 15,432,059,570 | 14.87 | 96.9 | 90.3 |
| 372 | SLSJ101-1 | Landrace | SLSJ101-1 | IT134220 | South Korea | 15,996,257,782 | 14.99 | 96.8 | 90 |
| 373 | SLSJ190-2 | Landrace | SLSJ190-2 | IT134248 | South Korea | 17,164,558,070 | 15.93 | 97.5 | 94.3 |
| 374 | Socheong | Improved | Socheong | | South Korea | 19,649,851,788 | 20.58 | 98.7 | 96.3 |
| 375 | Socheong2 | Improved | Socheong2 | | South Korea | 16,927,986,068 | 15.73 | 97.3 | 91.6 |
| 376 | Sogcheong | Landrace | Sogcheong | IT228629 | South Korea | 26,974,399,306 | 25.92 | 98.4 | 97.3 |
| 377 | Soheung2 | Improved | Soheung2 | | Japan | 18,243,212,678 | 17.81 | 98.1 | 96.3 |
| 378 | Soho | Improved | Bangsa | | South Korea | 16,389,802,740 | 15.69 | 97.1 | 91.4 |
| 379 | Sohwang | Improved | Sohwang | | South Korea | 16,650,221,266 | 15.8 | 96.6 | 89.4 |
| 380 | Somyeongnamul | Improved | Somyeong | | South Korea | 17,088,458,600 | 15.79 | 96.9 | 89.9 |
| 381 | Sowon | Improved | Sowon | A01 | South Korea | 28,645,208,324 | 26.18 | 98.5 | 97.2 |
| 382 | Sowon2010 | Improved | Sowon2010 | | South Korea | 25,957,564,400 | 25.31 | 98.4 | 97.1 |
| 383 | Sugae_30 | Improved | Sugye30 | IT023248 | South Korea | 20,150,884,768 | 19.66 | 98.1 | 96.4 |
| 384 | Sugae_42 | Improved | Sugae #42 | IT230802 | South Korea | 16,948,316,708 | 16.05 | 97.9 | 95.7 |
| 385 | Sugae_43 | Improved | Sugae #43(A) | IT229363 | South Korea | 28,943,666,108 | 28.27 | 98.5 | 97.4 |
| 386 | Suigen_Ao | Landrace | Suigen Ao | IT230695 | South Korea | 27,425,836,758 | 25.84 | 98.2 | 96.8 |
| 387 | Suwon_2 | Improved | Suwongaetong #2 | IT229364 | South Korea | 24,964,661,484 | 24.2 | 98.3 | 96.9 |
| 388 | Suwon103 | Improved | Suwon103 | IT024470 | South Korea | 34,559,267,320 | 32.53 | 98.6 | 97.8 |

| | | | | | | | | | |
|---|---|---|---|---|---|---|---|---|---|
| 389 | Suwon116 | Improved | Suwon116 | IT021748 | South Korea | 16,384,165,306 | 15.61 | 96.9 | 90.5 |
| 390 | Suwon98 | Improved | Suwon98 | IT024222 | South Korea | 15,281,708,266 | 14.57 | 96.8 | 89.9 |
| 391 | Taekwang | Improved | Taekwang | | South Korea | 16,055,646,988 | 15.4 | 97 | 91.1 |
| 392 | Uram | Improved | Uram | | South Korea | 15,081,272,678 | 14.36 | 97 | 90.1 |
| 393 | VIR_2962 | Landrace | VIR 2962 | IT230190 | North Korea | 16,363,502,768 | 15.84 | 97.6 | 95.3 |
| 394 | VIR2980 | Landrace | VIR 2980 | IT228335 | North Korea | 17,080,689,952 | 15.73 | 98 | 95.1 |
| 395 | Well-man | Landrace | Well-man | PI 157487 A | South Korea | 29,001,403,676 | 28.03 | 98.2 | 96.9 |
| 396 | White-soybean | Landrace | White-soybean | PI 157488 | South Korea | 34,478,938,944 | 33.46 | 98.3 | 97 |
| 397 | Williams_82K | Improved | Williams 82K | A10 | United States | 18,817,443,724 | 18.92 | 99.7 | 98.8 |
| 398 | WIR2962 | Landrace | WIR2962 | IT199101 | North Korea | 17,727,785,956 | 17.37 | 98 | 95.9 |
| 399 | Wonhwang | Improved | Wonhwang | | South Korea | 16,001,611,336 | 14.53 | 97.6 | 94 |
| 400 | X1000A | Improved | X1000A | SS0404-T5-76 | South Korea | 17,630,955,394 | 16 | 97.5 | 91.6 |
| 401 | YB156 | Landrace | YB156 | IT024985 | South Korea | 18,589,822,608 | 18.02 | 98.3 | 96.5 |
| 402 | YB186-1 | Landrace | YB186-1 | IT025018 | South Korea | 20,608,913,370 | 19.96 | 98.4 | 96.8 |
| 403 | YB316-3 | Landrace | YB316-3 | IT024393 | South Korea | 15,622,093,070 | 14.99 | 96.4 | 89.5 |
| 404 | Yemsol | Landrace | Yemsol | | South Korea | 20,646,354,350 | 17.98 | 98.7 | 96.6 |
| 405 | YJ174-2 | Landrace | YJ174-2 | IT024070 | South Korea | 18,826,367,430 | 18.4 | 98.5 | 96.6 |
| 406 | YJ217-2 | Landrace | YJ217-2 | IT024106 | South Korea | 25,026,267,068 | 24.56 | 98.4 | 97.2 |
| 407 | YJ229-3 | Landrace | YJ229-3 | IT024128 | South Korea | 15,713,243,918 | 14.95 | 96.5 | 89.4 |
| 408 | YJ90 | Landrace | YJ90 | IT024019 | South Korea | 17,524,984,198 | 17.08 | 98.2 | 96.3 |
| 409 | YN154 | Landrace | YN154 | IT023887 | South Korea | 15,850,995,178 | 15.19 | 96.3 | 89.5 |
| 410 | YN161-3 | Landrace | YN161-3 | IT024572 | South Korea | 17,990,198,286 | 17.35 | 98.3 | 96.2 |
| 411 | YN188-3 | Landrace | YN188-3 | IT023925 | South Korea | 19,490,681,360 | 18.75 | 98.4 | 96.8 |
| 412 | YN197-1 | Landrace | YN197-1 | IT024431 | South Korea | 17,479,139,088 | 16.71 | 96.4 | 90.1 |
| 413 | YN213-2 | Landrace | YN213-2 | IT023953 | South Korea | 25,176,014,372 | 24.08 | 98.2 | 96.8 |
| 414 | Yonpoong | Improved | Yonpoong | | South Korea | 14,901,049,346 | 14.24 | 97.3 | 91.2 |
| 415 | York | Improved | York | PI 553038 | United States | 20,085,026,722 | 19.47 | 98.4 | 96.9 |
| 416 | You_huang_dou | Landrace | You huang dou | PI 567354 | China | 19,601,037,294 | 19.03 | 98.1 | 96.3 |

| | | | | | | | | | |
|---|---|---|---|---|---|---|---|---|---|
| 417 | Youkwoo16 | Improved | Yugwu16 | IT021836 | Japan | 16,996,661,472 | 16.17 | 97.2 | 90.4 |
| 418 | Yu_tae | Landrace | Yu tae | IT228397 | South Korea | 37,854,736,016 | 36.44 | 98.4 | 97.4 |
| 419 | CD-0250 | G. soja | YWS415 | | South Korea, Gangwon | 19,736,711,990 | 16.72 | 97.7 | 94.4 |
| 420 | CD-1078 | G. soja | B01042 | JPN1 | Japan, Hokkaido | 26,106,365,870 | 19.56 | 98.1 | 95.8 |
| 421 | CD-1094 | G. soja | B05051 | JPN42 | Japan, Mie | 32,017,601,960 | 16.51 | 98.2 | 94.5 |
| 422 | CD-1123 | G. soja | PI 483468 B | CHN17 | China, Henan | 29,015,820,754 | 19.31 | 97.8 | 94.1 |
| 423 | CD-1125 | G. soja | PI 464934 | 81-200001, CHN19 | China, Jiangsu | 30,128,143,046 | 16.02 | 98.1 | 94.1 |
| 424 | CD-1148 | G. soja | PI 597455 | ZYD3024, CHN53 | China, Shaanxi | 21,798,414,284 | 19.45 | 97.1 | 93.1 |
| 425 | CD-1156 | G. soja | PI 407300 | CHN61 | China, Zhejiang | 23,885,114,078 | 19.57 | 97.7 | 94.1 |
| 426 | Cheorwon | G. soja | Cheorwon | IT184216 | South Korea, Gangwon | 17,460,667,862 | 15.69 | 96.7 | 92.9 |
| 427 | CHN13 | G. soja | PI 464866 A | L 79-0009, CHN13 | China, Heilongjiang | 15,151,224,636 | 14.22 | 95.8 | 88.6 |
| 428 | CHN20 | G. soja | PI 464935 | 81-200002, CHN20 | China, Jiangsu | 16,816,557,128 | 16.09 | 97.1 | 92.6 |
| 429 | CHN22 | G. soja | PI 464936 B | 81-200004, CNH22 | China, Jiangsu | 17,775,156,166 | 17 | 96 | 91.1 |
| 430 | CHN23 | G. soja | PI 464937 B | 81-200014, CHN23 | China, Jiangsu | 19,704,373,842 | 19.34 | 96.9 | 93.8 |
| 431 | CHN25 | G. soja | PI 464939 B | 81-200027, CHN25 | China, Jiangsu | 17,113,318,938 | 16.23 | 95.7 | 90.9 |
| 432 | CHN28 | G. soja | PI 440913 B | CHN28 | China, Jilin | 19,314,062,700 | 18.92 | 97.5 | 95.1 |
| 433 | CHN3 | G. soja | PI 483462 A | CHN3 | China, Beijing | 17,367,592,670 | 16.46 | 95.6 | 90.3 |
| 434 | CHN30 | G. soja | PI 532452 B | GD50549, CHN30 | China, Jilin | 18,075,329,670 | 17.21 | 97.4 | 94.4 |
| 435 | CHN31 | G. soja | PI 464928 | LS-008, CHN31 | China, Liaoning | 16,555,038,416 | 16.42 | 97 | 93.9 |
| 436 | CHN34 | G. soja | PI 464927 C | LS-005, CHN34 | China, Liaoning | 17,210,371,168 | 15.14 | 97.2 | 92.9 |
| 437 | CHN35 | G. soja | PI 483460 C | CCHN35 | China, Liaoning | 19,240,074,512 | 16.93 | 96.3 | 90.5 |
| 438 | CHN37 | G. soja | PI 447003 B | CHN37 | China, Nei Monggol | 20,709,763,250 | 19.71 | 97.3 | 94.7 |
| 439 | CHN38 | G. soja | PI 468400 B | CHN38 | China, Ningxia | 16,175,223,888 | 15.91 | 97 | 93.8 |
| 440 | CHN40 | G. soja | PI 483464 B | CHN40 | China, Ningxia | 20,826,841,100 | 19.58 | 96.8 | 93.8 |

| | | | | | | | | | |
|---|---|---|---|---|---|---|---|---|---|
| 441 | CHN41 | G. soja | PI 483465 | CHN41 | China, Shaanxi | 16,984,494,798 | 16.24 | 97.2 | 92.9 |
| 442 | CHN42 | G. soja | PI 549046 | ZYDO 3728, CHN42 | China, Shaanxi | 16,856,410,256 | 16.05 | 95.7 | 89.2 |
| 443 | CHN43 | G. soja | PI 483466 | CHN43 | China, Shandong | 17,820,671,190 | 16.76 | 96.3 | 92.2 |
| 444 | CHN45 | G. soja | PI 597458 C | ZYD3233, CHN45 | China, Shandong | 16,528,907,262 | 15.55 | 95.5 | 88.1 |
| 445 | CHN46 | G. soja | PI 597459 D | ZYD3234, CHN46 | China, Shandong | 24,181,992,546 | 22.93 | 97.4 | 95.2 |
| 446 | CHN49 | G. soja | PI 407305 | CHN49 | China, Shanghai | 20,488,757,234 | 19.19 | 95.8 | 90.3 |
| 447 | CHN54 | G. soja | PI 597456 | ZYD3036, CHN54 | China, Shaanxi | 16,686,635,520 | 15.39 | 95.5 | 87.3 |
| 448 | CHN55 | G. soja | PI 597454 A | ZYD3015, CHN55 | China, Shanxi | 16,936,103,828 | 15.8 | 95.3 | 87 |
| 449 | CHN56 | G. soja | PI 597454 B | ZYD3015, CHN56 | China, Shaanxi | 20,632,189,416 | 19.83 | 97.4 | 94.9 |
| 450 | CHN58 | G. soja | PI 468396 A | CHN58 | China, Shaanxi | 17,020,720,302 | 16.11 | 95.2 | 88.5 |
| 451 | CHN60 | G. soja | PI 468398 C | CHN60 | China, Shaanxi | 16,894,268,070 | 16.13 | 95.4 | 88.5 |
| 452 | CHN64 | G. soja | PI 407303 | CHN64 | China, Zhejiang | 16,067,350,092 | 14.89 | 96.8 | 93.2 |
| 453 | CHN7 | G. soja | PI 522180 | ZYD 68, CHN7 | China, Heilongjiang | 17,492,311,422 | 16.64 | 96 | 90.2 |
| 454 | IT_182932 | G. soja | IT182932 | | South Korea, Gyeonggi | 24,261,941,644 | 24.67 | 97.6 | 95 |
| 455 | IT162825 | G. soja | IT162825 | B01 | South Korea, Gyeongbuk | 18,936,024,188 | 18.82 | 97.6 | 95.1 |
| 456 | IT178480 | G. soja | IT178480 | | South Korea, Chungbuk | 21,314,125,546 | 21.43 | 97.6 | 95 |
| 457 | IT182840 | G. soja | IT182840 | B04 | South Korea, Jeonbuk | 29,991,591,008 | 20.06 | 97.5 | 95.1 |
| 458 | IT182847 | G. soja | IT182847 | Jeonbuk Namwon sujib | South Korea, Jeonbuk | 16,184,737,190 | 15.27 | 97.5 | 94.1 |
| 459 | IT182848 | G. soja | IT182848 | B05 | South Korea, Jeonnam | 18,191,227,766 | 18.15 | 97.6 | 95.3 |
| 460 | IT182864 | G. soja | IT182864 | Gyeongnam Geochang sujib | South Korea, Gyeongnam | 17,107,382,826 | 16.2 | 97.6 | 94.7 |
| 461 | IT182869 | G. soja | IT182869 | B03 | South Korea, Gyeongnam | 18,681,027,670 | 18.74 | 97.6 | 95.1 |
| 462 | IT182919 | G. soja | IT182819 | Chungnam GongJu sujib | South Korea, Chungnam | 17,644,808,738 | 16.69 | 97.3 | 94.2 |
| 463 | IT182946 | G. soja | IT182946 | Gyeonggi Hwaseong sujib | South Korea, Gyeonggi | 18,776,377,068 | 17.25 | 97.3 | 93.4 |

| | | | | | | | | | |
|---|---|---|---|---|---|---|---|---|---|
| 464 | IT182968 | G. soja | IT183036 | Chungbuk Jecheon sujib | South Korea, Chungbuk | 17,193,895,558 | 16.89 | 97.4 | 93.6 |
| 465 | IT182976 | G. soja | IT182976 | Gyeonggi Yongin sujib | South Korea, Gyeonggi | 19,039,662,480 | 18.19 | 97.3 | 94.4 |
| 466 | IT182993 | G. soja | IT182993 | Kangwon Wonju sujib | South Korea, Gangwon | 16,723,989,296 | 15.77 | 97.5 | 94.3 |
| 467 | IT183036 | G. soja | IT183036 | Kangwon Yeongwol sujib | South Korea, Gangwon | 16,207,203,272 | 15.11 | 97.3 | 93.9 |
| 468 | IT183037 | G. soja | IT183037 | Chungbuk Jecheon sujib | South Korea, Chungbuk | 17,623,755,412 | 16.63 | 97.6 | 94.7 |
| 469 | IT183045 | G. soja | IT183045 | | South Korea, Gyeongbuk | 18,096,818,178 | 17.28 | 97.5 | 94.5 |
| 470 | IT183105 | G. soja | IT183105 | Chungnam Hongseong sujib | South Korea, Chungnam | 16,238,214,142 | 15.28 | 97.3 | 94 |
| 471 | IT183115 | G. soja | IT183115 | Chungnam Dangjin sujib | South Korea, Chungnam | 16,833,379,736 | 15.63 | 97.4 | 94.1 |
| 472 | IT183120 | G. soja | IT183120 | Gyeonggi Pyeongtaek sujib | South Korea, Gyeonggi | 18,699,009,802 | 17.67 | 97.6 | 94.7 |
| 473 | IT184053 | G. soja | Gyeonggi yongin sujib | IT184053 | South Korea, Gyeonggi | 18,551,789,936 | 17.7 | 97.5 | 94.6 |
| 474 | IT184108 | G. soja | IT184108 | Jeonnam Gangjin sujib | South Korea, Jeonnam | 16,658,403,352 | 15.92 | 97.4 | 94.1 |
| 475 | IT184146 | G. soja | IT184146 | Kangwon Chunseong sujib | South Korea, Gangwon | 19,831,811,500 | 18.88 | 97.5 | 94.8 |
| 476 | IT184194 | G. soja | IT184194 | Gyeonggi gwacheon sujib | South Korea, Gyeonggi | 16,075,579,592 | 14.95 | 97.3 | 93.7 |
| 477 | IT184233 | G. soja | IT184233 | | South Korea, Gyeonggi | 17,057,557,658 | 16.02 | 97.4 | 94.3 |
| 478 | IT184254 | G. soja | IT184254 | Chungbuk Yeongdong sujib | South Korea, Chungbuk | 16,395,467,052 | 15.58 | 97.5 | 94.2 |
| 479 | IT184258 | G. soja | IT184258 | Gyeongbuk Yecheon sujib | South Korea, Gyeongbuk | 16,459,779,160 | 15.6 | 97.3 | 93.8 |
| 480 | IT188367 | G. soja | IT188367 | Kangwon Yanggu sujib | South Korea, Gangwon | 16,340,722,606 | 14.99 | 97.4 | 94 |

| | | | | | | | | | |
|---|---|---|---|---|---|---|---|---|---|
| 481 | IT188404 | G. soja | IT188404 | Gyeongbuk Yecheon sujib | South Korea, Gyeongbuk | 17,066,819,394 | 15.74 | 97.4 | 94.1 |
| 482 | IT188418 | G. soja | IT188418 | Gyeongbuk Geumreung sujib | South Korea, Gyeongbuk | 17,113,509,500 | 16.24 | 97.4 | 94.2 |
| 483 | IT195544 | G. soja | IT195544 | Chungbuk Jincheon sujib | South Korea, Chungbuk | 16,386,985,684 | 15.24 | 97.3 | 93.7 |
| 484 | IT195562 | G. soja | IT195562 | Gwangju Gwangsan sujib | South Korea, Jeonnam | 16,576,531,756 | 15.53 | 97.6 | 94.2 |
| 485 | Jangheung | G. soja | Jangheung | IT195572 | South Korea, Jeonnam | 24,415,913,592 | 20.2 | 97.3 | 95.2 |
| 486 | JPN10 | G. soja | B01164 | JPN10 | Japan, Hokkaido | 16,897,581,916 | 16.8 | 97.3 | 94.6 |
| 487 | JPN12 | G. soja | B02086 | JPN12 | Japan, Aomori | 16,732,767,832 | 16.26 | 97.4 | 94.6 |
| 488 | JPN16 | G. soja | B02165 | JPN16 | Japan, Iwate | 18,201,011,802 | 17.86 | 97.3 | 94.8 |
| 489 | JPN18 | G. soja | B02206 | JPN18 | Japan, Akita | 17,735,030,332 | 16.77 | 97.6 | 94.9 |
| 490 | JPN23 | G. soja | B03025 | JPN23 | Japan, Toyama | 15,721,738,272 | 14.71 | 95.8 | 89.1 |
| 491 | JPN24 | G. soja | B03028 | JPN24 | Japan, Toyama | 22,054,834,104 | 21.04 | 97.8 | 95.9 |
| 492 | JPN29 | G. soja | B03046 | JPN29 | Japan, Fukui | 16,800,515,492 | 16.49 | 97.4 | 94.4 |
| 493 | JPN30 | G. soja | B03051 | JPN30 | Japan, Fukui | 15,935,431,660 | 15.11 | 97.2 | 94.1 |
| 494 | JPN31 | G. soja | B04054 | JPN31 | Japan, Nagano | 16,070,948,422 | 15.33 | 96.7 | 92.1 |
| 495 | JPN32 | G. soja | B04057 | JPN32 | Japan, Nagano | 21,664,023,756 | 19.83 | 97.6 | 95.5 |
| 496 | JPN36 | G. soja | B04118 | JPN36 | Japan, Chiba | 17,642,907,648 | 16.34 | 97 | 93.9 |
| 497 | JPN38 | Hybrid | B04134 | JPN38 | Japan, Yamanashi | 20,358,498,292 | 19.21 | 96.7 | 92 |
| 498 | JPN39 | G. soja | B04148 | JPN39 | Japan, Saitama | 16,808,094,786 | 15.79 | 97 | 93.6 |
| 499 | JPN40 | G. soja | B05047 | JPN40 | Japan, Gifu | 16,931,196,328 | 16.77 | 97.6 | 95 |
| 500 | JPN43 | G. soja | B05053 | JPN43 | Japan, Mie | 16,592,807,442 | 16.32 | 97.4 | 94.7 |
| 501 | JPN46 | G. soja | B06033 | JPN46 | Japan, Hyogo | 16,679,831,460 | 16.33 | 97.4 | 94.8 |
| 502 | JPN51 | G. soja | B06103 | JPN51 | Japan, Hyogo | 16,349,010,090 | 16.09 | 97.3 | 94.3 |
| 503 | JPN56 | G. soja | B07162 | JPN56 | Japan, Ehime | 22,654,215,316 | 21.74 | 97.6 | 95.6 |
| 504 | JPN65 | G. soja | B09014 | JPN65 | Japan, Hokkaido | 17,875,148,064 | 17.65 | 97.4 | 94.9 |
| 505 | JPN66 | G. soja | B09058 | JPN66 | Japan, Miyazaki | 17,717,711,236 | 17.37 | 97.4 | 94.7 |
| 506 | JPN70 | G. soja | B09089 | JPN70 | Japan, Oita | 15,218,043,948 | 14.49 | 96 | 89.2 |

| | | | | | | | | | |
|---|---|---|---|---|---|---|---|---|---|
| 507 | JPN71 | G. soja | B09103 | JPN71 | Japan, Fukuoka | 18,149,268,632 | 17.82 | 97.4 | 94.9 |
| 508 | JPN8 | G. soja | B01151 | JPN8 | Japan, Hokkaido | 17,465,543,652 | 16.2 | 95.8 | 87.6 |
| 509 | Mungyeong | G. soja | IT183060 | Gyeongbuk Mungyeong sujib | South Korea, Gyeongbuk | 16,027,306,100 | 15.15 | 96.5 | 92.2 |
| 510 | PI_339871B | G. soja | PI 339871 B | | South Korea, Jeju | 19,451,864,114 | 16.81 | 97.5 | 94.7 |
| 511 | PI_342620_A | G. soja | PI 342620 A | | Russia, Primorye | 20,071,320,452 | 19 | 97.7 | 95.3 |
| 512 | PI_378685 | G. soja | PI 378685 | | Japan, Ehime | 17,171,056,506 | 16.45 | 97.5 | 94.7 |
| 513 | PI_378691 | G. soja | PI 378691 | | Japan, Miyazaki | 26,609,187,098 | 25.79 | 97.8 | 95.9 |
| 514 | PI_424002 | G. soja | PI 424002 | | Russia, Amur | 21,359,683,010 | 17.7 | 97.4 | 94.1 |
| 515 | PI_458537A | G. soja | PI 458537 A | | China | 19,649,851,788 | 19.95 | 97.3 | 93.9 |
| 516 | PI_507822 | G. soja | PI 507822 | | Russia, Amur | 20,646,354,350 | 17.12 | 97.2 | 93.2 |
| 517 | PI_518282 | G. soja | PI 518282 | | Taiwan | 25,961,725,588 | 26.41 | 97.5 | 95.5 |
| 518 | PI-378689 | G. soja | PI 378689 | | Japan, Niigata | 16,827,444,530 | 15.99 | 96 | 89.8 |
| 519 | PI378698 | G. soja | PI 378698 | | Japan, Yamanashi | 16,822,944,126 | 16.13 | 96.7 | 93.2 |
| 520 | PI-507800-B | G. soja | PI 507800 B | | Russia, Primorye | 21,199,502,758 | 20.19 | 97.2 | 94.8 |
| 521 | PI-65549 | G. soja | PI 65549 | | China, Heilongjiang | 18,254,437,112 | 17.9 | 97.2 | 94.5 |
| 522 | YWS1001 | G. soja | YWS1001 | | South Korea, Jeonbuk | 20,455,551,730 | 18.99 | 96.3 | 90.6 |
| 523 | YWS1016 | G. soja | YWS1016 | | South Korea, Jeonbuk | 18,056,888,946 | 17.63 | 97.5 | 94 |
| 524 | YWS1022 | G. soja | YWS1022 | | South Korea, Jeonnam | 18,633,538,920 | 17.78 | 97.5 | 94.6 |
| 525 | YWS1023 | G. soja | YWS1023 | | South Korea, Jeonnam | 17,351,337,822 | 16.44 | 97.4 | 94.4 |
| 526 | YWS1026 | G. soja | YWS1026 | | South Korea, Jeonnam | 18,281,833,042 | 17.24 | 97.4 | 94.5 |
| 527 | YWS1027 | G. soja | YWS1027 | | South Korea, Jeonnam | 16,838,450,920 | 16.1 | 97.5 | 94.5 |
| 528 | YWS1029 | G. soja | YWS1029 | | South Korea, Jeonnam | 17,726,443,868 | 15.8 | 97.4 | 94.3 |
| 529 | YWS1031 | G. soja | YWS1031 | | South Korea, Jeonnam | 19,312,253,116 | 17.97 | 96.3 | 90.9 |
| 530 | YWS1034 | G. soja | YWS1034 | | South Korea, Jeonnam | 16,071,189,418 | 14.28 | 96.7 | 85.8 |
| 531 | YWS1041 | G. soja | YWS1041 | | South Korea, Gyeongbuk | 19,326,800,154 | 18.21 | 97.4 | 94.7 |
| 532 | YWS1055 | G. soja | YWS1055 | | South Korea, Gyeongbuk | 20,252,888,892 | 18.9 | 96.5 | 91.5 |
| 533 | YWS1064 | G. soja | YWS1064 | | South Korea, Gyeongbuk | 18,199,840,646 | 17.01 | 96 | 89.7 |
| 534 | YWS1089 | G. soja | YWS1089 | | South Korea, Chungnam | 16,265,257,940 | 15.26 | 96.8 | 92.9 |

| | | | | | | | | | |
|---|---|---|---|---|---|---|---|---|---|
| 535 | YWS1090 | G. soja | YWS1090 | | South Korea, Chungnam | 17,928,158,728 | 16.92 | 95.9 | 90.1 |
| 536 | YWS1098 | G. soja | YWS1098 | | South Korea, Chungnam | 16,978,423,088 | 15.74 | 97.3 | 93.9 |
| 537 | YWS1099 | G. soja | YWS1099 | | South Korea, Chungnam | 16,917,549,552 | 15.78 | 97.5 | 94.1 |
| 538 | YWS1100 | G. soja | YWS1100 | | South Korea, Chungnam | 16,771,534,968 | 15.88 | 97.2 | 94 |
| 539 | YWS1113 | G. soja | YWS1113 | | South Korea, Gangwon | 16,448,696,968 | 15.62 | 97.5 | 93.8 |
| 540 | YWS1114 | G. soja | YWS1114 | | South Korea, Gangwon | 16,774,328,468 | 15.89 | 97.3 | 93.9 |
| 541 | YWS1120 | G. soja | YWS1120 | | South Korea, Gangwon | 16,262,811,740 | 15.4 | 97.4 | 93.8 |
| 542 | YWS1123 | G. soja | YWS1123 | | South Korea, Gangwon | 18,389,996,154 | 17.44 | 97.4 | 94.7 |
| 543 | YWS1145 | G. soja | YWS1145 | | South Korea, Jeonnam | 17,887,992,124 | 16.75 | 96.3 | 90.6 |
| 544 | YWS1150 | G. soja | YWS1150 | | South Korea, Chungnam | 17,767,809,110 | 16.67 | 97.4 | 94.2 |
| 545 | YWS1155 | G. soja | YWS1155 | | South Korea, Chungnam | 17,930,401,682 | 16.67 | 97.4 | 94.4 |
| 546 | YWS1158 | G. soja | YWS1158 | | South Korea, Chungnam | 18,057,733,942 | 17.12 | 97.4 | 94.5 |
| 547 | YWS1160 | G. soja | YWS1160 | | South Korea, Chungnam | 17,127,112,788 | 15.88 | 97.5 | 94.2 |
| 548 | YWS1168 | G. soja | YWS1168 | | South Korea, Chungnam | 16,624,197,624 | 15.37 | 95.8 | 88.8 |
| 549 | YWS1175 | G. soja | YWS1175 | | South Korea, Chungnam | 16,404,932,638 | 15.63 | 97.5 | 94.2 |
| 550 | YWS1176 | G. soja | YWS1176 | | South Korea, Chungnam | 23,022,113,830 | 21.79 | 97.6 | 95.2 |
| 551 | YWS-1178-R | G. soja | YWS1178 | | South Korea, Chungnam | 20,633,908,800 | 16.41 | 97.2 | 94.6 |
| 552 | YWS1190 | G. soja | YWS1190 | | South Korea, Chungbuk | 16,436,229,502 | 15.53 | 97.2 | 94 |
| 553 | YWS1207 | G. soja | YWS1207 | | South Korea, Chungbuk | 18,092,416,830 | 17.15 | 97.4 | 94.4 |
| 554 | YWS121 | G. soja | YWS 121 | IT188382 | South Korea, Gangwon | 16,905,645,920 | 15.94 | 97.3 | 93.9 |
| 555 | YWS1214 | G. soja | YWS1214 | | South Korea, Chungbuk | 17,535,376,924 | 16.69 | 97.4 | 93.7 |
| 556 | YWS1216 | G. soja | YWS1216 | | South Korea, Chungbuk | 16,066,573,348 | 15.27 | 97.3 | 93.6 |
| 557 | YWS1222 | G. soja | YWS1222 | | South Korea, Gyeongnam | 16,610,563,834 | 15.72 | 97.5 | 94.4 |
| 558 | YWS1225 | G. soja | YWS1225 | | South Korea, Gyeongnam | 17,800,787,812 | 16.86 | 97.4 | 94.4 |
| 559 | YWS1228 | G. soja | YWS1228 | | South Korea, Gyeongnam | 21,148,507,642 | 19.36 | 96.5 | 91.4 |
| 560 | YWS1229 | G. soja | YWS1229 | | South Korea, Gyeongnam | 17,601,592,236 | 16.85 | 97.5 | 94.4 |
| 561 | YWS1230 | G. soja | YWS1230 | | South Korea, Gyeongnam | 17,475,403,348 | 16.06 | 97.5 | 94.5 |
| 562 | YWS1237 | G. soja | YWS1237 | | South Korea, Gyeongnam | 18,932,455,802 | 18.16 | 97.6 | 95 |
| 563 | YWS124 | G. soja | YWS 124 | IT184226 | South Korea, Gangwon | 16,663,489,938 | 15.61 | 96.9 | 93.6 |
| 564 | YWS1242 | G. soja | YWS1242 | | South Korea, Gyeongnam | 16,396,129,640 | 14.95 | 96.1 | 88.2 |

| | | | | | | | | | |
|---|---|---|---|---|---|---|---|---|---|
| 565 | YWS1243 | G. soja | YWS1243 | | South Korea, Busan | 16,522,663,412 | 15.41 | 97.3 | 93.8 |
| 566 | YWS1244 | G. soja | YWS1244 | | South Korea, Busan | 17,565,343,780 | 16.66 | 97.4 | 94.5 |
| 567 | YWS1246 | G. soja | YWS1246 | | South Korea, Ulsan | 16,278,295,884 | 15.18 | 97.2 | 93.5 |
| 568 | YWS1247 | G. soja | YWS1247 | | South Korea, Gyeongnam | 18,177,773,204 | 17.82 | 97.7 | 94.4 |
| 569 | YWS1263 | G. soja | YWS1263 | | South Korea, Chungnam | 20,470,929,872 | 18.05 | 96.7 | 91.3 |
| 570 | YWS1273 | G. soja | YWS1273 | | South Korea, Gyeonggi | 18,207,994,948 | 17.25 | 97.3 | 94.1 |
| 571 | YWS1274 | G. soja | YWS1274 | | South Korea, Gyeonggi | 17,418,026,066 | 16.48 | 97.4 | 94.5 |
| 572 | YWS1284 | G. soja | YWS1284 | | South Korea, Gyeonggi | 18,165,213,930 | 17.67 | 97.6 | 94.6 |
| 573 | YWS1285 | G. soja | YWS1285 | | South Korea, Gyeonggi | 16,622,707,858 | 15.87 | 96 | 89.7 |
| 574 | YWS1290 | G. soja | YWS1290 | | South Korea, Gyeonggi | 17,864,823,590 | 16.49 | 96.2 | 89.7 |
| 575 | YWS1309 | G. soja | YWS1309 | | South Korea, Gyeonggi | 16,761,237,372 | 15.38 | 95.8 | 88.8 |
| 576 | YWS1324 | G. soja | YWS1324 | | South Korea, Gyeongnam | 17,720,663,588 | 17.21 | 97.4 | 94.1 |
| 577 | YWS1328 | G. soja | YWS1328 | | South Korea, Gyeongnam | 17,193,957,468 | 16.08 | 96 | 89.6 |
| 578 | YWS-1338-R | G. soja | YWS1338 | | South Korea, Gyeongnam | 20,646,234,900 | 16.53 | 97.3 | 94.8 |
| 579 | YWS1354 | G. soja | YWS1354 | | South Korea, Gyeongbuk | 17,934,368,452 | 17.59 | 97.6 | 94.1 |
| 580 | YWS137 | G. soja | YWS 137 | IT182940 | South Korea, Gyeonggi | 17,338,506,446 | 16.28 | 97.5 | 94.5 |
| 581 | YWS1376 | G. soja | YWS1376 | | South Korea, Gyeongbuk | 18,969,708,408 | 17.84 | 97.4 | 94.6 |
| 582 | YWS139 | G. soja | YWS 139 | IT183026 | South Korea, Gyeonggi | 16,371,047,634 | 14.58 | 96 | 85.4 |
| 583 | YWS14 | G. soja | YWS 14 | | South Korea, Gyeongbuk | 15,795,893,164 | 14.54 | 96.7 | 92.5 |
| 584 | YWS1446 | G. soja | YWS1446 | | South Korea, Gyeongnam | 17,803,071,234 | 15.68 | 96.2 | 88.6 |
| 585 | YWS1447 | G. soja | YWS1447 | | South Korea, Gyeongnam | 17,478,348,150 | 16.38 | 97.5 | 94.4 |
| 586 | YWS1452 | G. soja | YWS1452 | | South Korea, Gyeongnam | 17,580,306,974 | 16.34 | 97.5 | 94.4 |
| 587 | YWS1457 | G. soja | YWS1457 | | South Korea, Gyeongbuk | 16,369,114,834 | 15.6 | 95.9 | 89.4 |
| 588 | YWS148 | G. soja | YWS 148 | | South Korea, Gyeongnam | 17,278,571,526 | 16.19 | 97.3 | 94.2 |
| 589 | YWS1502 | G. soja | YWS1502 | | South Korea, Gyeonggi | 15,550,412,464 | 14.81 | 96.6 | 91.9 |
| 590 | YWS1507 | G. soja | YWS1507 | | South Korea, Gyeongnam | 17,241,103,292 | 16.39 | 97.5 | 94.5 |
| 591 | YWS1512 | G. soja | YWS1512 | | South Korea, Gyeongnam | 17,111,146,954 | 16.09 | 97.5 | 94.4 |
| 592 | YWS1516 | G. soja | YWS1516 | | South Korea, Gyeongnam | 16,807,596,486 | 15.89 | 97.5 | 94.5 |
| 593 | YWS1518 | G. soja | YWS1518 | | South Korea, Gyeongnam | 17,817,194,868 | 16.54 | 97.4 | 94.4 |
| 594 | YWS1528 | G. soja | YWS1528 | | South Korea, Gyeongbuk | 18,088,876,484 | 16.96 | 97.5 | 94.6 |

| | | | | | | | | | |
|---|---|---|---|---|---|---|---|---|---|
| 595 | YWS1533 | G. soja | YWS1533 | | South Korea, Jeonnam | 18,007,500,470 | 17.01 | 97.5 | 94.8 |
| 596 | YWS1540 | G. soja | YWS1540 | | South Korea, Jeonnam | 17,786,873,464 | 16.52 | 97.6 | 94.7 |
| 597 | YWS1549 | G. soja | YWS1549 | | South Korea, Jeonnam | 18,183,092,330 | 17.67 | 97.6 | 94.6 |
| 598 | YWS1558 | G. soja | YWS1558 | | South Korea, Chungnam | 20,835,278,678 | 19.43 | 97.6 | 95.2 |
| 599 | YWS156 | G. soja | YWS 156 | | South Korea, Gyeongbuk | 17,281,487,638 | 16.31 | 97.3 | 94.1 |
| 600 | YWS1562 | G. soja | YWS1562 | | South Korea, Chungnam | 19,100,472,596 | 18.44 | 97.5 | 94.8 |
| 601 | YWS1577 | G. soja | YWS1577 | | South Korea, Jeonnam | 16,551,334,386 | 15.35 | 97.4 | 93.8 |
| 602 | YWS1580 | G. soja | YWS1580 | | South Korea, Jeonnam | 18,789,617,352 | 16.37 | 96.4 | 90.7 |
| 603 | YWS1582 | G. soja | YWS1582 | | South Korea, Jeonnam | 16,240,562,796 | 14.97 | 97.4 | 93.9 |
| 604 | YWS1584 | G. soja | YWS1584 | | South Korea, Jeonnam | 16,922,929,682 | 15.75 | 97.4 | 94 |
| 605 | YWS1588 | G. soja | YWS1588 | | South Korea, Jeonnam | 25,542,604,018 | 23.93 | 97.8 | 95.6 |
| 606 | YWS1592 | G. soja | YWS1592 | | South Korea, Jeonbuk | 17,484,633,978 | 16.54 | 97.4 | 94.4 |
| 607 | YWS1598 | G. soja | YWS1598 | | South Korea, Jeonbuk | 18,360,404,684 | 17.26 | 97.5 | 94.6 |
| 608 | YWS1605 | G. soja | YWS1605 | | South Korea, Jeonbuk | 17,261,463,226 | 16.2 | 97.5 | 94.6 |
| 609 | YWS1607 | G. soja | YWS1607 | | South Korea, Jeonbuk | 18,663,188,978 | 17.65 | 97.3 | 94.5 |
| 610 | YWS1608 | G. soja | YWS1608 | | South Korea, Jeonbuk | 17,268,286,614 | 15.92 | 97.4 | 93.7 |
| 611 | YWS1610 | G. soja | YWS1610 | | South Korea, Jeonbuk | 17,346,122,282 | 16.23 | 97.4 | 94 |
| 612 | YWS1611 | G. soja | YWS1611 | | South Korea, Jeonbuk | 17,888,862,186 | 17.01 | 97.5 | 94.7 |
| 613 | YWS1616 | G. soja | YWS1616 | | South Korea, Jeju | 17,018,748,242 | 16.12 | 97.6 | 94.5 |
| 614 | YWS1618 | G. soja | YWS1618 | | South Korea, Jeju | 18,443,981,372 | 17.55 | 97.5 | 93.9 |
| 615 | YWS1620 | G. soja | YWS1620 | | South Korea, Jeonnam | 17,353,514,034 | 16.41 | 97.3 | 94.1 |
| 616 | YWS1621 | G. soja | YWS1621 | | South Korea, Gwangju | 20,887,502,632 | 18.13 | 96.6 | 90.7 |
| 617 | YWS164 | G. soja | YWS 164 | IT188415 | South Korea, Gyeongbuk | 16,690,272,204 | 15.72 | 97.5 | 94.4 |
| 618 | YWS-17 | G. soja | YWS 17 | | South Korea, Gyeongbuk | 17,543,788,832 | 16.54 | 96.3 | 90.9 |
| 619 | YWS177 | G. soja | YWS 177 | IT178539 | South Korea, Chungnam | 19,229,258,080 | 18.35 | 97.5 | 94.9 |
| 620 | YWS192 | G. soja | YWS 192 | IT188399 | South Korea, Chungbuk | 16,091,762,262 | 14.84 | 97.3 | 93.8 |
| 621 | YWS193 | G. soja | YWS 193 | IT183098 | South Korea, Chungbuk | 17,731,375,528 | 16.74 | 97.5 | 94.5 |
| 622 | YWS199 | G. soja | YWS 199 | IT188396 | South Korea, Chungbuk | 18,262,747,548 | 17.24 | 97.5 | 94.2 |
| 623 | YWS204 | G. soja | YWS 204 | IT183052 | South Korea, Chungnam | 16,208,042,832 | 14.58 | 96 | 86.8 |
| 624 | YWS219 | G. soja | YWS219 | | South Korea, Jeonnam | 15,598,161,684 | 14.66 | 95.9 | 88.9 |

| | | | | | | | | |
|---|---|---|---|---|---|---|---|---|
| 625 | YWS229 | G. soja | YWS229 | South Korea, Jeonnam | 16,955,119,560 | 15.79 | 97.4 | 94.1 |
| 626 | YWS255 | G. soja | YWS255 | South Korea, Jeonnam | 16,160,344,650 | 15.09 | 97.4 | 93.9 |
| 627 | YWS26 | G. soja | YWS 26 | South Korea, Gyeongnam | 17,018,663,380 | 15.49 | 97.5 | 94.2 |
| 628 | YWS265 | G. soja | YWS265 | South Korea, Jeonbuk | 21,477,609,726 | 18.94 | 96.5 | 91.4 |
| 629 | YWS276 | G. soja | YWS276 | South Korea, Jeonbuk | 16,828,708,702 | 15.68 | 97.4 | 94.1 |
| 630 | YWS294 | G. soja | YWS294 | South Korea, Chungnam | 18,028,026,806 | 17.13 | 97.4 | 94.5 |
| 631 | YWS298 | G. soja | YWS298 | South Korea, Chungnam | 16,106,113,906 | 14.87 | 97.4 | 93.8 |
| 632 | YWS3 | G. soja | YWS 3 | South Korea, Gyeongbuk | 17,237,241,618 | 15.02 | 95.8 | 86.3 |
| 633 | YWS300 | G. soja | YWS300 | South Korea, Chungnam | 17,014,468,902 | 15.84 | 96.9 | 93.6 |
| 634 | YWS303 | G. soja | YWS303 | South Korea, Chungnam | 18,374,110,350 | 15.66 | 95.9 | 87.4 |
| 635 | YWS304 | G. soja | YWS304 | South Korea, Chungnam | 19,005,724,022 | 17.89 | 96.5 | 91.1 |
| 636 | YWS308 | G. soja | YWS308 | South Korea, Chungnam | 18,908,300,936 | 17.84 | 97.5 | 94.7 |
| 637 | YWS311 | G. soja | YWS311 | South Korea, Chungnam | 15,561,504,622 | 14.44 | 96.3 | 88.8 |
| 638 | YWS318 | G. soja | YWS318 | South Korea, Chungbuk | 18,399,533,012 | 17.27 | 97.7 | 94.4 |
| 639 | YWS331 | G. soja | YWS331 | South Korea, Chungbuk | 18,316,454,624 | 17.39 | 97.3 | 94.4 |
| 640 | YWS333 | G. soja | YWS333 | South Korea, Chungbuk | 22,035,802,366 | 20.66 | 96.4 | 91.8 |
| 641 | YWS34 | G. soja | YWS 34 | South Korea, Gyeongnam | 17,824,943,584 | 16.85 | 97.5 | 94.7 |
| 642 | YWS341 | G. soja | YWS341 | South Korea, Chungbuk | 17,321,073,496 | 16.45 | 97.4 | 94.3 |
| 643 | YWS347 | G. soja | YWS347 | South Korea, Chungbuk | 18,096,444,302 | 17.24 | 97.4 | 94.4 |
| 644 | YWS355 | G. soja | YWS355 | South Korea, Chungbuk | 16,706,787,980 | 15.51 | 97.4 | 94 |
| 645 | YWS361 | G. soja | YWS361 | South Korea, Chungbuk | 15,790,945,498 | 14.96 | 96 | 89.6 |
| 646 | YWS366 | G. soja | YWS366 | South Korea, Chungbuk | 17,450,634,516 | 16.59 | 97.4 | 94.5 |
| 647 | YWS368 | G. soja | YWS368 | South Korea, Chungbuk | 17,842,837,990 | 17.15 | 97.5 | 93.6 |
| 648 | YWS371 | G. soja | YWS371 | South Korea, Chungbuk | 16,153,542,704 | 15.25 | 97.3 | 93.9 |
| 649 | YWS39 | G. soja | YWS 39 | South Korea, Gyeongnam | 20,260,122,698 | 19.13 | 97.6 | 94.9 |
| 650 | YWS393 | G. soja | YWS393 | South Korea, Gyeonggi | 15,207,436,500 | 14.42 | 95.5 | 87.9 |
| 651 | YWS-4 | G. soja | YWS 4 | South Korea, Gyeongnam | 18,564,034,828 | 17.46 | 96.3 | 91.1 |
| 652 | YWS40 | G. soja | YWS 40 | South Korea, Gyeongbuk | 16,145,513,430 | 15.22 | 97.4 | 93.9 |
| 653 | YWS412 | G. soja | YWS412 | South Korea, Gangwon | 16,402,001,728 | 15.27 | 95.9 | 89.5 |
| 654 | YWS419 | G. soja | YWS419 | South Korea, Gangwon | 16,101,188,890 | 15.03 | 97.3 | 93.7 |

| | | | | | | | | |
|---|---|---|---|---|---|---|---|---|
| 655 | YWS421 | G. soja | YWS420A | South Korea, Gangwon | 16,089,507,228 | 14.97 | 95.9 | 88.6 |
| 656 | YWS426 | G. soja | YWS426 | South Korea, Gangwon | 16,936,013,228 | 15.83 | 97.1 | 93.7 |
| 657 | YWS427 | G. soja | YWS427 | South Korea, Gangwon | 18,345,999,888 | 17.28 | 97.4 | 94.4 |
| 658 | YWS432 | G. soja | YWS432 | South Korea, Gangwon | 16,362,685,858 | 15.41 | 97.3 | 94.1 |
| 659 | YWS433 | G. soja | YWS433 | South Korea, Gangwon | 18,100,696,764 | 17.22 | 97.4 | 94.6 |
| 660 | YWS435 | G. soja | YWS435 | South Korea, Gangwon | 18,669,383,300 | 18.27 | 97.4 | 94.4 |
| 661 | YWS438 | G. soja | YWS438 | South Korea, Gangwon | 17,147,832,102 | 16.05 | 97.5 | 94.4 |
| 662 | YWS448 | G. soja | YWS448 | South Korea, Gangwon | 16,091,246,748 | 15.02 | 97.4 | 93.7 |
| 663 | YWS449 | G. soja | YWS449 | South Korea, Gangwon | 18,015,798,524 | 17.09 | 97.4 | 94.4 |
| 664 | YWS45 | G. soja | YWS 45 | South Korea, Gyeongnam | 16,319,359,428 | 15.35 | 97.2 | 93.7 |
| 665 | YWS459 | G. soja | YWS459 | South Korea, Gangwon | 16,954,824,204 | 15.94 | 97.2 | 93.7 |
| 666 | YWS467 | G. soja | YWS467 | South Korea, Gangwon | 17,449,599,864 | 16.71 | 96.5 | 92.7 |
| 667 | YWS471 | G. soja | YWS471 | South Korea, Gangwon | 21,935,491,254 | 19.59 | 96.2 | 89.3 |
| 668 | YWS474 | G. soja | YWS474 | South Korea, Gangwon | 19,275,109,230 | 18.31 | 97.6 | 95 |
| 669 | YWS476 | G. soja | YWS476 | South Korea, Gangwon | 32,113,578,606 | 20.16 | 97.7 | 95.3 |
| 670 | YWS477 | G. soja | YWS477 | South Korea, Gangwon | 16,258,114,432 | 15.12 | 97.3 | 93.4 |
| 671 | YWS479 | G. soja | YWS479 | South Korea, Gangwon | 16,179,019,424 | 15.1 | 96.8 | 93.1 |
| 672 | YWS483 | G. soja | YWS483 | South Korea, Gangwon | 17,680,780,864 | 16.57 | 97.4 | 94.3 |
| 673 | YWS505 | G. soja | YWS505 | South Korea, Gangwon | 18,891,707,546 | 17.86 | 97.4 | 94.5 |
| 674 | YWS506 | G. soja | YWS506 | South Korea, Gangwon | 17,376,793,100 | 16.07 | 96.3 | 89.6 |
| 675 | YWS518 | G. soja | YWS518 | South Korea, Gangwon | 16,287,581,780 | 15.17 | 97.3 | 93.8 |
| 676 | YWS519 | G. soja | YWS519 | South Korea, Gyeonggi | 17,570,122,326 | 16.59 | 97.5 | 94.4 |
| 677 | YWS525 | G. soja | YWS525 | South Korea, Gyeonggi | 19,208,831,404 | 18.2 | 96.1 | 90.6 |
| 678 | YWS534 | G. soja | YWS534 | South Korea, Gyeonggi | 17,177,005,302 | 16.07 | 97.4 | 94.1 |
| 679 | YWS538 | G. soja | YWS538 | South Korea, Gyeonggi | 16,012,165,028 | 14.91 | 95.7 | 88.7 |
| 680 | YWS55 | G. soja | YWS 55 | South Korea, Gyeongbuk | 18,601,052,478 | 17.58 | 97.4 | 94.1 |
| 681 | YWS555 | G. soja | YWS555 | South Korea, Gyeonggi | 16,690,603,800 | 14.84 | 96.4 | 89.8 |
| 682 | YWS560 | G. soja | YWS560 | South Korea, Gyeonggi | 17,470,715,100 | 16.53 | 97.5 | 94.4 |
| 683 | YWS565 | G. soja | YWS565 | South Korea, Gyeonggi | 16,640,798,564 | 15.6 | 97.5 | 94.2 |
| 684 | YWS574 | G. soja | YWS574 | South Korea, Gyeonggi | 17,977,099,036 | 16.48 | 95.8 | 87.2 |

| | | | | | | | | |
|---|---|---|---|---|---|---|---|---|
| 685 | YWS580 | G. soja | YWS580 | South Korea, Gyeonggi | 17,104,391,818 | 15.87 | 97.5 | 94.3 |
| 686 | YWS583 | G. soja | YWS583 | South Korea, Gyeonggi | 16,930,528,002 | 15.79 | 96.1 | 89.4 |
| 687 | YWS586 | G. soja | YWS586 | South Korea, Gyeonggi | 18,094,908,330 | 17.25 | 97.4 | 94.4 |
| 688 | YWS59 | G. soja | YWS 59 | South Korea, Gyeongbuk | 17,058,472,416 | 15.9 | 97.6 | 94.3 |
| 689 | YWS6 | G. soja | YWS 6 | South Korea, Gyeongnam | 17,851,875,038 | 16.86 | 97.4 | 94.3 |
| 690 | YWS61 | G. soja | YWS 61 | South Korea, Gyeongbuk | 16,258,055,542 | 15.19 | 97.3 | 93.3 |
| 691 | YWS615 | G. soja | YWS615 | South Korea, Gyeonggi | 20,595,528,126 | 19.08 | 96.7 | 91.7 |
| 692 | YWS-63 | G. soja | YWS 63 | South Korea, Gyeongnam | 17,511,908,202 | 16.04 | 96.2 | 89.3 |
| 693 | YWS64 | G. soja | YWS 64 | South Korea, Gyeongnam | 17,320,872,364 | 16.47 | 97.4 | 94.4 |
| 694 | YWS645 | G. soja | YWS645 | South Korea, Jeju | 18,269,523,220 | 17.52 | 97.4 | 94.4 |
| 695 | YWS649 | G. soja | YWS649 | South Korea, Jeju | 17,097,948,648 | 16.22 | 97.4 | 94.3 |
| 696 | YWS65 | G. soja | YWS 65 | South Korea, Gyeongnam | 18,351,489,946 | 17.33 | 97.6 | 94.8 |
| 697 | YWS661 | G. soja | YWS661 | South Korea, Jeju | 16,886,749,176 | 16.1 | 97.5 | 94.4 |
| 698 | YWS662 | G. soja | YWS662 | South Korea, Jeju | 16,038,880,854 | 15.14 | 97.4 | 93.7 |
| 699 | YWS663 | G. soja | YWS663 | South Korea, Jeju | 16,484,295,218 | 15.46 | 96.2 | 89.8 |
| 700 | YWS669 | G. soja | YWS669 | South Korea, Jeju | 16,804,578,902 | 15.89 | 97.6 | 94.6 |
| 701 | YWS677 | G. soja | YWS677 | South Korea, Jeju | 16,851,100,190 | 15.93 | 95.9 | 89.8 |
| 702 | YWS678 | G. soja | YWS678 | South Korea, Jeju | 17,977,314,966 | 16.86 | 97.5 | 94.5 |
| 703 | YWS-68 | G. soja | YWS 68 | South Korea, Gyeongnam | 20,362,896,620 | 18.59 | 96.6 | 91.3 |
| 704 | YWS681 | G. soja | YWS681 | South Korea, Jeju | 16,756,279,438 | 15.57 | 97.4 | 94 |
| 705 | YWS689 | G. soja | YWS689 | South Korea, Chungbuk | 15,705,795,994 | 14.58 | 96.7 | 92.4 |
| 706 | YWS709 | G. soja | YWS709 | South Korea, Chungbuk | 16,982,926,814 | 16.27 | 96.5 | 92.3 |
| 707 | YWS712 | G. soja | YWS712 | South Korea, Chungbuk | 16,592,216,428 | 15.9 | 97.4 | 93.8 |
| 708 | YWS726 | G. soja | YWS726 | South Korea, Chungnam | 17,327,314,326 | 16.31 | 97.4 | 94.3 |
| 709 | YWS729 | G. soja | YWS729 | South Korea, Chungnam | 17,484,638,206 | 16.52 | 97.5 | 94.5 |
| 710 | YWS73 | G. soja | YWS 73 | South Korea, Gyeongnam | 18,453,124,120 | 17.46 | 97.6 | 94.8 |
| 711 | YWS738 | G. soja | YWS738 | South Korea, Chungnam | 17,343,507,868 | 16.37 | 97.4 | 94.2 |
| 712 | YWS742 | G. soja | YWS742 | South Korea, Chungnam | 16,259,113,750 | 15.34 | 96.9 | 93.2 |
| 713 | YWS75 | G. soja | YWS 75 | South Korea, Gyeongnam | 16,278,474,366 | 15.39 | 97 | 93.7 |
| 714 | YWS751 | G. soja | YWS751 | South Korea, Chungnam | 18,406,939,864 | 17.97 | 97.5 | 94.3 |

| | | | | | | | | | |
|---|---|---|---|---|---|---|---|---|---|
| 715 | YWS761 | G. soja | YWS761 | | South Korea, Chungnam | 16,669,185,356 | 15.48 | 97.4 | 94.2 |
| 716 | YWS765 | G. soja | YWS765 | | South Korea, Jeonbuk | 16,678,384,276 | 15.48 | 96.1 | 89.6 |
| 717 | YWS769 | G. soja | YWS769 | | South Korea, Jeonbuk | 16,856,888,926 | 15.58 | 97.4 | 94.1 |
| 718 | YWS785 | G. soja | YWS785 | | South Korea, Jeonbuk | 18,404,804,724 | 16.44 | 96.3 | 90.4 |
| 719 | YWS801 | G. soja | YWS801 | | South Korea, Jeonnam | 20,404,668,958 | 19.25 | 96.1 | 90.9 |
| 720 | YWS806 | G. soja | YWS806 | | South Korea, Jeonnam | 23,419,884,070 | 20.79 | 96.8 | 92.2 |
| 721 | YWS818 | G. soja | YWS818 | | South Korea, Jeonnam | 17,039,595,000 | 16.21 | 95.8 | 89.3 |
| 722 | YWS821 | G. soja | YWS821 | | South Korea, Jeonnam | 17,045,195,288 | 16.11 | 97.3 | 94.1 |
| 723 | YWS828 | G. soja | YWS828 | | South Korea, Gangwon | 18,224,995,736 | 17.23 | 97.4 | 94.6 |
| 724 | YWS837 | G. soja | YWS837 | | South Korea, Gangwon | 17,971,744,274 | 17.04 | 97.5 | 94.7 |
| 725 | YWS844 | G. soja | YWS844 | | South Korea, Gangwon | 17,288,145,832 | 16.31 | 96.6 | 92.8 |
| 726 | YWS847 | G. soja | YWS847 | | South Korea, Gangwon | 16,960,799,576 | 16.12 | 96.7 | 93.4 |
| 727 | YWS854 | G. soja | YWS853A | | South Korea, Gyeongbuk | 15,153,457,624 | 14.39 | 95.5 | 87.8 |
| 728 | YWS856 | G. soja | YWS856 | | South Korea, Gyeongbuk | 17,729,446,654 | 16.04 | 96 | 88.4 |
| 729 | YWS867 | G. soja | YWS846A | | South Korea, Gangwon | 17,181,917,030 | 16.26 | 95.9 | 88.9 |
| 730 | YWS870 | G. soja | YWS870 | | South Korea, Gyeongbuk | 16,288,878,568 | 15.32 | 97.4 | 94 |
| 731 | YWS871 | G. soja | YWS871 | | South Korea, Gyeongbuk | 17,647,531,268 | 16.65 | 95.9 | 89.7 |
| 732 | YWS876 | G. soja | YWS876 | | South Korea, Gyeongbuk | 18,333,129,554 | 17.57 | 97.5 | 94.8 |
| 733 | YWS878 | G. soja | YWS878 | | South Korea, Jeonnam | 17,384,790,966 | 16.58 | 97.5 | 94.4 |
| 734 | YWS88 | G. soja | YWS 88 | | South Korea, Gyeongnam | 16,939,157,048 | 15.75 | 97.4 | 94.1 |
| 735 | YWS885 | G. soja | YWS885 | | South Korea, Gyeongnam | 16,254,929,238 | 15.36 | 97.5 | 94.3 |
| 736 | YWS891 | G. soja | YWS891 | | South Korea, Gyeongnam | 16,516,635,190 | 15.59 | 97.3 | 94 |
| 737 | YWS900 | G. soja | YWS900 | | South Korea, Gyeongnam | 17,162,056,302 | 15.68 | 97.5 | 94.1 |
| 738 | YWS908 | G. soja | YWS908 | | South Korea, Gyeongnam | 16,667,978,564 | 15.88 | 97.5 | 94.1 |
| 739 | YWS909 | G. soja | YWS909 | | South Korea, Gyeongnam | 16,618,880,612 | 15.27 | 97.5 | 94.2 |
| 740 | YWS916 | G. soja | YWS916 | | South Korea, Gyeongnam | 17,921,753,610 | 17.04 | 97.5 | 94.5 |
| 741 | YWS923 | G. soja | YWS923 | | South Korea, Gyeonggi | 17,160,052,834 | 16.22 | 97.5 | 94.3 |
| 742 | YWS924 | G. soja | YWS926A | | South Korea, Gyeonggi | 18,082,082,994 | 17.23 | 96.4 | 91.4 |
| 743 | YWS932 | G. soja | YWS932 | | South Korea, Gyeonggi | 15,212,512,214 | 14.3 | 95.6 | 88.4 |
| 744 | YWS936 | G. soja | YWS936 | | South Korea, Chungbuk | 18,583,809,486 | 18.18 | 97.5 | 94.4 |

| | | | | | | | | | |
|---|---|---|---|---|---|---|---|---|---|
| 745 | YWS939 | G. soja | YWS939 | | South Korea, Gyeongbuk | 18,100,491,102 | 17.49 | 97.5 | 94.5 |
| 746 | YWS941 | G. soja | YWS941 | | South Korea, Gyeongbuk | 18,712,792,780 | 17.61 | 97.4 | 94.4 |
| 747 | YWS950 | G. soja | YWS950 | | South Korea, Jeonnam | 16,377,131,726 | 15.32 | 96.9 | 93.4 |
| 748 | YWS957 | G. soja | YWS957 | | South Korea, Jeonnam | 17,398,107,958 | 16.97 | 97.2 | 93.8 |
| 749 | YWS959 | G. soja | YWS959 | | South Korea, Jeonnam | 18,125,280,772 | 16.48 | 95.9 | 89.6 |
| 750 | YWS963 | G. soja | YWS963 | | South Korea, Jeonnam | 17,857,288,992 | 17.37 | 97.6 | 94 |
| 751 | YWS967 | G. soja | YWS967 | | South Korea, province unknown | 17,324,925,506 | 16.48 | 96 | 90.2 |
| 752 | YWS97 | G. soja | YWS 97 | | South Korea, Gyeongbuk | 17,805,431,062 | 16.52 | 97.5 | 94.4 |
| 753 | YWS971 | G. soja | YWS971 | | South Korea, Jeonbuk | 16,210,717,042 | 15.14 | 97.2 | 93.4 |
| 754 | YWS973 | G. soja | YWS973 | | South Korea, Jeonbuk | 16,303,611,034 | 15.2 | 97.4 | 94.1 |
| 755 | YWS975 | G. soja | YWS975 | | South Korea, Jeonbuk | 19,501,613,760 | 18.59 | 95.9 | 90.5 |
| 756 | YWS976 | G. soja | YWS976 | | South Korea, Jeonbuk | 16,733,383,006 | 15.83 | 97.4 | 94 |
| 757 | YWS978 | G. soja | YWS978 | | South Korea, Jeonbuk | 16,530,510,278 | 15.25 | 95.9 | 87.6 |
| 758 | YWS979 | G. soja | YWS979 | | South Korea, Jeonbuk | 19,824,744,700 | 18.4 | 96.3 | 90.8 |
| 759 | YWS982 | G. soja | YWS982 | | South Korea, Jeonbuk | 19,170,912,586 | 17.88 | 97.7 | 95 |
| 760 | YWS985 | G. soja | YWS985 | | South Korea, Jeonbuk | 17,331,258,748 | 16.16 | 97.5 | 94.4 |
| 761 | YWS988 | G. soja | YWS988 | | South Korea, Jeonbuk | 17,972,677,152 | 16.93 | 97.3 | 94 |
| 762 | YWS992 | G. soja | YWS992 | | South Korea, Jeonbuk | 17,425,578,784 | 16.32 | 97.5 | 94.3 |
| 763 | YWS996 | G. soja | YWS996 | | South Korea, Jeonbuk | 18,952,761,678 | 18.01 | 96 | 90.5 |
| 764 | 7402-18 | hybrid | 7402-18 | IT191214 | South Korea | 17,665,829,146 | 15.93 | 96.2 | 87.5 |
| 765 | 7402-29 | hybrid | 7402-29 | IT191225 | South Korea | 20,047,125,118 | 18.03 | 97 | 91.6 |
| 766 | 7402-46 | hybrid | 7402-46 | IT191242 | South Korea | 16,103,610,628 | 14.45 | 97.2 | 88.8 |
| 767 | 74052-5 | hybrid | 74052-5 | IT183431 | South Korea | 17,606,706,908 | 16.78 | 96.4 | 90.5 |
| 768 | CD-0073 | Hybrid | YWS 94 | | South Korea | 25,961,725,588 | 23.58 | 98.2 | 96.1 |
| 769 | CHN2 | Hybrid | PI 549048 | CHN2 | China | 19,141,877,098 | 17.65 | 96.6 | 90 |
| 770 | CHN57 | Hybrid | PI 597452 C | ZYD2900, CHN57 | China | 17,339,197,422 | 15.43 | 95.9 | 86.7 |
| 771 | YWS106 | Hybrid | YWS 106 | | China | 25,809,233,174 | 23.33 | 97 | 92.4 |
| 772 | YWS-125 | Hybrid | YWS 125 | | South Korea, province unknown | 15,557,172,432 | 14.48 | 96.3 | 88.9 |

| | | | | | | | | | |
|---|---|---|---|---|---|---|---|---|---|
| 773 | YWS1270 | Hybrid | YWS1270 | | South Korea | 23,469,454,350 | 18.96 | 96.2 | 88.1 |
| 774 | YWS136-1 | hybrid | YWS 136-1 | IT183014 | South Korea, Gyeonggi | 18,653,025,168 | 18.27 | 98.2 | 95.8 |
| 775 | YWS-16 | Hybrid | YWS 16 | | South Korea | 16,748,477,570 | 15.9 | 96.6 | 90.8 |
| 776 | YWS269 | Hybrid | YWS269 | | South Korea | 19,231,997,522 | 17.26 | 97.5 | 93.1 |
| 777 | YWS67 | Hybrid | YWS 67 | | South Korea | 16,538,489,420 | 14.73 | 96.3 | 86.9 |
| 778 | YWS74 | Hybrid | YWS 74 | | South Korea, Gyeongnam | 17,446,880,354 | 16.43 | 98.3 | 96.1 |
| 779 | YWS-74 | hybrid | YWS 74 | | South Korea | 18,768,903,776 | 17.01 | 97.1 | 92 |
| 780 | YWS740 | G. soja | YWS740 | | South Korea, Chungnam | 18,711,791,046 | 18.18 | 97.5 | 94.2 |
| 781 | YWS91 | hybrid | YWS 91 | | South Korea, Ulsan | 16,804,954,874 | 15.61 | 96.2 | 88 |
| Excluded | 74IT104887 | Landrace | IT104887 | | South Korea | 20,648,313,196 | 19.42 | 97.7 | 93.5 |
| Excluded | Baec-moc-sa-ryu | Landrace | Baec-moc-sa-ryu | PI 157398 | South Korea | 27,751,863,274 | 26.91 | 98.7 | 97.6 |
| Excluded | CS_02005 | Landrace | CS 02005 | IT229081 | South Korea | 31,679,765,270 | 30.76 | 98.8 | 97.9 |
| Excluded | IT104887 | Landrace | IT104887 | | South Korea | 15,264,186,226 | 14.71 | 96.9 | 90.4 |
| Excluded | KAS183-2 | Landrace | KAS183-2 | IT181404 | South Korea | 21,578,803,886 | 20.89 | 98.6 | 97.2 |
| Excluded | KAS351-20 | Landrace | KAS351-20 | PI 424352 | South Korea | 15,731,633,604 | 15.15 | 97.1 | 90.7 |
| Excluded | KAS351-20R | Landrace | KAS351-20 | PI 424352 | South Korea | 16,766,949,000 | 8.05 | 98.3 | 79.8 |
| Excluded | KLS101-2 | Landrace | KLS101-2 | PI 398866 | South Korea | 17,017,573,462 | 16.22 | 97.5 | 90 |
| Excluded | KLS101-2A | Landrace | KLS101-2 | PI 398866 | South Korea | 18,218,533,842 | 17.83 | 99 | 97.3 |
| Excluded | KLS-101-2-R_L1 | Landrace | KLS101-2 | PI 398866 | South Korea | 16,380,855,900 | 12.08 | 98.9 | 94.4 |
| Excluded | KLS86083 | Landrace | KLS86083 | IT142839 | South Korea | 18,018,900,064 | 17.33 | 97.6 | 91 |
| Excluded | PI_171434 | Landrace | PI 171434 | | China | 20,982,857,622 | 19.82 | 98.4 | 96.6 |
| Excluded | PI_458209 | Landrace | PI 458209 | | South Korea | 23,373,972,822 | 22.41 | 98.9 | 97.8 |
| Excluded | PI_86982 | Landrace | PI 86982 | | South Korea | 42,635,333,804 | 40.64 | 98.9 | 98.1 |
| Excluded | PI97150 | Landrace | PI 97150 | | North Korea | 29,393,920,324 | 28.25 | 99 | 98 |
| Excluded | Ryumuo | Landrace | Ryumuo | PI 88813 | North Korea | 21,513,873,584 | 20.89 | 98.7 | 97.4 |
| Excluded | VIR_2977 | Landrace | VIR 2977 | IT228332 | North Korea | 21,459,776,324 | 20.89 | 99.2 | 98.2 |
| Excluded | VIR_2978 | Landrace | VIR 2978 | IT228333 | North Korea | 15,949,872,696 | 15.12 | 98.5 | 96.3 |
| Excluded | WIR2987 | Landrace | WIR2987 | IT199107 | North Korea | 20,447,684,932 | 18.55 | 98.7 | 97.3 |

| | | | | | | | | | |
|---|---|---|---|---|---|---|---|---|---|
| Excluded | YJ87 | Landrace | YJ87 | IT024167 | South Korea | 24,316,250,270 | 23.35 | 98.7 | 97.7 |
| Excluded | YN213-1 | Landrace | YN213-1 | IT023952 | South Korea | 16,229,153,236 | 15.88 | 98.7 | 96.3 |
| Excluded | 19YWS1280 | G. soja | YWS1280 | | South Korea, Gyeonggi | 22,393,537,674 | 20.82 | 97.5 | 93.4 |
| Excluded | 48YWS1178 | G. soja | YWS1178 | | South Korea, Chungnam | 23,774,098,058 | 22.55 | 97.9 | 92.8 |
| Excluded | Buyeo | G. soja | Buyeo | IT182822 | South Korea, Chungnam | 17,312,813,796 | 16.4 | 97 | 93 |
| Excluded | IT188353 | G. soja | IT188353 | Gyeonggi Gapyeong sujib | South Korea, Gyeonggi | 17,607,552,206 | 16.59 | 98.5 | 95.5 |
| Excluded | Jecheon | G. soja | IT188385 | Chungbuk Jecheon sujib | South Korea, Chungbuk | 17,903,334,932 | 16.78 | 98 | 95 |
| Excluded | JPN21 | G. soja | B03016 | JPN21 | Japan, Niigata | 18,713,326,716 | 17.62 | 98 | 95.5 |
| Excluded | JPN62 | G. soja | B08050 | JPN62 | Japan, Shimane | 18,699,185,566 | 18.43 | 98.7 | 96.3 |
| Excluded | YWS1006 | G. soja | YWS1006 | | South Korea, Jeonbuk | 18,959,158,944 | 17.91 | 96.8 | 91.6 |
| Excluded | YWS-1006-R_L1 | G. soja | YWS1006 | | South Korea, Jeonbuk | 16,629,513,300 | 11.68 | 97.7 | 91.2 |
| Excluded | YWS1009 | G. soja | YWS1009 | | South Korea, Jeonbuk | 15,439,557,324 | 14.26 | 96.5 | 88.8 |
| Excluded | YWS-1009-R_L1 | G. soja | YWS1009 | | South Korea, Jeonbuk | 16,183,887,900 | 11.56 | 97.8 | 91.1 |
| Excluded | YWS1015 | G. soja | YWS1015 | | South Korea, Jeonbuk | 17,360,194,576 | 16.37 | 98.7 | 95.9 |
| Excluded | YWS1018 | G. soja | YWS1018 | | South Korea, Jeonbuk | 17,153,564,062 | 16.16 | 98.6 | 95.6 |
| Excluded | YWS-104887 | Landrace | IT104887 | | South Korea | 20,464,845,000 | 16.77 | 98.4 | 96.7 |
| Excluded | YWS1178 | G. soja | YWS1178 | | South Korea, Chungnam | 16,048,149,536 | 15.02 | 97.2 | 89.5 |
| Excluded | YWS1206 | G. soja | YWS1206 | | South Korea, Chungbuk | 16,098,493,842 | 15.04 | 97.3 | 90.3 |
| Excluded | YWS-1206-R | G. soja | YWS1206 | | South Korea, Chungbuk | 20,834,212,800 | 16.8 | 98.4 | 95.8 |
| Excluded | YWS1280 | G. soja | YWS1280 | | South Korea, Gyeonggi | 16,914,467,038 | 15.13 | 97 | 89.3 |
| Excluded | YWS-1280-R | G. soja | YWS1280 | | South Korea, Gyeonggi | 22,179,198,600 | 17.7 | 98 | 95.6 |
| Excluded | YWS1282 | G. soja | YWS1282 | | South Korea, Gyeonggi | 18,671,948,790 | 17.54 | 96.6 | 90.3 |
| Excluded | YWS-1282-R | G. soja | YWS1282 | | South Korea, Gyeonggi | 20,596,056,900 | 17.05 | 97.7 | 95.3 |
| Excluded | YWS1338 | G. soja | YWS1338 | | South Korea, Gyeongnam | 18,722,744,586 | 16.49 | 97.1 | 91.4 |
| Excluded | YWS1356 | G. soja | YWS1356 | | South Korea, Gyeongbuk | 19,085,154,250 | 18.76 | 98.2 | 95.2 |
| Excluded | YWS1462 | G. soja | YWS1462 | | South Korea, Gyeongbuk | 17,328,761,510 | 16.06 | 96.8 | 89.5 |
| Excluded | YWS1462R | G. soja | YWS1462 | | South Korea, Gyeongbuk | 32,452,332,300 | 26.62 | 98.2 | 96.7 |

| | | | | | | | | | |
|---|---|---|---|---|---|---|---|---|---|
| Excluded | YWS1513 | G. soja | YWS1513 | | South Korea, Gyeongnam | 18,293,646,678 | 17.61 | 98 | 95.2 |
| Excluded | YWS1541 | G. soja | YWS1541 | | South Korea, Jeonnam | 15,162,876,400 | 14.09 | 96.3 | 88.9 |
| Excluded | YWS1541R | G. soja | YWS1541 | | South Korea, Jeonnam | 30,036,280,200 | 25.06 | 98.1 | 96.3 |
| Excluded | YWS1553 | G. soja | YWS1553 | | South Korea, Chungnam | 17,055,428,860 | 16.35 | 98.2 | 94.8 |
| Excluded | YWS1601 | G. soja | YWS1601 | | South Korea, Jeonbuk | 16,179,823,650 | 15.3 | 98.1 | 94.7 |
| Excluded | YWS189 | G. soja | YWS 189 | IT184253 | South Korea, Chungbuk | 16,609,726,690 | 15.42 | 97.6 | 94.3 |
| Excluded | YWS263 | G. soja | YWS263 | | South Korea, Jeonbuk | 18,495,716,388 | 16.66 | 97.3 | 93.4 |
| Excluded | YWS330 | G. soja | YWS330 | | South Korea, Chungbuk | 20,521,868,212 | 19.16 | 98.2 | 95.8 |
| Excluded | YWS387 | G. soja | YWS387 | | South Korea, Chungbuk | 18,375,498,644 | 17.81 | 97.9 | 95.2 |
| Excluded | YWS405 | G. soja | YWS405 | | South Korea, Gyeongbuk | 16,419,070,466 | 15.22 | 97.8 | 94.3 |
| Excluded | YWS411 | G. soja | YWS411 | | South Korea, Gyeongnam | 16,370,701,542 | 15.38 | 98.1 | 94.6 |
| Excluded | YWS468 | G. soja | YWS468 | | South Korea, Gangwon | 17,427,697,918 | 16.55 | 98.1 | 95.1 |
| Excluded | YWS473 | G. soja | YWS473 | | South Korea, Gangwon | 19,357,540,130 | 18.05 | 98.3 | 95.7 |
| Excluded | YWS481 | G. soja | YWS481 | | South Korea, Gangwon | 16,453,871,738 | 15.13 | 97.2 | 87.7 |
| Excluded | YWS481R | G. soja | YWS481 | | South Korea, Gangwon | 29,789,853,600 | 24.1 | 98.8 | 97.2 |
| Excluded | YWS486 | G. soja | YWS486 | | South Korea, Gangwon | 17,056,326,102 | 15.94 | 96.7 | 89.6 |
| Excluded | YWS486R | G. soja | YWS486 | | South Korea, Gangwon | 24,964,923,300 | 21.51 | 97.9 | 95.9 |
| Excluded | YWS5 | G. soja | YWS 5 | | South Korea, Gyeongnam | 17,687,391,946 | 16.92 | 98.7 | 95.9 |
| Excluded | YWS511 | G. soja | YWS511 | | South Korea, Gangwon | 21,513,140,932 | 20.63 | 98.3 | 95.8 |
| Excluded | YWS633 | G. soja | YWS633 | | South Korea, Gyeonggi | 16,890,854,564 | 15.37 | 97.7 | 90.3 |
| Excluded | YWS-633-R | G. soja | YWS633 | | South Korea, Gyeonggi | 21,264,416,400 | 16.66 | 97.7 | 95 |
| Excluded | YWS66 | G. soja | YWS 66 | | South Korea, Gyeongnam | 18,709,326,726 | 17.84 | 98.2 | 95.5 |
| Excluded | YWS70 | G. soja | YWS 70 | | South Korea, Gyeongnam | 18,764,688,762 | 17.73 | 98.1 | 95.7 |
| Excluded | YWS83 | G. soja | YWS 83 | | South Korea, Gyeongnam | 15,623,942,820 | 14.83 | 97.5 | 94.1 |
| Excluded | YWS843 | G. soja | YWS843 | | South Korea, Gangwon | 16,702,332,876 | 15.51 | 98.2 | 94.8 |
| Excluded | YWS934 | G. soja | YWS934 | | South Korea, Chungnam | 17,612,203,006 | 16.6 | 98.1 | 95.2 |
| Excluded | YWS944 | G. soja | YWS944 | | South Korea, Gyeongbuk | 15,806,567,052 | 15.14 | 97.1 | 90 |
| Excluded | YWS944R | G. soja | YWS944 | | South Korea, Gyeongbuk | 26,448,974,700 | 22.57 | 98.9 | 97 |

**Supplementary Table 2. Whole-genome SNP, indel, structural variant distribution in 781 soybean genomes**

| Type | | Population | | | |
|---|---|---|---|---|---|
| | | Total | Glycine max | Glycine soja | Hybrid |
| Variants | SNPs | 10,597,683 | 6,085,061 | 10,323,813 | 6,134,427 |
| | Indels | 5,578,041 | 1,638,457 | 4,863,771 | 1,053,640 |
| | Structural variants | 139,011 | 35,156 | 116,039 | 17,017 |
| SNPs in genes | 5' UTR | 178,621 | 104,662 | 174,910 | 107,018 |
| | CDS | 397,869 | 230,796 | 386,815 | 239,617 |
| | 3' UTR | 196,040 | 114,478 | 191,977 | 119,930 |
| SIFT | Synomymous | 170,427 | 99,806 | 166,930 | 105,539 |
| | Non-synomymous tolerant | 149,077 | 89,916 | 145,017 | 92,679 |
| | Non-synomymous deleterious | 70,795 | 36,992 | 67,613 | 37,268 |
| | Stop-gain | 6,266 | 3,274 | 5,991 | 3,308 |
| | Stop-loss | 656 | 435 | 638 | 442 |

**Supplementary Table 3. Spearman rank's correlation coefficients between patterns of sequence variables (window size, 100 kb) observed across the genome.** All variables were significantly correlated at $P < 2.2e\text{-}16$

| | Gene | SNP | Indel | Rho | GERP | LD distance | SV |
|---|---|---|---|---|---|---|---|
| gene | | 0.294 | 0.725 | 0.133 | 0.925 | -0.786 | 0.234 |
| SNP | | | 0.591 | 0.297 | 0.314 | -0.424 | 0.255 |
| Indel | | | | 0.344 | 0.726 | -0.806 | 0.426 |
| Rho | | | | | 0.109 | -0.231 | 0.208 |
| GERP | | | | | | -0.800 | 0.200 |
| LD | | | | | | | -0.327 |

SNP, single nucleotide polymorphism; indel, insertion/deletion; Rho, historical recombination rate; GERP, genomic evolutionary rate profiling; LD, linkage disequilibrium; SV, structural variant (> 50 base pairs).

**Supplementary Table 4. List of genes that contain novel or standing domestication alleles[a]**

| ChrID | Position | Ref | Alt | Gene_name | Domestication allele frequency | | Mutation type | Function description |
|---|---|---|---|---|---|---|---|---|
| | | | | | Domest-icated | Wild | | |
| Chr01 | 26698265 | G | A | Glyma.01G088200 | 1.000 | 0.254 | Standing | NAC domain-containing protein 8-like |
| Chr02 | 8698038 | A | G | Glyma.02G096700 | 0.995 | 0.029 | Novel | histidine kinase 4-like isoform X2 |
| Chr02 | 38145634 | C | G | Glyma.02G199200 | 0.988 | 0.331 | Standing | Domain of unknown function DUF223 |
| Chr02 | 38257543 | G | A | Glyma.02G199400 | 1.000 | 0.458 | Standing | SERINE/THREONINE KINASE |
| Chr02 | 43679484 | A | G | Glyma.02G249200 | 1.000 | 0.006 | Novel | Unknown protein |
| Chr02 | 46816374 | T | C | Glyma.02G287800 | 0.983 | 0.445 | Standing | 50S/60S RIBOSOMAL PROTEIN L14/L23 |
| Chr02 | 46837711 | A | G | Glyma.02G288100 | 0.975 | 0.350 | Standing | Unknown protein |
| Chr03 | 29859032 | A | C | Glyma.03G104300 | 0.990 | 0.473 | Standing | 1-aminocyclopropane-1-carboxylate oxidase homolog 1 |
| Chr03 | 32149815 | C | G | Glyma.03G114400 | 0.990 | 0.038 | Novel | HEAT SHOCK PROTEIN 90 |
| Chr03 | 32149826 | G | A | Glyma.03G114400 | 0.990 | 0.065 | Standing | |
| Chr03 | 32150111 | C | A | Glyma.03G114400 | 0.998 | 0.327 | Standing | |
| Chr03 | 32150930 | G | T | Glyma.03G114400 | 0.990 | 0.035 | Novel | |
| Chr03 | 32151077 | G | C | Glyma.03G114400 | 0.993 | 0.297 | Standing | |
| Chr03 | 32151561 | A | G | Glyma.03G114400 | 0.990 | 0.033 | Novel | |
| Chr03 | 32151652 | C | A | Glyma.03G114400 | 0.993 | 0.298 | Standing | |
| Chr03 | 32151714 | T | C | Glyma.03G114400 | 0.990 | 0.036 | Novel | |
| Chr03 | 32330947 | C | A | Glyma.03G115500 | 0.971 | 0.276 | Standing | Phosphoglycerate mutase family protein |
| Chr03 | 32389132 | T | C | Glyma.03G115900 | 1.000 | 0.484 | Standing | GLUCOSYL/GLUCURONOSYL TRANSFERASES |
| Chr03 | 36544632 | G | A | Glyma.03G150300 | 0.998 | 0.221 | Standing | Kinesin motor domain |
| Chr03 | 37016008 | G | C | Glyma.03G154900 | 0.995 | 0.012 | Novel | Unknown protein |
| Chr03 | 37145505 | A | T | Glyma.03G156300 | 0.995 | 0.293 | Standing | Pentatricopeptide repeat (PPR) superfamily protein |
| Chr03 | 37388431 | C | T | Glyma.03G158600 | 0.957 | 0.134 | Standing | auxin response factor 11 |
| Chr03 | 40361746 | T | A | Glyma.03G192700 | 0.998 | 0.471 | Standing | ATP-binding ABC transporter |
| Chr03 | 40710113 | G | T | Glyma.03G197800 | 0.972 | 0.494 | Standing | uncharacterized protein |
| Chr03 | 42983644 | C | A | Glyma.03G227800 | 0.973 | 0.395 | Standing | transcription factor PIF1-like isoform X2 |
| Chr04 | 222549 | C | G | Glyma.04G002700 | 0.995 | 0.365 | Standing | Unknown protein |
| Chr04 | 767477 | A | G | Glyma.04G009600 | 0.995 | 0.348 | Standing | BEL1-like homeodomain protein 6-like isoform X3 |
| Chr04 | 866248 | C | T | Glyma.04G011200 | 0.955 | 0.251 | Standing | Protein of unknown function DUF241, plant |
| Chr04 | 873773 | G | A | Glyma.04G011300 | 0.954 | 0.106 | Standing | Protein of unknown function DUF241, plant |
| Chr04 | 3122316 | G | C | Glyma.04G039400 | 0.988 | 0.429 | Standing | THO complex, subunit 5; Fms-interacting protein |
| Chr04 | 48781472 | A | C | Glyma.04G216600 | 0.966 | 0.247 | Standing | endonuclease or glycosyl hydrolase |
| Chr04 | 48926556 | G | T | Glyma.04G218300 | 0.988 | 0.479 | Standing | ATP binding/protein serine/threonine kinase |

| | | | | | | | | |
|---|---|---|---|---|---|---|---|---|
| Chr04 | 49016112 | G | C | Glyma.04G219200 | 0.995 | 0.498 | Standing | shikimate kinase 1 |
| Chr05 | 4910515 | G | T | Glyma.05G054100 | 0.998 | 0.457 | Standing | MATE efflux family protein |
| Chr05 | 4911427 | T | C | Glyma.05G054100 | 0.990 | 0.239 | Standing | |
| Chr05 | 6893684 | T | A | Glyma.05G068100 | 1.000 | 0.482 | Standing | PLACENTAL PROTEIN 11-RELATED |
| Chr05 | 7237778 | G | A | Glyma.05G069500 | 1.000 | 0.491 | Standing | STEROL DESATURASE |
| Chr05 | 7328459 | G | A | Glyma.05G070100 | 1.000 | 0.432 | Standing | DNA helicase PIF1/RRM3 |
| Chr05 | 7328833 | C | T | Glyma.05G070100 | 1.000 | 0.435 | Standing | |
| Chr05 | 7330380 | G | A | Glyma.05G070100 | 1.000 | 0.435 | Standing | |
| Chr05 | 7330775 | G | T | Glyma.05G070100 | 1.000 | 0.437 | Standing | |
| Chr05 | 7330868 | G | T | Glyma.05G070100 | 1.000 | 0.449 | Standing | |
| Chr05 | 7332806 | C | T | Glyma.05G070100 | 1.000 | 0.432 | Standing | |
| Chr05 | 7332825 | T | G | Glyma.05G070100 | 1.000 | 0.438 | Standing | |
| Chr05 | 7332933 | C | T | Glyma.05G070100 | 1.000 | 0.429 | Standing | |
| Chr05 | 7677699 | C | A | Glyma.05G071100 | 1.000 | 0.361 | Standing | LEUCINE-RICH REPEAT RECEPTOR-LIKE PROTEIN KINASE |
| Chr05 | 7798430 | C | T | Glyma.05G071400 | 1.000 | 0.369 | Standing | F-box family protein |
| Chr05 | 8418504 | A | G | Glyma.05G073900 | 1.000 | 0.476 | Standing | Unknown protein |
| Chr05 | 9635497 | C | T | Glyma.05G076000 | 1.000 | 0.490 | Standing | pentatricopeptide (PPR) repeat-containing protein |
| Chr05 | 9951165 | G | C | Glyma.05G077000 | 1.000 | 0.194 | Standing | lysosomal beta glucosidase-like isoform X1 |
| Chr05 | 17038230 | A | G | Glyma.05G086800 | 1.000 | 0.053 | Standing | Unknown protein |
| Chr06 | 3710869 | T | A | Glyma.06G049300 | 0.986 | 0.142 | Standing | transcription factor bHLH74-like |
| Chr06 | 3736593 | T | A | Glyma.06G049500 | 0.986 | 0.402 | Standing | squalene monooxygenase 2 |
| Chr06 | 4677645 | G | C | Glyma.06G062200 | 0.956 | 0.111 | Standing | Unknown protein |
| Chr06 | 9585101 | A | G | Glyma.06G117700 | 0.998 | 0.496 | Standing | histone-lysine N-methyltransferase ASHH2-like isoform X1 |
| Chr06 | 11522690 | A | T | Glyma.06G141300 | 0.986 | 0.336 | Standing | centromere protein F-like isoform X3 |
| Chr07 | 13613966 | G | A | Glyma.07G119200 | 0.988 | 0.118 | Standing | Pentatricopeptide repeat (PPR) superfamily protein |
| Chr07 | 13760893 | T | C | Glyma.07G120000 | 0.980 | 0.312 | Standing | Unknown protein |
| Chr07 | 13762257 | A | C | Glyma.07G120000 | 0.974 | 0.300 | Standing | |
| Chr07 | 39487663 | C | A | Glyma.07G220300 | 0.988 | 0.314 | Standing | Cytochrome P450 superfamily protein |
| Chr07 | 39498121 | A | G | Glyma.07G220400 | 0.988 | 0.068 | Standing | Cytochrome P450 superfamily protein |
| Chr07 | 39611456 | A | T | Glyma.07G221000 | 0.990 | 0.050 | Novel | Membrane-associated apoptosis protein |
| Chr07 | 39611802 | A | T | Glyma.07G221000 | 0.988 | 0.084 | Standing | |
| Chr07 | 39612094 | G | A | Glyma.07G221000 | 0.990 | 0.085 | Standing | |
| Chr07 | 40138366 | T | C | Glyma.07G224100 | 0.978 | 0.383 | Standing | B3 DNA-binding domain protein |
| Chr07 | 41007164 | C | T | Glyma.07G229900 | 0.985 | 0.218 | Standing | Unknown protein |
| Chr07 | 41008638 | G | C | Glyma.07G229900 | 0.983 | 0.217 | Standing | |
| Chr08 | 10179 | G | A | Glyma.08G000100 | 1.000 | 0.492 | Standing | RNA-binding protein 24-like |
| Chr08 | 67241 | C | A | Glyma.08G000900 | 1.000 | 0.500 | Standing | uncharacterized protein LOC100305573 isoform X2 |
| Chr08 | 111715 | C | A | Glyma.08G001500 | 1.000 | 0.206 | Standing | COPINE FAMILY PROTEIN 2-RELATED |
| Chr08 | 957508 | C | A | Glyma.08G012000 | 0.986 | 0.264 | Standing | methyl-CpG-binding domain-containing protein 13-like isoform X2 |

| | | | | | | | | |
|---|---|---|---|---|---|---|---|---|
| Chr08 | 4056894 | A | T | Glyma.08G052200 | 0.988 | 0.265 | Standing | Unknown protein |
| Chr08 | 4387848 | T | C | Glyma.08G057200 | 0.988 | 0.066 | Standing | DUAL SPECIFICITY PROTEIN KINASE |
| Chr08 | 4402803 | C | A | Glyma.08G057300 | 0.993 | 0.193 | Standing | Domain of unknown function |
| Chr08 | 4486841 | G | C | Glyma.08G058600 | 0.993 | 0.391 | Standing | Calmodulin-binding protein |
| Chr08 | 4487186 | T | G | Glyma.08G058600 | 1.000 | 0.500 | Standing | |
| Chr08 | 6688253 | G | A | Glyma.08G088400 | 0.993 | 0.464 | Standing | cellulose synthase A4 |
| Chr08 | 6799055 | G | A | Glyma.08G090200 | 0.995 | 0.460 | Standing | UvrD-like Helicase, ATP-binding domain |
| Chr08 | 6839004 | T | A | Glyma.08G090900 | 0.995 | 0.020 | Novel | Unknown protein |
| Chr08 | 7074349 | T | A | Glyma.08G093300 | 0.977 | 0.192 | Standing | purple acid phosphatase 3 |
| Chr08 | 22117550 | T | C | Glyma.08G252100 | 0.988 | 0.063 | Standing | SERINE/THREONINE-PROTEIN KINASE PLK |
| Chr08 | 22637168 | T | C | Glyma.08G255400 | 0.997 | 0.349 | Standing | BETA-CAROTENE DIOXYGENASE |
| Chr09 | 2572792 | G | C | Glyma.09G031200 | 0.993 | 0.372 | Standing | evolutionarily conserved C-terminal region 7 (YTH domain) |
| Chr09 | 2572804 | T | A | Glyma.09G031200 | 0.993 | 0.006 | Novel | |
| Chr09 | 33308818 | G | A | Glyma.09G133900 | 0.993 | 0.264 | Standing | Deleted Pfam family |
| Chr09 | 33333060 | G | A | Glyma.09G134000 | 0.990 | 0.245 | Standing | MULTIDRUG RESISTANCE PROTEIN |
| Chr09 | 33333808 | G | C | Glyma.09G134000 | 0.990 | 0.260 | Standing | |
| Chr09 | 35793009 | G | C | Glyma.09G144100 | 0.993 | 0.098 | Standing | hAT family C-terminal dimerisation region |
| Chr10 | 45346535 | T | C | Glyma.10G222100 | 0.993 | 0.264 | Standing | one-helix protein 2 |
| Chr10 | 50741281 | T | A | Glyma.10G288500 | 1.000 | 0.411 | Standing | PROTEASE FAMILY S9A OLIGOPEPTIDASE |
| Chr10 | 51173313 | A | C | Glyma.10G294600 | 0.998 | 0.299 | Standing | Unknown protein |
| Chr11 | 10037575 | C | A | Glyma.11G131300 | 0.988 | 0.073 | Standing | receptor-like protein kinase 2 |
| Chr11 | 32199189 | G | A | Glyma.11G227100 | 0.993 | 0.327 | Standing | FRUCTOSE-1,6-BISPHOSPHATASE-RELATED |
| Chr11 | 32200152 | G | C | Glyma.11G227100 | 0.990 | 0.460 | Standing | |
| Chr11 | 32756745 | A | T | Glyma.11G232000 | 0.990 | 0.377 | Standing | mitotic checkpoint serine/threonine-protein kinase BUB1-like |
| Chr12 | 6728934 | A | T | Glyma.12G084200 | 1.000 | 0.360 | Standing | ADAPTIN, ALPHA/GAMMA/EPSILON; AP2 gene family |
| Chr12 | 6840846 | G | A | Glyma.12G084700 | 1.000 | 0.499 | Standing | histone deacetylase 2C |
| Chr12 | 19165287 | A | G | Glyma.12G144700 | 0.993 | 0.301 | Standing | Unknown protein |
| Chr12 | 19171416 | T | A | Glyma.12G144800 | 0.993 | 0.344 | Standing | receptor-like serine/threonine kinase 2 |
| Chr12 | 19190002 | G | A | Glyma.12G144900 | 0.993 | 0.334 | Standing | autoinhibited Ca2+ -ATPase, isoform 8 |
| Chr12 | 19190038 | T | C | Glyma.12G144900 | 0.993 | 0.335 | Standing | |
| Chr12 | 21441834 | G | T | Glyma.12G150000 | 1.000 | 0.457 | Standing | GAG/POL/ENV POLYPROTEIN |
| Chr12 | 21659261 | A | C | Glyma.12G150800 | 0.995 | 0.359 | Standing | choline transporter-like protein 2-like |
| Chr12 | 31859063 | G | C | Glyma.12G163900 | 0.990 | 0.176 | Standing | Unknown protein |
| Chr12 | 32039143 | G | A | Glyma.12G166000 | 0.998 | 0.206 | Standing | PIF1-like helicase |
| Chr12 | 32040407 | G | C | Glyma.12G166000 | 0.998 | 0.219 | Standing | |
| Chr12 | 33973586 | G | T | Glyma.12G179400 | 0.993 | 0.339 | Standing | SERINE PROTEASE FAMILY S10 SERINE CARBOXYPEPTIDASE |
| Chr12 | 36127088 | A | C | Glyma.12G199900 | 0.978 | 0.038 | Novel | anthocyanin 5-aromatic acyltransferase-like |
| Chr12 | 36132887 | C | G | Glyma.12G200000 | 1.000 | 0.370 | Standing | HXXXD-type acyl-transferase family protein |

| | | | | | | | | |
|---|---|---|---|---|---|---|---|---|
| Chr12 | 36221661 | T | G | Glyma.12G201700 | 0.971 | 0.282 | Standing | Folic acid binding / transferase |
| Chr13 | 22864538 | C | T | Glyma.13G114900 | 0.986 | 0.352 | Standing | conserved oligomeric Golgi complex subunit 3-like isoform X1 |
| Chr13 | 22868440 | G | T | Glyma.13G114900 | 0.985 | 0.350 | Standing | |
| Chr13 | 35231968 | A | C | Glyma.13G242900 | 0.995 | 0.416 | Standing | Toll-Interleukin-Resistance (TIR) domain family protein |
| Chr13 | 43088169 | T | C | Glyma.13G338100 | 0.995 | 0.218 | Standing | LACCASE-7-RELATED; Multicopper oxidase |
| Chr13 | 43088421 | T | C | Glyma.13G338100 | 0.995 | 0.287 | Standing | |
| Chr14 | 45068479 | G | T | Glyma.14G186300 | 1.000 | 0.389 | Standing | Pentatricopeptide repeat (PPR) superfamily protein |
| Chr14 | 45079666 | C | A | Glyma.14G186500 | 1.000 | 0.356 | Standing | E3 ubiquitin-protein ligase SHPRH-like isoform X2 |
| Chr14 | 45100710 | T | C | Glyma.14G186600 | 1.000 | 0.206 | Standing | ATP binding; GTP binding; nucleotide binding; nucleoside-triphosphatases |
| Chr14 | 45167254 | G | T | Glyma.14G187100 | 0.983 | 0.000 | Novel | Ribonuclease H-like superfamily protein |
| Chr15 | 11141248 | A | T | Glyma.15G137200 | 0.998 | 0.319 | Standing | GLYCOSYLTRANSFERASE 8 DOMAIN-CONTAINING PROTEIN |
| Chr15 | 11141559 | T | C | Glyma.15G137200 | 0.995 | 0.333 | Standing | |
| Chr15 | 11141560 | T | C | Glyma.15G137200 | 0.995 | 0.327 | Standing | |
| Chr15 | 11983924 | G | A | Glyma.15G145500 | 0.993 | 0.239 | Standing | Protein of unknown function |
| Chr15 | 12031428 | T | C | Glyma.15G146100 | 0.993 | 0.387 | Standing | disease-resistance response protein |
| Chr16 | 7134000 | G | A | Glyma.16G071100 | 0.966 | 0.145 | Standing | Ribonucleases P/MRP protein subunit POP1-like |
| Chr16 | 7134384 | G | A | Glyma.16G071100 | 0.966 | 0.200 | Standing | |
| Chr16 | 7151659 | A | G | Glyma.16G071400 | 0.964 | 0.304 | Standing | Unknown protein |
| Chr16 | 7174647 | A | G | Glyma.16G071500 | 0.961 | 0.044 | Novel | titin-like isoform X1 |
| Chr17 | 9164086 | T | C | Glyma.17G115900 | 0.998 | 0.003 | Novel | Lactoylglutathione lyase / glyoxalase I family protein |
| Chr17 | 10407797 | A | G | Glyma.17G129900 | 0.988 | 0.339 | Standing | Unknown protein |
| Chr17 | 10611472 | A | C | Glyma.17G132100 | 0.981 | 0.417 | Standing | Glucose-methanol-choline (GMC) oxidoreductase family protein |
| Chr19 | 38024288 | C | G | Glyma.19G122000 | 0.961 | 0.063 | Standing | Alpha-L-arabinofuranosidase C-terminus |
| Chr20 | 7649319 | G | A | Glyma.20G042500 | 0.995 | 0.251 | Standing | GLUTAMYL-TRNA(GLN) AMIDOTRANSFERASE SUBUNIT B |
| Chr20 | 7649623 | G | T | Glyma.20G042500 | 0.998 | 0.255 | Standing | |
| Chr20 | 8113895 | A | T | Glyma.20G044200 | 1.000 | 0.247 | Standing | rootletin-like isoform X3 |
| Chr20 | 8538821 | C | T | Glyma.20G045900 | 0.990 | 0.307 | Standing | Nucleoside diphosphate kinase |
| Chr20 | 8585160 | A | G | Glyma.20G046000 | 0.995 | 0.341 | Standing | Cytochrome P450 superfamily protein |
| Chr20 | 33631528 | A | G | Glyma.20G092900 | 0.986 | 0.197 | Standing | myosin-10 isoform X3 |
| Chr20 | 40460821 | A | G | Glyma.20G166700 | 0.998 | 0.404 | Standing | gene_model |

[a] Only deleterious alleles (nonsynonymous and stop mutations, SIFT score < 0.05 and GERP score > 0), which were detected in selective sweep regions, were considered. Domestication alleles that have allele frequency (AF) > 0.95 in domesticated soybean accessions and AF < 0.05 in wild soybean accessions were categorized as novel mutation type. Domestication alleles that have allele frequency (AF) > 0.95 in domesticated soybean accessions and 0.05 < AF < 0.50 in wild soybean accessions were categorized as standing mutation type.

**Supplementary Table 5. Comparison of position numbers of SNPs significantly associated with either protein or oil on chromosome 20 between Glyma1 (Ws82.a1.v1) and Ws82.a2.v1**

| SNP (Ws82.a1.v1) | SNP (Ws82.a2.v1) | | |
|---|---|---|---|
| Maker name with chromosome and position number | Marker name | Position on chromosome 20 | comment |
| BARC_1.01_Gm_20_29594697_A_G | ss715637225 | 30,744,968 | |
| BARC_1.01_Gm_20_29983050_A_G | ss715637240 | 31,129,249 | |
| BARC_1.01_Gm_20_30930931_A_G | ss715637283 | 21,083,902 | displaced |
| BARC_1.01_Gm_20_31150279_T_C | ss715637294 | 32,282,623 | |
| BARC_1.01_Gm_20_31243150_C_T | ss715637299 | 32,384,035 | |
| BARC_1.01_Gm_20_31436069_A_G | ss715637309 | 32,576,952 | |
| BARC_1.01_Gm_20_31580769_A_G | ss715637315 | 32,721,955 | |
| BARC_1.01_Gm_20_31610452_T_C | ss715637316 | 32,752,215 | |
| BARC_1.01_Gm_20_31640038_A_G | ss715637318 | 32,781,800 | |
| BARC_1.01_Gm_20_31687470_A_C | ss715637321 | 32,835,139 | |
| BARC_1.01_Gm_20_31972955_G_A | ss715637329 | 33,121,197 | |
